# Phage-mediated intercellular CRISPRi for biocomputation in bacterial consortia

**DOI:** 10.1101/2024.09.02.610857

**Authors:** Abhinav Pujar, Amit Pathania, Corbin Hopper, Amir Pandi, Cristian Ruiz Calderón, Matthias Függer, Thomas Nowak, Manish Kushwaha

**Affiliations:** Université Paris-Saclay, INRAE, AgroParisTech, Micalis Institute, 78352 Jouy-en-Josas, France; Université Paris-Saclay, CNRS, ENS Paris-Saclay, Laboratoire Méthodes Formelles, 91190 Gif-sur-Yvette, France; Université Paris-Saclay, CNRS, Laboratoire Interdisciplinaire des Sciences du Numérique, 91405 Orsay, France.trep; Institut Universitaire de France

## Abstract

Coordinated actions of cells in microbial communities and multicellular organisms enable them to perform complex tasks otherwise difficult for single cells. This has inspired biological engineers to build cellular consortia for larger circuits with improved functionalities, while implementing communication systems for coordination among cells. Here, we investigate the signalling dynamics of a phage-mediated synthetic DNA messaging system, and couple it with CRISPR interference to build distributed circuits that perform logic gate operations in multicellular bacterial consortia. We find that growth phases of both sender and receiver cells, as well as resource competition between them, shape communication outcomes. Leveraging the easy programmability of DNA messages, we build 8 orthogonal signals and demonstrate that intercellular CRISPRi (i-CRISPRi) regulates gene expression across cells. Finally, we multiplex the i-CRISPRi system to implement several multicellular logic gates that involve up to 7 cells and take up to 3 inputs simultaneously, with single- and dual-rail encoding: NOT, YES, AND, and AND-AND-NOT. The communication system developed here lays the groundwork for implementing complex biological circuits in engineered bacterial communities, using phage signals for communication.

## Introduction

Over the past two decades, synthetic biology has advanced the ability of engineered biological systems to sense environmental information and respond in a programmed manner. These capabilities have found applications in detecting pollutants (1) and disease biomarkers (2), smart therapeutics (3), and information processing logic circuits (4–6). This has been made possible by designing genetic circuits using molecular components such as biosensors, transcription factors, regulatory RNAs, riboswitches, or CRISPR systems for “internal wiring” (7, 8). While most engineered circuits are unicellular, the number of multicellular designs has been gradually increasing due to their several advantages (9–11), including reduced metabolic burden due to division of labour, minimised cross-talk, specialised sub-functions, distributed information processing, concurrency, redundancy, and fault tolerance (12). These properties contribute to the notion of “cellular supremacy” in biocomputing (13), whereby multicellular circuits are expected to enable more complex information processing (14).

However, multicellular circuits do introduce additional challenges, such as the need to balance populations of co-cultured cells that communicate through “external wiring”. To establish communication channels between cells in these circuits, several natural signalling molecules have been exploited; for example, homoserine lactones (HSL) from bacterial quorum sensing systems (15), yeast pheromones from mating signalling (10), and mammalian surface receptor-ligand pairs (11). Additionally, non-signalling molecules like secondary metabolites and synthetic coiled-coil peptides have been repurposed for signalling (16, 17). Despite these developments, the current repertoire of communication molecules—four HSLs (15), three pheromones (10, 18), two receptor-ligand pairs (11), six metabolites (16), and two coiled-coil ligands (17)—remains limited in both orthogonality and information capacity for higher-order multicellular circuits.

In nature, intercellular communication is not confined to small molecules but extends to information-rich nucleic acid molecules (DNA or RNA) transferred through cell junctions and vesicles (19), or mechanisms of horizontal gene transfer (20): transformation, conjugation, and transduction. These nucleic acid messages can confer new functions on receiver cells (21, 22), and can be rationally engineered to generate new orthogonal variants with altered specificities (23, 24). For instance, DNA delivery via conjugation has been used to modify undomesticated bacteria (25), target pathogenic strains (26), and selectively deliver DNA messages to specific cells within a population (27). Similarly, DNA delivery via bacteriophages has been utilised for phage therapy (28), inactivation of antimicrobial resistance (29), microbiome editing (30, 31), and metabolic pathway introduction (32). Despite their potential, phages are not commonly used for DNA propagation, possibly because most lyse their host cells upon release (33). Some exceptions are the non-lytic filamentous phages, which continuously secrete from infected cells, transmitting packaged DNA to new susceptible hosts (34). Of these, the M13 phage has been extensively studied (35), and is widely used for applications in nanotechnology (36), phage display (37), vaccine development (38), biosensing (39), and directed evolution (40, 41).

In fact, M13 was also used in the first demonstration of DNA-based communication for a multicellular circuit (42). This pioneering work from over a decade ago delineated a key property of nucleic acid signalling, “message-channel decoupling” (42), by which multiple message variants can be transmitted through a single communication channel. Despite this early advance, progress in M13-mediated DNA messaging has been limited. Possible challenges include its low adsorption rates (43), secretion heterogeneity (44), metabolic burden of infection (45), and superinfection immunity that prevents multiple phages from infecting the same cell (46).

In this study, we engineered two M13 phagemid variants (*-gp3φ* and *+gp3φ*), suited for different applications, and investigated how cell growth phase determines phage secretion and infection kinetics. We studied how phage particle concentration and receiver cell density impact infection rates in a well-mixed culture. These insights were used to implement a cell-to-cell communication system, quantifying its communication kinetics in a co-culture with resource competition. Next, we applied the phage communication system to engineer intercellular CRISPR interference (i-CRISPRi), where a single guide RNA (sgRNA) gene encoded on a phagemid is transmitted from a sender cell to a receiver cell, regulating gene expression across the extracellular space. We quantified the CRISPRi gene regulation kinetics for both isolated phages and sender cells. Finally, we demonstrated the multiplexing capability of the i-CRISPRi system by building multicellular logic gates with up to seven cells, which accept single (NOT, YES), double (AND), and triple (AND-AND-NOT) senders as input.

## Materials and Methods

### Bacterial strains, growth conditions, and cloning

All bacteria used in this study are *Escherichia coli* strains (**Table** S1). They were grown at 37 °C in LB media (liquid with shaking at 180 rpm, or solid LB plates with 1.5% w/v agar) supplemented with the appropriate antibiotics at the following concentrations (unless otherwise indicated): kanamycin (kan 30 µg mL^-1^), ampicillin (amp 100 µg mL^-1^), gentamycin (gent 10 µg mL^-1^), tetracycline (tet 10 µg mL^-1^), and spectinomycin (spc 50 µg mL^-1^); concentrations were halved when using multiple antibiotics for selection. Strains and antibiotics used are listed in **Table** S1. Core parental strains are listed in **Table** S3.

To calculate the growth rates of strains, cells were diluted 100x from overnight cultures and re-grown in a 96-well plate (200 µL per well) in a plate-reader (Biotek Synergy HTX) until they reached an OD_600_ of 0.2 to 0.3, following which they were diluted again by 40x into a new plate. Cultures in the second plate were grown overnight, recording their OD_600_ at 15 min intervals. The data generated were used to calculate the Specific Growth Rates (μ).

Cloning was performed by Golden Gate Assembly of PCR-amplified DNA fragments using NEB enzymes: Q5 DNA polymerase (#M0492), BsaI-HFv2 (#R3733), and T4 DNA ligase (#M0202M). *E. coli* strains DH5α and TOP10 were used for cloning. All plasmids constructed were verified by Sanger sequencing. Plasmids used in this study are listed in **Table** S2, and plasmid maps included with the Supplementary materials.

### Sender growth and phage preparation

Sender strains (**Table** S1) were streaked on LBA plates (1.5% w/v agar) and grown overnight at 37 °C. Single colonies were inoculated in 5 mL LB media with appropriate antibiotics and incubated overnight at 37 °C, with 180 rpm shaking. Overnight cultures were diluted 1000x in 100 mL fresh LB media with antibiotics and incubated for ∼15 hours at 37 °C, 180 rpm. Periodically, optical density was recorded (OD_600_; spectrophotometer UVisco V-1100D) and 1 mL culture sample was spun down at 4500x g for 10 mins, supernatant was filtered (0.22 µm filter, Millex SLGP033RS), and the resulting phage preps stored at 4 °C. Phage titres at each time-point were estimated using CFU or PFU assays (see below).

### Phage counting

#### CFU assay

Receiver cells (ER2738F) grown overnight were diluted 1000x and re-grown at 37 °C in LB (+tet) until they reached a spectrophotometer OD_600_ between 1 and 1.5. Cells were chilled on ice for at least 30 min, and then 90 µL aliquoted into eppendorf tubes. The tubes were moved to room temperature (RT) for 5 min before adding phages to the cells. 10 µL of different phage dilutions (10^-1^ to 10^-14^) were mixed with the receiver cells and incubated at RT for 20 min. Thereafter, the mixtures were plated on LBA plates with the appropriate antibiotic concentration. Colonies on the LBA plates were counted the next day after incubation at 37 °C for ∼16 h. Colony counts from plates were used to determine the mL^-1^ titres of the phage preps according to the formula: CFU count / (phage dilution * phage volume used in mL).

#### PFU assay

Receiver cells (ER2738F_HΔgIII) grown overnight were diluted 1000x and re-grown at 37 °C in LB (+tet+gent) until they reached a spectrophotometer OD_600_ between 1 and 1.5. Cells were chilled on ice for at least 30 min, and then 90 µL aliquoted into eppendorf tubes. The tubes were moved to RT shortly before mixing 10 µL of different phage dilutions (10^-1^ to 10^-14^) with the receiver cells, and then adding the mix to 10 mL of soft LBA (0.75% w/v agar with 0.2 mM IPTG and 40 µg mL^-1^ X-gal), previously aliquoted into a 15 mL tube and kept molten at 50 °C. The phage+receiver mix in the soft agar was immediately poured onto a solid plate with 20 mL hard LBA (1.5% w/v agar), and after the soft LBA had solidified the plate was incubated at 37 °C for 16-24 h. Plaques of the non-lytic M13 phage are turbid/ diffused, usually making them harder to see. IPTG and X-gal colours the plaques blue (LacZω in the F-plasmid is complemented by the LacZα in the phagemid), making them easier to visualise. Plaque counts from plates were used to determine the mL^-1^ titres of the phage preps according to the formula: PFU count / (phage dilution * phage volume used in mL).

### Instantaneous Growth and Secretion Rate analysis

Growth rates between two consecutive time-points were calculated according to the following formula: Specific Growth Rate (μ) = ln (OD_2_/OD_1_) / (t_2_-t_1_), where OD_1_ and OD_2_ are the OD_600_ values at time-points t_1_ and t_2_.

Secretion rates between two consecutive time-points were calculated according to the following formula: Secretion Rate = μ * (P_2_-P_1_) / (C_2_-C_1_), where P_1_ and P_2_ are the phage concentrations and C_1_ and C_2_ are the cell concentrations at time-points t_1_ and t_2_. μ is the specific growth rate calculated above. OD_600_ of sender cells was converted to cell concentration values using the fit in **Note** S2 (**Fig.** S2.1).

### Receiver infection analysis

To determine the effect of cell physiology on phage infection, receiver strain was streaked on LBA plates (1.5% w/v agar) and grown overnight at 37 °C. Single colony was inoculated in 5 mL LB media with appropriate antibiotics and incubated overnight at 37 °C, with 180 rpm shaking. Overnight cultures were diluted 1000x in 100 mL fresh LB media with antibiotics, and incubated at 37 °C, 180 rpm. OD_600_ of the culture was periodically monitored, and 10 mL culture samples were stored at 4 °C when they reached ODs: 0.05, 0.1, 0.5, 1, 1.5, 2, 2.5. Cultures were cooled on ice for at least 30 mins. Appropriate volumes of each culture sample was spun down at 4500x g for 10 mins to wash pellets with LB media without antibiotics and adjust (normalise) cell densities. These receiver culture samples were used for CFU and PFU assays with the same phage concentrations.

To determine the effect of cell density on phage infection, receiver strain was streaked on LBA plates (1.5% w/v agar) and grown overnight at 37 °C. Single colony was inoculated in 5 mL LB media with appropriate antibiotics and incubated overnight at 37 °C, 180 rpm. Overnight cultures were diluted 1000x in 25 mL fresh LB media with antibiotics and incubated at 37 °C, 180 rpm, till the OD_600_ reached ∼1.5. Culture was cooled on ice for at least 30 mins and then spun down at 4500x g for 10 mins to wash pellets with LB media without antibiotics. Culture was adjusted to different densities 0.05, 0.1, 0.5, 1, 2, 3, 4 and 5. These receiver cultures were used for CFU and PFU assays with the same phage concentrations.

### Receiver infection in growing conditions and quantifying unadsorbed phages

For the infection plate-reader experiments, overnight cultures of receiver cells (ER2738F) were diluted 1000x and re-grown to a spectrophotometer OD_600_ of 0.6-0.7, following which they were cooled on ice for ∼30 min and their OD_600_ re-adjusted to different densities (0.25, 0.125, 0.0625. 0.03125) while still on ice. Several serial dilutions (3^N^-fold, N = 0 to 10) of the phage prep (pSB1K3_M13ps_LacZα_gIII, undiluted concentration of 36.2x10^5^ PFU mL^-1^), and a no-phage control, were prepared and 45 µL aliquoted into a 96-well plate at RT. 150 µL of the different receiver dilutions were added to the plate and incubated at RT for 20 min. Next, 5 µL of LB was added to each well, without or with kanamycin (end concentration 30 µg mL^-1^), and the plate incubated overnight in a plate-reader (Biotek Synergy HTX) at 37 °C, 205 cpm, while recording OD_600_ at 15 min intervals. The above experiments were repeated four times, each with a different set of phage dilutions added to the plate: (a) 3x technical replicates of dilutions 3^8^-3^10^ and the no-phage control (continuous), (b) 3x technical replicates of dilutions 3^1^-3^3^ and the no-phage control (continuous), (c) 3x technical replicates of dilutions 3^1^-3^3^ and the no-phage control (discontinuous), and (d) single replicate of dilutions 3^0^-3^10^ and the no-phage control (continuous).

In the discontinuous run above, the plate was paused at several time-points (2, 6, and 10 h) to draw a 3 µL sample from each well of the third column (phage dilution 3^2^), which was added to 200 µL of LB (+gent) to kill all cells and later used to quantify by PFU assay the unadsorbed phages in the well. The pauses for phage sampling resulted in an average gap of ∼45 min between plate reader measurements before and after the pause, which was taken into account for plotting OD vs time curves.

### Phage mediated sender-to-receiver communication in co-cultures

For the communication plate-reader experiments, an overnight culture of receiver cells (ER2738F) was diluted 1000x and re-grown to a spectrophotometer OD_600_ of 0.4, following which it was cooled on ice for ∼30 min, pellets were washed, and several OD_600_ dilutions made (0.136, 0.068, 0.034, and 0.0) while still on ice. Overnight culture of senders (TOP10_H_gIII-KanΦ and TOP10_H_-KanΦ) was diluted 500x and re-grown to a spectrophotometer OD_600_ of 0.2, following which it was cooled on ice for ∼30 min, pellets were washed, and several OD_600_ dilutions made (0.125, 0.062, 0.031, 0.015, 0.007, and 0.0) while still on ice. 90 µL of receiver cell dilutions were added per well to a 96-well plate in quadruplet (for the four different growth conditions), followed by 90 µL of the sender cell dilutions also in quadruplet. The plate was run at 37 °C for 1 h, following which 20 µL of LB with the appropriate antibiotics (10x concentrated, to achieve the 1x end-concentration) was added to each well, and the plate was grown overnight at 37 °C, 205 cpm, while recording OD_600_ at 15 min intervals.

### Receiver infection CRISPRi time lapse in growing conditions

Receiver strain (carrying F-plasmid + dCas9-GFP plasmid) and sender strains (carrying helper plasmid + sgRNA phagemid) were streaked on LBA plates (1.5% w/v agar), with appropriate antibiotics, and grown overnight at 37 °C. Single colonies were inoculated in 5 mL LB media with appropriate antibiotics and incubated overnight at 37 °C, with 180 rpm shaking. Overnight cultures were diluted 1000x in 100 mL fresh LB media with antibiotics and incubated at 37 °C, 180 rpm, until they reached OD_600_ ∼0.3. Cultures were cooled on ice for at least 30 mins, pellets were washed, and several dilutions made with different ODs (0.12, 0.06, 0.03, and 0.0), while still on ice. The supernatant was filtered through 0.22 µm filters to collect phages and several serial dilutions were made (10x, 20x, 40x, 0). 100 µL of receiver cultures were mixed with 100 µL of sender cultures in a 96-well plate and grown in plate reader for 5 hours with no antibiotic, while recording OD_600_ and GFP fluorescence at 15 min intervals. At 0, 1, 2, 3, 4 and 5 hours, 10 µL co-culture was added to 1x PBS containing 2 mg mL^-1^ kanamycin to stop protein expression and kill the cells. Following this, the cells in 1x PBS + kan (2 mg mL^-1^) were stored at 4 °C for later flow cytometry analysis.

### Phage transfer frequency calculations

Transfer frequencies were calculated from the co-culturing flow cytometry data in **Fig.** 4d-e and S6.5. The events recorded at each time-point were gated by fluorescence to obtain the number of cells of each type: senders (S, no GFP), uninfected receivers (Ru, GFP ON) and infected receivers (Ri, GFP OFF). The transfer frequency was calculated as Ri/S*(Ri+Ru), according to the formula used to obtain transconjugant frequency in conjugation experiments (27).

### Single input CRISPRi biological circuits

Receiver strain (carrying F-plasmid + dCas9-GFP plasmid) and sender strains (carrying helper plasmid + sgRNA phagemid) were streaked on LBA plates (1.5% w/v agar), with appropriate antibiotics, and grown overnight at 37 °C. Single colonies were inoculated in 5 mL LB media with appropriate antibiotics and incubated overnight at 37 °C, with 180 rpm shaking. Overnight cultures were diluted 1000x in 100 mL fresh LB media with antibiotics and incubated at 37 °C, 180 rpm, until they reached OD_600_ ∼0.3. Cultures were cooled on ice for at least 30 mins, spun down and washed. Several dilutions made to different ODs (0.12, 0.06, 0.03, 0.01, 0.007 and 0.0), while still on ice. 90 µL of receiver cell dilutions were added per well to a 96-well plate in quadruplet (for the four different growth conditions), followed by 90 µL of the sender cell dilutions also in quadruplet. The plate was run at 37 °C for 4 h, while recording OD_600_ and GFP fluorescence at 15 min intervals, following which 20 µL of LB with the appropriate antibiotics (10x concentrated, to achieve the 1x end-concentration) was added to each well, and the plate was grown overnight at 37 °C, 205 cpm, while recording OD_600_ and GFP fluorescence at 15 min intervals. After 16 hours of growth in selection media, 10 µL of cultures were transferred to PBS for flow cytometry analysis performed immediately afterwards.

### Multi-input CRISPRi biological circuits

Receiver strain (carrying F-plasmid + dCas9-GFP plasmid) and sender strains (carrying helper plasmid + sgRNA phagemid) were streaked on LBA plates (1.5% w/v agar), with appropriate antibiotics, and grown overnight at 37 °C. Single colonies were inoculated in 5 mL LB media with appropriate antibiotics and incubated overnight at 37 °C, with 180 rpm shaking. Overnight cultures were diluted 1000x in 100 mL fresh LB media with antibiotics and incubated at 37 °C, 180 rpm, until they reached OD_600_ ∼0.3. Cultures were cooled on ice for at least 30 mins, spun down and washed. Several dilutions made to different ODs (0.12, 0.06, 0.03, and 0.0), while still on ice. 100 µL of receiver cell dilutions were added per well to a 96-well plate, followed by 100 µL (total) of the sender cell dilutions. For the 2-input gate, 50 µL each of the 2 sender dilutions was mixed together before adding to the receivers. For the 3-input gate, 33.3 µL each of the 3 sender dilutions was mixed together before adding to the receivers. The plate was run at 37 °C for 4 h, while recording OD_600_ and GFP fluorescence at 15 min intervals, following which the co-culture was diluted 20x in 200 µL fresh LB media with antibiotics, selecting only infected receivers. The plate was grown overnight while recording OD_600_ and GFP fluorescence at 15 min intervals. After 16 hours of growth in selection media, cultures were used to prepare PBS+kan plates for flow cytometer analysis performed immediately afterwards.

### Flow cytometry analysis

Flow cytometry samples containing cells in 1x PBS (with 2 mg mL^-1^ of kanamycin) were analysed using the Attune NxT flow cytometer (Thermofisher) equipped with a Cytkick autosampler. Samples were fed into the flow cytometer using 96-well plates. ∼20, 000 bacterial events were recorded per sample, excluding dead cells and debris from the analysis using FSC and SSC thresholds of 100. GFP fluorescence was measured using excitation by a 488 nm laser and a 530/30 nm filter (BL1 channel). The BL1 fluorescence threshold for each ON/OFF circuit was defined as the lower extreme of the fluorescence distribution from the ON receiver population. In samples with a mix of sender and receiver cells, the autofluorescent sender population was gated as BL1<100 before gating the ON/OFF receiver cells. Voltages used were FSC: 265, SSC: 273, BL1: 278, for all experiments. Data collected were analysed using Attune Cytometric v5.3, and plotted using python scripts. Flow cytometry overlays were plotted using the online tool: floreada.io

## Results

### Sender physiology impacts phage secretion rates

Many previous studies have examined the infection and secretion kinetics of the wild-type M13 phage (35). M13 is a filamentous bacteriophage that infects F+ *Escherichia coli* cells, using the host cell’s F-pilus as its primary receptor. Once inside the host, the single-stranded phage DNA (ssDNA) rapidly converts into a double-stranded replicative form (RF) DNA, which is then replicated by the host machinery. The 6.4 kb phage genome encodes for 11 phage proteins (gp1-11), including the structural coat proteins (gp3, gp6–9), and those involved in DNA synthesis, packaging, and secretion (gp1–2, gp4–5, gp10–11) (**Fig.** 1a) (35, 45). Of these, the gp3 protein is responsible for superinfection immunity that prevents the same cell from being infected multiple times (46). In addition, a packaging signal (ps) is required for mobilisation of the phage DNA for packaging into a new phage particle (47, 48). In several previous works, M13 phage components have been engineered to package other plasmid DNA by adding a packaging signal (48); or, to make phage secretion dependent on the conditional expression of an essential phage protein like: gp3 (40, 49), gp6 (41), or gp8 (50).

**Fig. 1:**
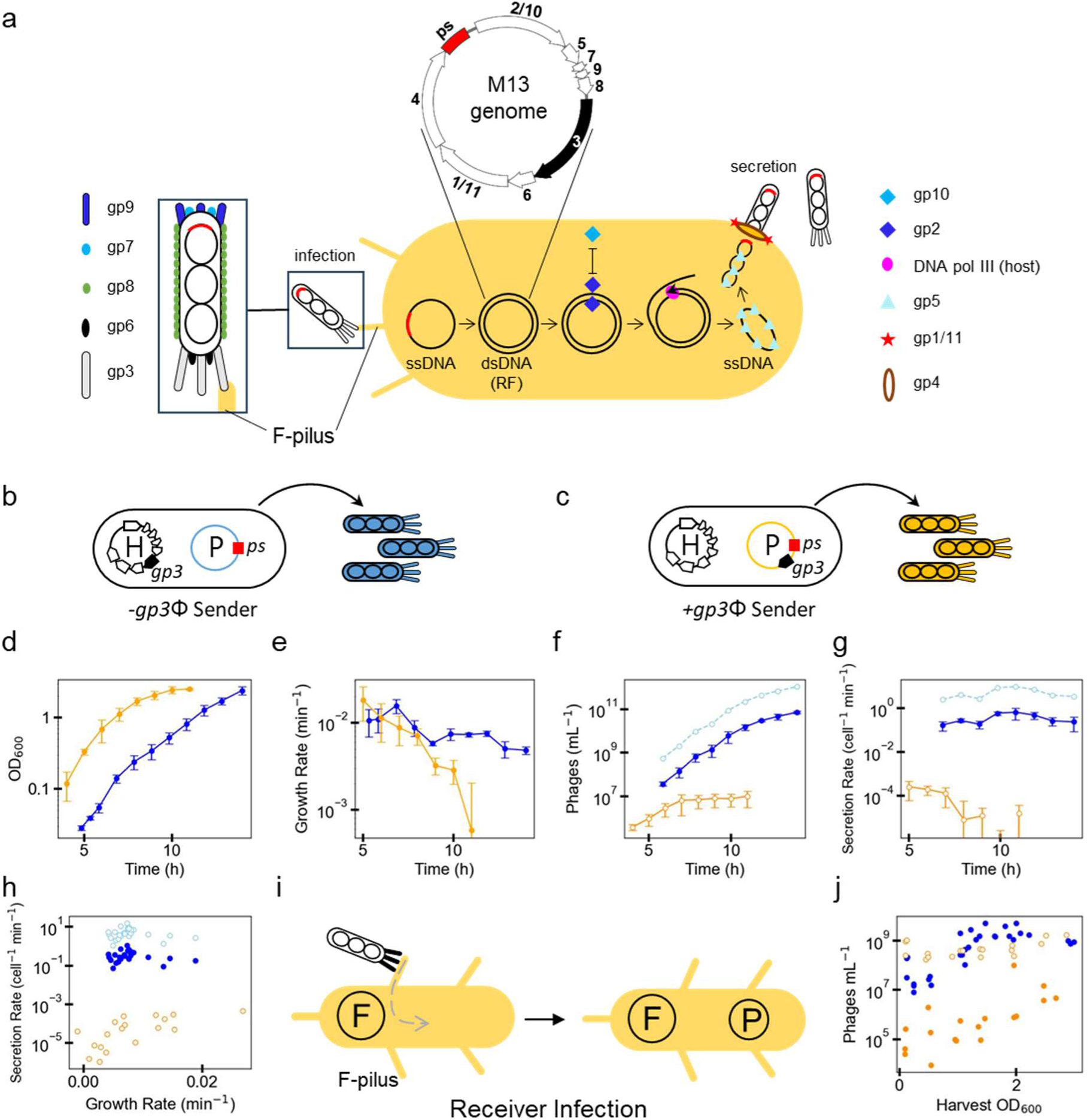
Secretion and infection kinetics of M13 phages are growth-phase dependent. **(a)** Schematic representation of the wild-type M13 bacteriophage’s life cycle in *E. coli*. The M13 phage, with single-stranded DNA (ssDNA) packaged in multiple coat proteins (gp3, gp6–9), enters the host cell following attachment to the F-pilus receptor on the bacterial cell surface. The ssDNA delivered is rapidly converted to double-stranded replicative form DNA (dsDNA, RF). While the host machinery produces multiple copies of the dsDNA, the ssDNA is regenerated and assembled into phage particles by the phage protein machinery (gp1–2, gp4–5, gp10–11). **(b-c)** Schematics of phage-secreting sender cell variants: *(b) -gp3φ* (TOP10_H_KanΦ) and (c) *+gp3φ* (TOP10_HΔgIII_gIII-KanΦ) sender. Both variants carry an M13 helper plasmid (H) that encodes the phage machinery and a phagemid (P), a plasmid that carries a packaging signal (ps) for secretion. The essential minor coat protein gene, *gp3*, is encoded on the helper in the *-gp3φ* sender and on the phagemid in the *+gp3φ* sender. **(d)** Growth curves of *-gp3φ* senders (blue) and *+gp3φ* senders (orange), plotted as OD_600_ against time. **(e)** Instantaneous growth rates of the two senders plotted against time, calculated between each pair of consecutive time-points from the growth data in (d). **(f)** Phage secretion curves of *-gp3φ* and *+gp3φ* senders plotted as phage titres against time. Titres of phages obtained from the sender time-points in (d) were estimated using a CFU assay (*-gp3φ*, filled circles) or a PFU assay (*+gp3φ*, empty circles). (f-h) Re-calculated PFU estimates for the *-gp3φ* phage titres are shown as empty light-blue circles. **(g)** Instantaneous secretion rates were calculated for each consecutive time-point pair from secretion curves obtained in (f). Data in (d)-(g) show mean±SD from N=3 repeats. **(h)** Instantaneous secretion rates from (g) plotted against instantaneous growth rates from (e). **(i)** Schematic of a phage particle infecting a receiver cell. The receiver carries an F-plasmid encoding the F-pilus, the primary receptor for M13 phage infection. **(j)** Receiver cells (ER2738F) harvested at different growth phases (harvest OD_600_) were re-adjusted to the same density, and infected with the same number of isolated phage particles (*-gp3φ* or *+gp3φ*). The number of infected cells were counted using a CFU assay (filled circles) or a PFU assay (empty circles). Data in (j) show individual points from N=3 repeats.

In this study, we investigate the secretion kinetics of M13 phages produced by engineered sender cells. Similar to prior works (42), our senders contain two plasmids: a helper (H) encoding the functional M13 proteins (gp1-gp11) and a phagemid (P) carrying the M13 packaging signal for packaging and secretion. For some applications, a phage without re-secretion suffices (28), while others require amplification through re-secretion (40). We constructed two sender variants to compare: -gp3φ for delivery only, and +gp3φ for delivery and re-secretion. Both variants have a *kanR* gene on the phagemid and constitutively secrete phage particles. They differ in whether the minor coat protein *gp3* gene is encoded on the helper or on the phagemid (**Fig.** 1b and 1c).

To study the kinetics of secretion, we monitored sender cultures for ∼15 hours, measuring OD_600_ and collecting phage samples at 1 h intervals (**Fig.** 1d and 1f). -gp3φ and +gp3φ phages were quantified using an assay for colony-forming units (CFU) or plaque-forming units (PFU), respectively (**Fig.** 1f). -gp3φ phages cannot repackage inside receiver cells due to the absence of the essential *gp3* gene, while +gp3φ phages can re-secrete if receivers have the complementary phage machinery. Consequently, -gp3φ phages are quantified using CFU assay whereas +gp3φ phages can be quantified using both CFU and PFU assays (see **Note** S1 for differences between CFU and PFU assays). CFU counts from the -gp3φ phages were converted to PFU estimates (**Fig.** 1f-h), using the experimentally determined multiplication factor of 14.7 (**Fig.** S1.3). This allows for direct comparisons with the -gp3φ phage counts from PFU assays, which provide more accurate measures of phage numbers (**Note** S1).

Although both sender variants reached similar end-point ODs, -gp3φ senders had a longer lag phase compared to +gp3φ senders (**Fig.** 1d). -gp3φ senders showed a higher instantaneous growth rate (**Fig.** 1e), and produced 5 orders of magnitude more phages at the end-point than +gp3φ senders (**Fig.** 1f). The per cell secretion rate of -gp3φ phages remained relatively constant, while that of +gp3φ phages decreased over time, with maximum secretion rates of 9.7 and 0.0002 phages min^-1^ cell^-1^, respectively (**Fig.** 1g; see **Note** S2 for the relationship between OD and sender cell numbers). The secretion rates of -gp3φ senders are comparable to the 2–6 phages min^-1^ cell^-1^ reported for wild-type filamentous phages (35, 50), but those for +gp3φ are much lower. Differences in secretion rates probably stem from differences in expression levels of the *gp3* gene (50) from the different plasmids: helper plasmid for -gp3φ (pBBR1, copy number 4.7 (51)) and phagemid for +gp3φ (pUC, copy number 8.9 (51)). Consistent with prior findings (52), we observe a positive correlation between growth rates and secretion rates per cell (**Fig.** 1h), suggesting that higher growth rates are required to support phage secretion. Both phages show a plateau in secretion rate at a growth rate of 0.0073 min^-1^ (0.44 h^-1^), indicating that other bottlenecks limit secretion beyond this growth rate. Overall, we find that secretion kinetics of M13 phages are specific to each phagemid, and depend on sender cell growth phase and phage machinery expression levels.

### Receiver physiology impacts phage infection rates

Having established that the growth phase significantly affects sender cell secretion rates, next we investigated its impact on receiver cell infection rates. Whereas in a growing batch culture, cell density and growth phase vary simultaneously, we designed our experiments to separate the two effects (**Fig.** 1j, S1.1-S1.2). Receiver cells were harvested at different ODs, and resuspended to the same OD of 0.5, a commonly used density for PFU and CFU assays. These resuspended cells were then infected with identical phage concentrations and counted using a CFU assay for both -gp3φ and +gp3φ phages, and a PFU assay for +gp3φ phages (**Fig.** 1i–j). Phage CFU counts in the late growth phase were 217-fold higher for -gp3φ and 323-fold higher for +gp3φ phages than in the early growth phase, despite the same receiver cell OD. This suggests that receiver cells at different phases of growth have different infectability, possibly due to more phage receptors (F-pili) per cell in the late-log to early stationary phase (53). However, this effect is not apparent when using PFU assays (**Fig.** 1j, **Note** S1).

To assess the effect of receiver density on infection, cells were grown to an OD ∼1, harvested, and resuspended to five different ODs before infection with the same phage concentrations using a CFU assay (**Fig.** S1.2). Consistent with the law of mass action, higher receiver densities resulted in more infected cells (CFU counts). Together, these findings emphasise the impact of receiver cell physiology and density on infection rates, as well as the limitations of the CFU method in accurately quantifying phage numbers (**Note** S1).

### The role of stochastic interactions in infection dynamics of growing receiver cells

After examining the effects of physiology and density on receiver infection independently (**Fig.** 1), we investigated how phage infection impacts growing receiver cells in batch cultures where both factors change simultaneously. We used the +gp3φ phage that can infect receiver cells only once (**Fig.** S3.1), due to superinfection immunity from gp3 expression in receivers (46). Receiver cells at four starting densities (OD_600_: 0.25, 0.125, 0.0625, 0.03125) were incubated with 12 concentrations of isolated phages (fold-dilutions: 3^0 to 3^10, and no-phage control) for 20 minutes in a 96-well plate (**Fig.** 2a), followed by growth with or without antibiotic selection (kanamycin) for ∼20 hours (**Fig.** S4.1).

**Fig. 2:**
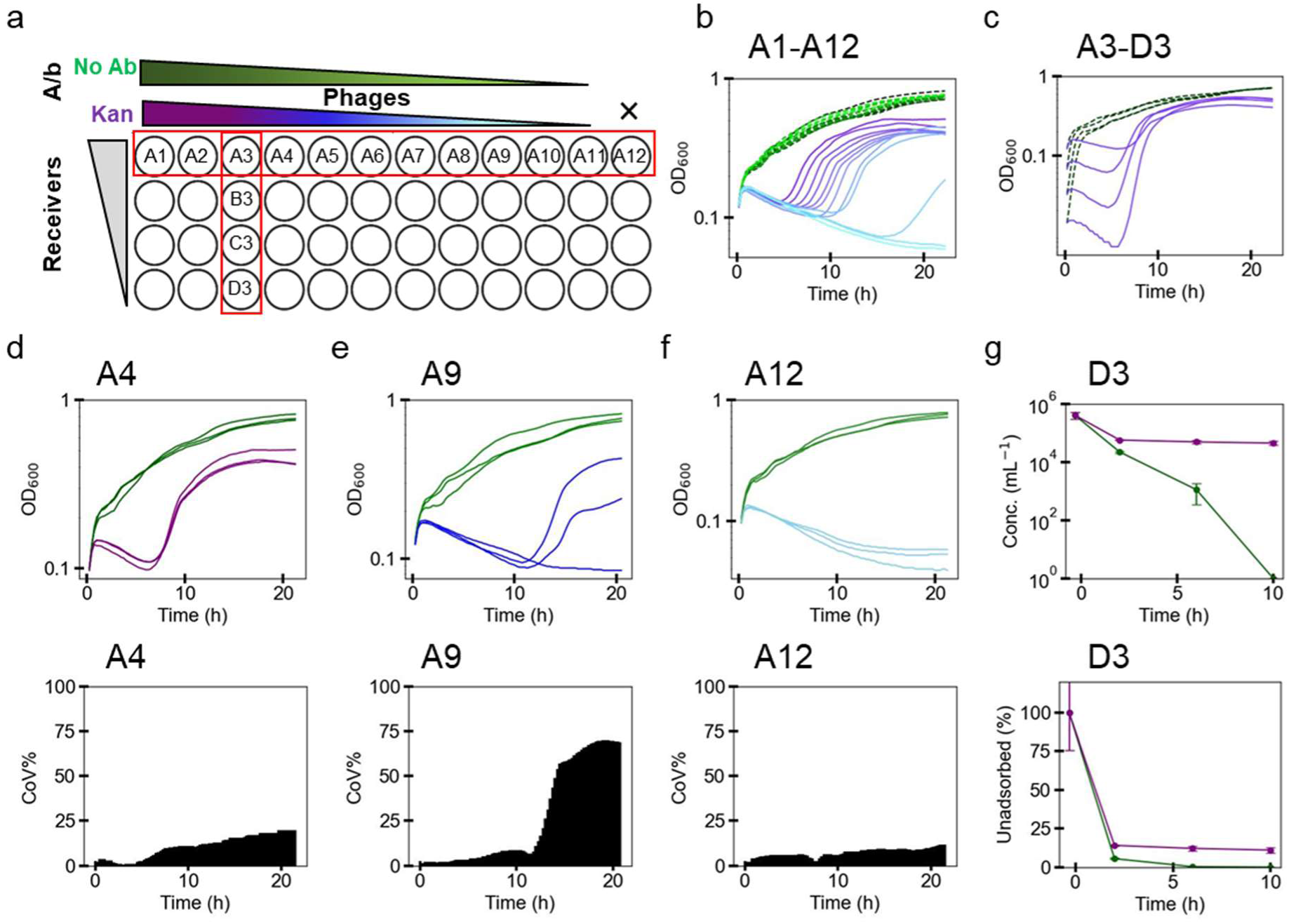
Phage infection dynamics in growing receiver cells exhibits stochasticity. **(a)** A schematic of the experimental setup. Different concentrations of isolated phages (*+gp3φ*) were incubated with receiver cultures (ER2738F) at different densities in a 96-well plate for 20 mins without selection, and subsequently grown with (purple to sky blue) or without (green to light green) antibiotic (kanamycin) selection for infected receivers for ∼20 hours (**Fig.** S4.1). **(b)** Growth curves of receiver cells incubated with varying concentrations of purified phages (wells A1–A12), plotted as OD_600_ against time. **(c)** Growth curves of receiver cells at varying starting densities incubated with a fixed concentration of isolated phages (wells A3–D3), plotted as OD_600_ against time. N=1 for (b) and (c). **(d-f)** (top panel) Repeats of growth curves from wells A4, A9 and A12 (includes data from (b)), plotted as OD_600_ against time (bottom panel) Coefficients of variation (CoV) from the growth curve repeats in the top panel, plotted as CoV against time. Data in (d)-(f) are from N=3 repeats. **(g)** (top panel) Number of unabsorbed phages over time and (bottom panel) %unadsorbed phages over time in well D3, grown with (purple) or without (green) antibiotic selection. Data from N=3 repeats. (b-g) use the colour code defined in (a).

With antibiotic selection, we observed a phage dose-dependent bacterial growth rescue at a fixed starting receiver density (**Fig.** 2b, S4.2). When the same number of phages were incubated with different densities of receiver cells, a similar growth rescue dependent on receiver density was observed (**Fig.** 2c, **Fig.** S4.3). Repeating these experiments three times with a subset of phage dilutions (3^1, 3^2, 3^3, 3^8, 3^9, 3^10) confirmed that growth rescue is more likely at higher phage concentrations (= lower dilutions) across all receiver densities (**Fig.** 2d-f, top panel; **Fig.** S4.4). This is also indicated by the lower coefficients of variation (%CoV) at higher phage concentrations (**Fig.** 2d-e, bottom panel; **Fig.** S4.5). These differences in growth rescue reflect varying probabilities of infection across different phage-receiver concentration settings.

We visualised growth rescue variability using heatmaps of %CoV at the 18 h end-point (**Fig.** S4.6) and cumulative %CoVs over time (**Fig.** S4.7). Both heatmaps confirmed that infection variability is lower at high phage concentrations and higher at low phage concentrations. Interestingly, variability peaked at low phage and low receiver concentrations (**Fig.** S4.6 and S4.7), before reducing again at the lowest receiver OD of 0.03. This indicates that infection probabilities are in the stochastic regime when phage and receiver concentrations are low. At high phage and high receiver numbers, infection probability is high; at low phage and very low receiver numbers, the infection probability is low, with both these conditions representing deterministic infection regimes.

To observe phage uptake rates, we quantified unadsorbed phages from wells A3–D3 (**Fig.** 2a, phage dilution 3^2) at 2, 6, and 10 hours (**Fig.** 2g). In kan(-) wells, phages were progressively adsorbed until almost none remained by 10 hours, driven by continuous multiplication of uninfected receiver cells. In kan(+) wells, ∼85% of phages were adsorbed within 2 hours, with no further change later, indicating that uninfected cells were killed by kanamycin, and only infected receivers grew thereafter (**Fig.** 2g, S4.8). The absence of further phage depletion in kan(+) wells also suggests that phage depletion primarily occurs due to adsorption by uninfected receiver cells, not repeat adsorption by infected receivers. Infected receivers in these experiments are immune to a secondary infection due to the expression of the *gp3* gene delivered by the +gp3φ phages (**Fig.** S3.1).

### Resource competition and antibiotic selection during communication between sender and receiver cells

Processes of horizontal gene transfer transmit valuable functions encoded on DNA from one cell to another. When these cells compete for growth in the same environment, competition for growth can affect the recipient’s ability to take advantage of the newly acquired functions (54). Using a synthetic setup, we next studied intercellular communication dynamics between phage-secreting sender and susceptible receiver cells in a co-culture. Senders (-gp3φ and +gp3φ) and receivers were co-incubated without antibiotics for 1 h, followed by growth under four conditions: without antibiotics, or with antibiotics to select for receivers only (tetracycline), senders only (gentamycin), or infected-receivers only (kanamycin+tetracycline) (**Fig.** 3a, **Note** S5).

**Fig. 3:**
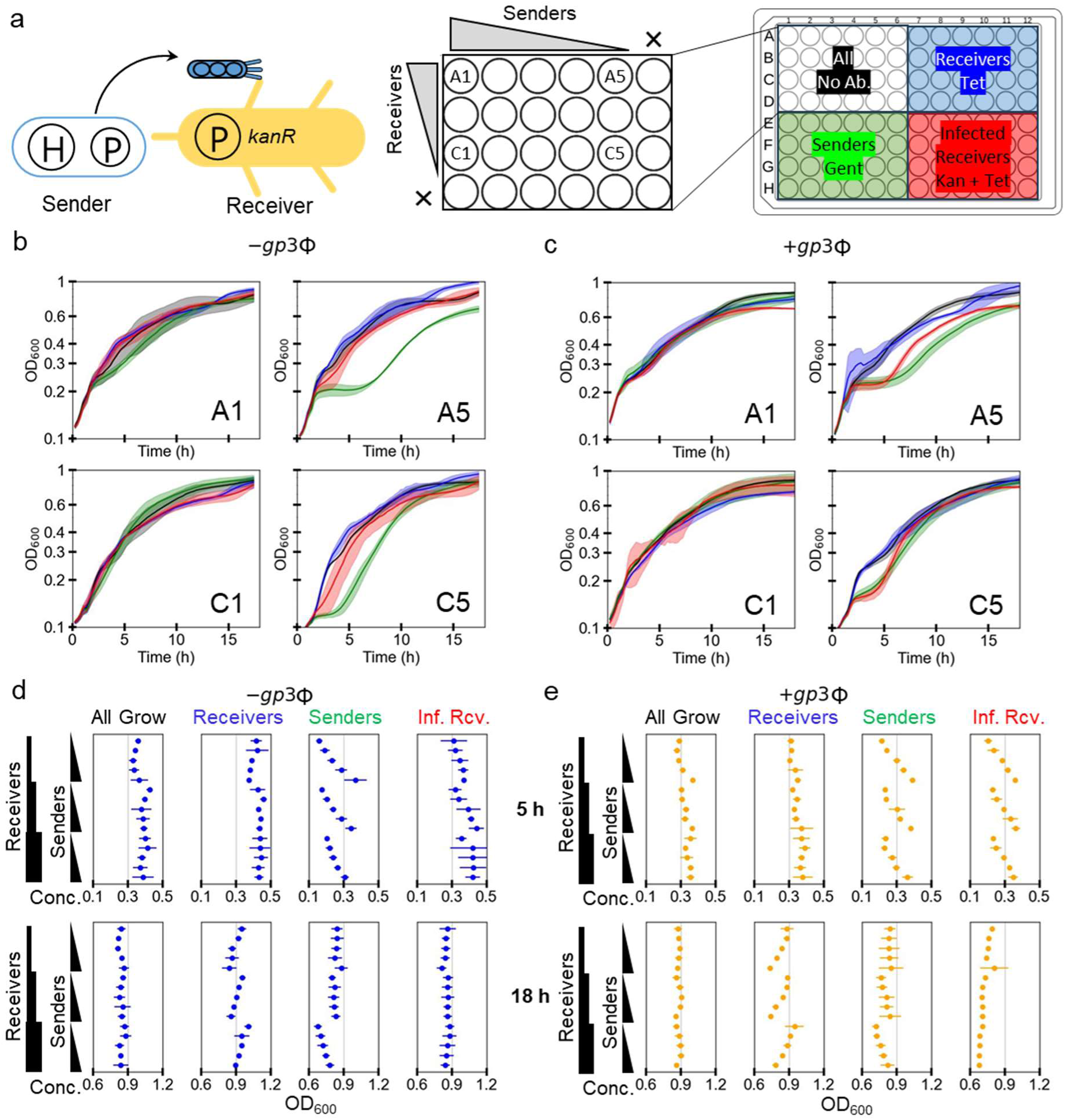
Phage-mediated communication of DNA messages from growing senders to growing receivers. **(a)** A schematic of the experimental setup. Sender (*-gp3φ* or *+gp3φ*) and receiver cells were mixed at different starting densities in a 96-well plate, in four sets, and grown for 1 h without selection. Following that, different antibiotic selections were applied to three of the four sets to select for different cells: tetracycline for receivers only, gentamycin for senders only, and tetracycline+kanamycin for infected receivers only. For each set, initial sender (x-axis) and receiver densities (y-axis) were varied, starting from the undiluted (OD_600_ = 0.136), in 2-fold dilution steps, with the rightmost/bottommost column/row having no sender/receiver cells, respectively. **(b-c)** Growth curves of co-cultures in selected wells (A1, A5, C1, C5) plotted as OD_600_ against time in all the four sets, with different starting sender and receiver densities, using the colour code defined in (a). Data from *-gp3φ* senders are in (b) and those from *+gp3φ* are in (c). **(d-e)** Plots of cell densities (OD_600_) reached at a middle (5 h, top panel) and the end (18 h, bottom panel) time-points in wells with different starting sender and receiver densities, under the different selection conditions in the four sets. X-axis shows the density reached, and y-axis shows the starting sender and receiver cell densities added. (d) and (e) show data from both *-gp3φ* and *+gp3φ* senders for growth in the four selection conditions: no antibiotic (all cells), tetracycline (receivers only), gentamycin (senders only), and tetracycline+kanamycin (infected receivers only). Data in (b)-(e) show mean±SD from N=3 repeats.

Growth dynamics in these conditions revealed how resource competition and antibiotic selection impacted communication (**Fig.** 3b-c, S5.1, S5.5). Without antibiotic selection, both senders and receivers showed a monophasic growth. Similarly, growth in the receivers only condition showed monophasic growth indicating that senders are rapidly killed by tetracycline, and subsequent growth is by receivers only. In co-cultures where two cell types compete for the same resources, biphasic growth curves often reflect sequential dominance due to the initial growth of one cell type followed by the subsequent growth of the second one (55), with a biphasic lag between the two growth phases. Such biphasic growth is seen in the senders only (gentamycin) condition, and is especially apparent when starting sender OD is low (**Fig.** S5.1). As gentamycin killing of receiver cells is slow, they grow for some time while competing with the senders for nutritional resources before dying. Therefore, wells with more starting receivers (A5) show the biphasic lag later than those with fewer starting receivers (C5), even when the starting senders in the two wells are the same. These observations are broadly similar between the *-gp3φ* and *+gp3φ* senders (**Fig.** S5.1 & S5.5). Biphasic growth was also observed in infected-receivers only (kanamycin+tetracycline) conditions, especially for +gp3φ senders. This is because the secretion rates of *+gp3φ* senders are ∼1000-fold lower than those of *-gp3φ* senders (**Fig.** 1f), resulting in more receivers remaining uninfected at the end of the 1 h pre-selection incubation. In the -gp3φ case, we estimate that >75% receivers are infected before the selection is applied, allowing them to quickly outcompete both senders and uninfected receivers without an apparent biphasic lag. In contrast, <1% receivers are infected in the +gp3*φ* case during the same pre-selection period.

To further understand the contributions of sender and receiver numbers, we plotted ODs of several wells at an early time-point (5 h) and the end-point (18 h) of post-selection growth (**Fig.** 3d-e). More detailed plots are available in **Note** S5. Without antibiotics in the - gp3φ experiments, wells with only senders added had lower end-point ODs than those with only receivers added, suggesting that in 1:1 sender:receiver co-cultures -gp3φ senders would contribute less to the final OD than receivers (**Fig.** S5.2-S5.3).

In the +gp3φ experiments, wells with only senders added reach end-point ODs similar to wells with only receivers added (**Fig.** S5.6-S5.7). These differences in end-point ODs reflect the fact that growth rate difference between receivers and -gp3φ senders is much higher than that between receivers and +gp3φ senders (**Fig.** S5.11). In receiver only growth conditions, end-point ODs in both -gp3φ and +gp3φ experiments were lower in wells with higher starting senders (**Fig.** 3d-e), an effect more pronounced for the +gp3φ senders. This is due to early resource competition between receivers and senders before the latter are killed by tetracycline. In sender only growth conditions, ODs were higher at 5 h in wells with more starting senders as expected, but plateaued by 18 h with weaker dependence on starting sender numbers (**Fig.** 3d-e). However, the effect of early resource competition with the receivers was still visible, resulting in lower ODs in wells with higher starting receivers (**Fig.** 3d-e). These results were also seen when comparing a smaller subset of the data from pairs of wells with similar total starting ODs but different sender:receiver ratios (**Fig.** S5.4, S5.8). Overall, the resource competition effect was much smaller when selecting for senders only than seen above when selecting for receivers only, due to receivers being less rapidly killed by gentamycin than senders are killed by tetracycline.

In the infected-receivers condition at an early time-point (5 h), as expected the OD is higher for wells with higher starting senders. This effect of sender dose dependence was more pronounced for +gp3φ senders than -gp3φ senders (**Fig.** 3d-e), due to the former’s lower secretion rates that result in lower phage titres by the end of the 1 h incubation (**Fig.** 1g). Variability between repeats in sender-receiver communication experiments (**Fig.** S5.9-S5.10), for both senders, was lower than in phage-receiver infections (**Fig.** 2d-f, S4.5). We estimate that <2 cells per mL of -gp3φ senders or <∼22000 cells per mL of +gp3φ senders would be needed for sender-receiver infections to exhibit high variability similar to phage-receiver infections. The end-point ODs plateaued if any senders were present, with higher ODs for - gp3φ than +gp3φ senders. Within each set with fixed starting receivers, increasing sender numbers marginally reduced the end-point OD (**Fig.** 3d-e, S5.2, S5.6). For -gp3φ, the starting receiver OD had little effect on infected-receiver end-point ODs (**Fig.** S5.3), but for +gp3φ, higher starting receiver OD led to lower final OD (**Fig.** S5.7). This is due to early resource competition with more receivers that later die if uninfected. This suggests that when the phage-mediated communication rate is low (as for +gp3φ) and conditions are selective for infection, vertical transfer of the phagemid from mother to daughter cell is more efficient than horizontal transfer from sender to receiver cell.

### Intercellular gene regulation using CRISPRi circuits

Over the past decade, programmable CRISPR nucleases and their catalytically dead mutants have been widely used in genome engineering and gene regulation applications (56). CRISPR systems are standalone in their function, with the expression of the CRISPR nuclease and guide RNA being sufficient for activity, even in heterologous settings or *in vitro* (56). The specificity of CRISPR systems can be programmed by modifying the guide RNA sequence to direct the ribonucleoprotein to a specific DNA or RNA target. This versatility has made CRISPR systems valuable tools for building information processing genetic circuits (8, 23), acting as internal wires in the cell, or functioning as DNA payload selectors (27).

Building on the phage-mediated cell-to-cell communication (**Fig.** 3), we aimed to create multicellular circuits where DNA messages coding for CRISPR guide RNAs are secreted by sender cells and delivered by phages to modify receiver gene expression. We term this process “intercellular CRISPRi” (i-CRISPRi). We began by examining the infection dynamics and circuit response of a simple i-CRISPRi circuit (**Fig.** 4a), where a single guide RNA (*sgRNA-1*) targets the template DNA strand downstream of the GFP promoter for transcriptional repression (see **Fig.** S10.1a). The sgRNA is encoded on the -gp3φ phagemid variant, which has a higher secretion rate and lacks superinfection inhibition. Co-transformation of the dCas9-GFP plasmid, which expresses dCas9 from a constitutive promoter and GFP from an sgRNA-repressible promoter, with a phagemid constitutively expressing *sgRNA-1* showed high repression of the GFP (**Fig.** S6.1). Next, isolated sgRNA-1 phages (3.6x10^13^ mL^-1^, CFU assay) were incubated with receiver cells at 0.125 OD without antibiotic selection, GFP expression was monitored for 5 hours by flow cytometry, and receiver cells were gated into ‘OFF’ and ‘ON’ populations (**Fig.** 4b).

Upon infection, the phagemid delivered into the receivers expressed *sgRNA-1* from a strong constitutive promoter (J23119), and in turn formed the dCas9-sgRNA complex to repress the *GFP* gene, thereby increasing the %OFF cells in the population (**Fig.** 4c). GFP expression was repressed in 84% of receiver cells by 2 hours, and 90% cells by 5 hours. These results are consistent with those reported in the first phage-derived DNA messaging system where 92% receivers got infected by phages from sender cells within 5 h without antibiotic selection (42). To compare phage-mediated CRISPRi against small-molecule inducible CRISPRi, we separately co-transformed the dCas9-GFP plasmid with two *sgRNA-1* plasmids, inducible by arabinose and IPTG. GFP repression was monitored after 16 hours of induction using varying inducer concentrations (**Fig.** S6.2, S6.3). For comparison, we also incubated the previously used dCas9-GFP receiver cells with varying concentrations of *sgRNA-1* phages for 16 h (**Fig.** S6.4). Flow cytometry data show that small molecule induction of *sgRNA-1*, with both arabinose and IPTG, results in tunable control of *GFP* repression, whereas the delivery of the sgRNA-1 phagemid results in a digital ON-OFF behaviour. This indicates that DNA messages, unlike small molecules, behave like digital signals; a single successful transmission delivers the full message to the receiver cell where its expression can act on the downstream circuit.

After confirming GFP repression by *sgRNA-1* in phage-transduced receiver cells, we constructed the i-CRISPRi circuit with senders secreting the sgRNA phagemid and receivers expressing dCas9 and GFP. Upon co-culturing of the sender and receiver cells, sgRNA phages produced by the senders can transduce into the receivers (**Fig.** S6.5). Co-culturing sender and receiver cells in varying ratios showed that 87% of receivers expressed ∼20-fold lower GFP at a 1:4 sender:receiver ratio within 5 hours (**Fig.** 4d). Higher sender ratios (2:4 or 4:4) further increased the GFP-repressed population to 92% and 95%, respectively. Increasing sender numbers, while keeping the receiver numbers the same, slightly reduced the time to repress 50% of receivers from 2.29 h (1:4 ratio) to 1.98 h (2:4 ratio) and 1.76 h (4:4 ratio). Sender:receiver ratios of 4:4, 4:2, and 4:1 showed no significant difference in repression rates, suggesting excess phage secretion at those sender numbers (**Fig.** 4e).

We calculated the phagemid transfer frequency from senders to receivers, using a method used to obtain transconjugation frequency in conjugation experiments (27). We found that the highest transfer frequency of 3x10^-4^ mL cell^-1^ was observed at 3 hours of co-incubation for low sender and high receiver starting densities (**Fig.** 4f, S6.6). This transfer rate vastly exceeds the 5.2x10^-9^ reported for transconjugation at 6 hours (27). Two additional i-CRISPRi circuit variants with different sgRNA and cognate promoter sequences, as well as different replication origins, confirmed the general applicability of these circuits (**Fig.** S6.7, S6.8). However, it also revealed differences in the rate of GFP repression (**Fig.** S6.9), possibly because the different replication origins used affect phage communication rates, or the different sgRNAs have different rates of repression after phagemid delivery to the receiver cells.

### Single-input Boolean logic gates using intercellular CRISPRi

After demonstrating i-CRISPRi regulation (**Fig.** 4), we designed multicellular circuits implementing single-input Boolean logic gates. Previously, we had tested i-CRISPRi circuits without antibiotic selection, but selection may be needed as circuit complexity and the number of inputs increase. So, we examined how circuit output changes with and without selection for two single-input gates: NOT and YES.

The NOT gate was built using sgRNA-1 encoding phagemid secreted by senders to infect receiver cells expressing GFP and dCas9 (**Fig.** 5a). This circuit uses a “single-rail” encoding, where the sender’s presence represents input ‘1’ and its absence represents input ‘0’. Sender-receiver co-cultures were grown for 4 hours without selection, followed by 16 hours of growth under four conditions: no antibiotic, selection for receivers only (spectinomycin), senders only (gentamycin), and infected-receivers only (spectinomycin+ampicillin) (**Fig.** 5b, S7.1). Flow cytometry showed that infected receivers exhibited a 21.3-fold decrease in GFP expression compared to the uninfected receiver control (**Fig.** 5c). Interestingly, selection for sgRNA phagemid was not necessary, as setups without antibiotics or selection for receivers only (spectinomycin) showed similar results (**Fig.** 5d, S7.2).

**Fig. 4:**
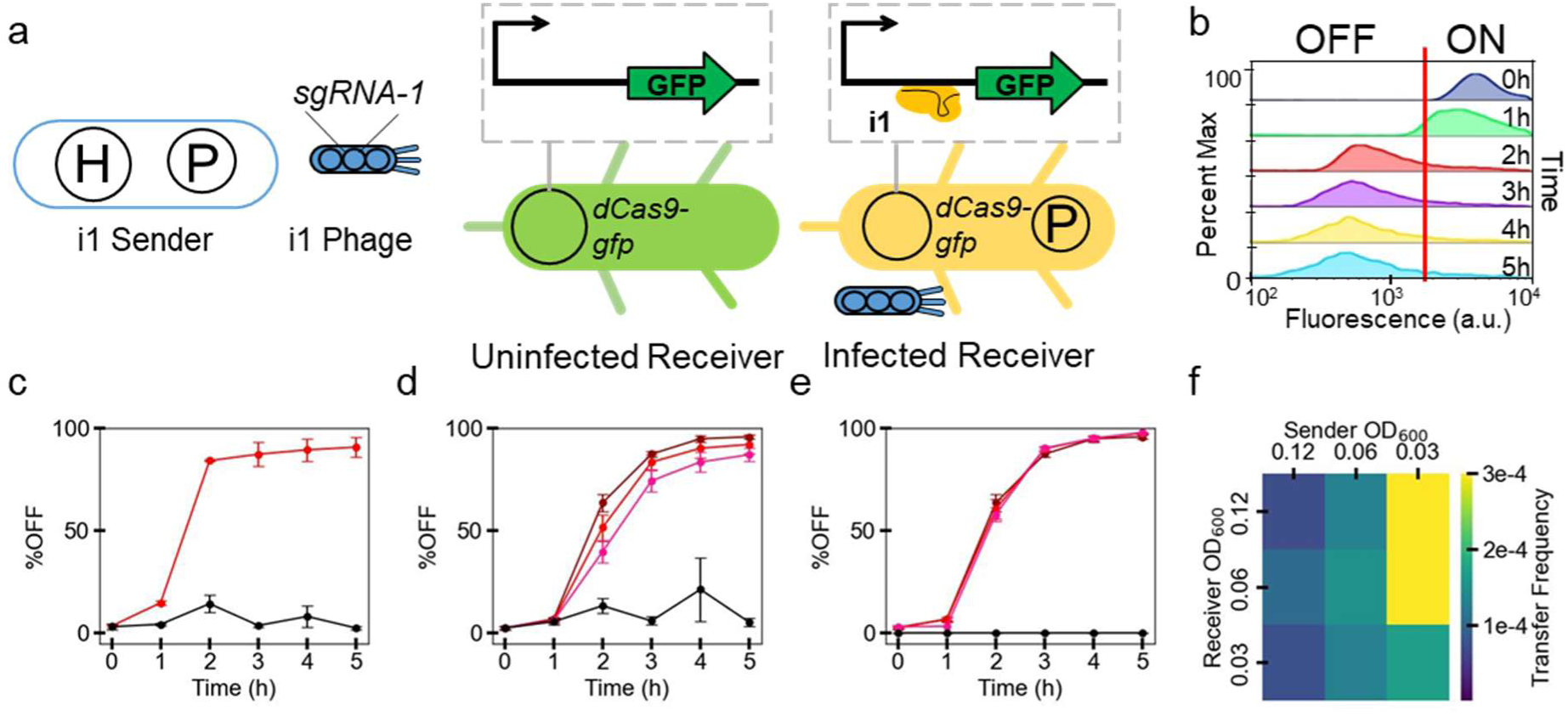
Phagemids encoding CRISPR single guide RNA can regulate receiver gene expression. **(a)** A schematic of the intercellular CRISPR interference system. Sender i1 secretes phage particles encoding *sgRNA-1*. The phage particle infects a receiver cell (TOP10F_dCas9-GFP_NOT_i1), resulting in the delivery of the sgRNA-encoding DNA that subsequently expresses *sgRNA-1* in the receiver cells. sgRNA-1 forms a complex with available dCas9 and binds to the promoter region of *GFP*, causing repression. **(b)** Distribution of GFP fluorescence in receiver cells incubated with isolated sgRNA-1 phages for varying durations of up to 5 h, without antibiotic selection. Number of GFP ON and OFF cells was determined using a fluorescence threshold value (vertical red line). These representative plots are from a single experiment (N=1). **(c)** Percentage of repressed (GFP OFF state) cells in the receiver population, sampled every hour for 5 hours. Repression is shown from receiver cells with (red) or without (black) added phages. Data show mean±SD from N=3 repeats. **(d-e)** Receiver GFP repression across time is shown when incubated with sgRNA-1 senders (TOP10_H_sgRNA-1_AmpΦ) without selection. Like in (c), the percentage of repressed receivers is plotted against time. Data in (d) are from a fixed starting receiver density (OD_600_ = 0.125), and varying starting sender densities: OD_600_ 0.125 (dark red), 0.0625 (red), 0.0312 (pink), no sender (black). Data in (e) are from a fixed starting sender density (OD_600_ = 0.125), and varying starting receiver densities: OD_600_ 0.125 (dark red), 0.0625 (red), 0.0312 (pink), no receiver (black). Data in (c)-(e) show mean±SD from N=3 repeats. **(f)** Heatmap showing transfer frequency (mL cell^-1^) at 3 h of co-culturing, calculated for different starting sender and receiver ODs.

**Fig. 5:**
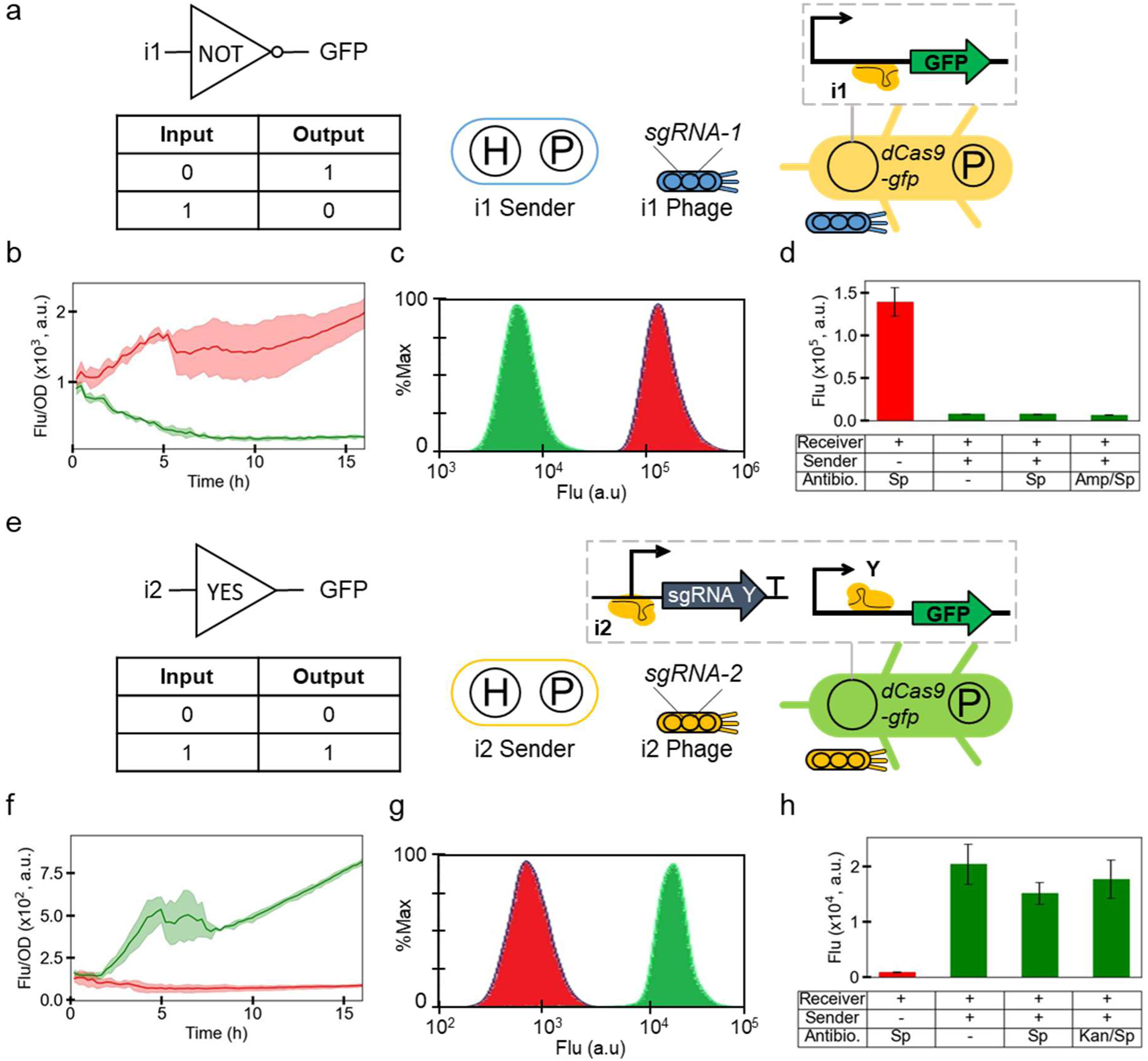
Intercellular CRISPRi for digit logic gates NOT and YES. **(a)** Schematic of the intercellular CRISPR interference system for NOT logic gate behaviour. Sender i1 secretes phage particles encoding *sgRNA-1*. Upon infection and expression in the receiver cells, *sgRNA-1* represses *GFP* expression by CRISPRi. **(b)** NOT-gate uninfected receivers (red), and infected receivers (green), were grown for 16 hours in a plate reader, with OD_600_ and fluorescence recorded at regular intervals. A time-course of Fluorescence/OD is plotted here. **(c)** Distribution of GFP fluorescence in receiver cells incubated without or with sgRNA-1 senders for 16 hours, with appropriate antibiotic selection (without sender = red, spec; with sender = green, spec+amp). These representative plots are from a single experiment (N=1). **(d)** Same experiment as in (c), except two additional growth conditions with senders are shown (no antibiotic and spec). Mean±SD of fluorescence data from N=3 repeats are plotted here. **(e)** Schematic of the intercellular CRISPR interference system for YES logic gate behaviour. Sender i2 (TOP10_H_sgRNA-2_KanΦ) secretes phage particles encoding *sgRNA-2*. Upon infection and expression in the receiver cells (TOP10F_dCas9-GFP_YES_i2), *sgRNA-2* represses *sgRNA-Y*, in turn depressing *GFP* expression. **(f)** YES-gate uninfected receivers (red), and infected receivers (green), were grown for 16 hours in a plate reader, with OD_600_ and fluorescence recorded at regular intervals. A time-course of Fluorescence/OD is plotted here. **(g)** Distribution of GFP fluorescence in receiver cells incubated without or with sgRNA-2 senders for 16 hours, with appropriate antibiotic selection (without sender = red, spec; with sender = green, spec+kan). These representative plots are from a single experiment (N=1). **(h)** Same experiment as in (g), except two additional growth conditions with senders are shown (no antibiotic and spec). Mean±SD of fluorescence data from N=3 repeats are plotted here.

The YES gate (buffer gate) circuit used sender cells encoding sgRNA-2 and a receiver circuit with two sequential NOT gates (inverters) (**Fig.** 5e). Receiver cells carried the YES gate circuit on a plasmid expressing *sgRNA-Y* and dCas9, with their complex repressing the GFP promoter. sgRNA-Y promoter, in turn, was regulated by the *sgRNA-2*. YES gate sender and receiver cells were co-cultured for 4 hours without selection, followed by 16 hours under the same four conditions used previously (**Fig.** 5f, S7.3). Flow cytometry showed a 20.7-fold increase in GFP expression in infected receivers compared to the uninfected receiver control (**Fig.** 5g). Similar to the NOT gate, selection for sgRNA was not required in the receiver cells (**Fig.** 5h, S7.4). High GFP activation was observed after 16 hours without selection, but not after 4 hours of pre-incubation (**Fig.** S7.5). This is different to the circuit behaviour previously observed with the sgRNA-1 NOT gate senders (**Fig.** 4d), where 4 h of pre-incubation without selection was already sufficient to reach 95% repression. This suggests either lower sgRNA-2 phage secretion rates or a longer delay between infection and activation in the YES gate circuit, or a combination of both.

Post-selection data showed that the YES gate fluorescence switch requires more time than the NOT gate, where a response was seen almost immediately (**Fig.** 5b, 5f, S9.1). Defining circuit switching time as a ≥20% change from the no-input control fluorescence, the NOT gate switch begins at 39.6 min, while the YES gate switch starts at 173.4 min (2.89 hours) (**Fig.** S9.1). The longer YES gate switch time is likely due to the additional CRISPRi step in the repression cascade.

### Multi-input Boolean logic gates using intercellular CRISPRi

Before implementing multi-input logic gates, we checked if receiver cells could be infected by multiple phages simultaneously. Receiver cells were incubated with 1, 2, or 3 phage types (differing in replication origins and antibiotic markers) for 1 hour, then subjected to antibiotic selection. Results showed that cells could be co-infected by multiple -gp3φ phagemids simultaneously (**Fig.** S3.2), but not if already infected with a +gp3φ phagemid (**Fig.** S3.1). While multiple -gp3φ infections are possible, multiple antibiotic selections reduced receiver cell growth. Yet, given the lack of circuit activation at 4 h without selection seen earlier in the YES gate (**Fig.** S7.5), antibiotic selection seems necessary if circuits are to be activated by multiple inputs. However, comparing fluorescence output under different growth conditions is challenging due to growth rate differences affecting GFP dilution (57). To address both these requirements together, we used “dual-rail” encoding, where circuit input comes from two types of sender cells delivering a ‘0’ signal (dummy sgRNA) or a ‘1’ signal (targeting sgRNA).

The AND gate (A.B) receiver uses a plasmid encoding GFP, dCas9, *sgRNA-X*, and *sgRNA-Y*. In complex with dCas9, either of the two sgRNAs represses GFP by binding downstream of the promoter (**Fig.** 6a). This promoter configuration with tandem repressor binding sites behaves like a NOR gate, which owing to its functional completeness can be layered to build any digital logic (9). Promoters of *sgRNA-X* and *sgRNA-Y* are regulated by *sgRNA-A* and *sgRNA-B*, respectively. Two sender cells secrete phages encoding sgRNA-A and sgRNA-B for signal input ‘1’ (TOP10_H_sgRNA-A1_KanΦ and TOP10_H_sgRNA-B1_GentΦ), while two others secrete null phages encoding dummy sgRNAs for signal input ‘0’ (TOP10_H_sgRNA-A0_KanΦ and TOP10_H_sgRNA-B0_GentΦ). The ‘1’ and ‘0’ senders of any input set are never used together. The AND gate was implemented in a 2-sender 1-receiver circuit, incubated with all combinations of senders (‘00’, ‘01’, ‘10’, and ‘11’) for 4 hours without selection, then grown for 16 hours with multi-antibiotic selection in a plate reader (**Fig.** 6b, S8.1). Using a conservative fold-change estimate proposed in the Cello paper (58), flow cytometry showed a 14.3-fold increase in fluorescence with the ‘11’ sender combination compared to the highest of the other three combinations (**Fig.** 6cd, S8.2).

**Fig. 6:**
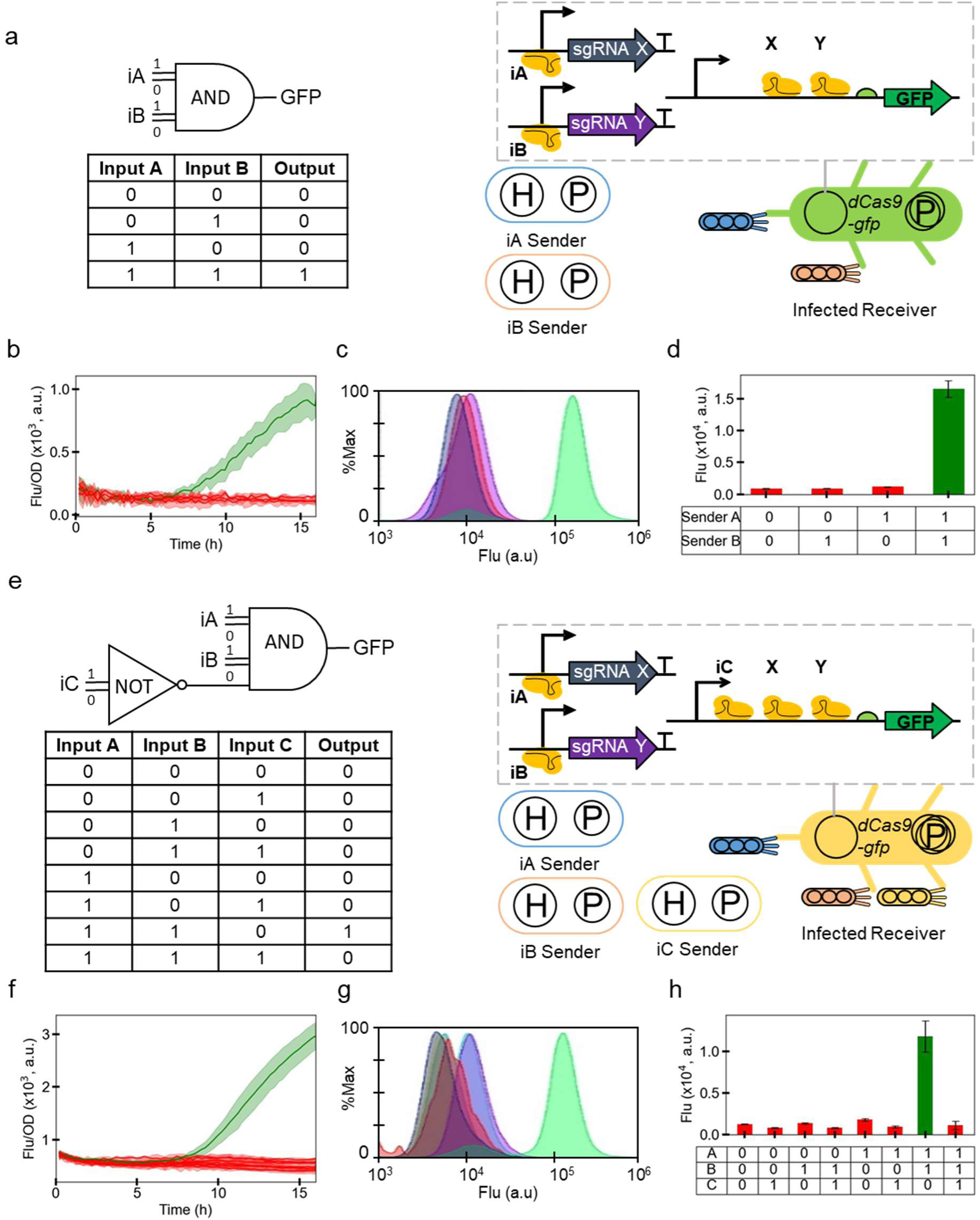
Intercellular CRISPRi for digit logic gates AND and AND-AND-NOT. **(a)** Schematic of the intercellular CRISPR interference system for AND logic gate behaviour. Senders iA and iB secrete phage particles encoding either null (state ‘0’) phages, or *sgRNA-A* and *sgRNA-B* (state ‘1’) phages. Only upon infection of receiver cells by both state ‘1’ phages, *sgRNA-A* and *sgRNA-B* repress *sgRNA-X* and *sgRNA-Y*, respectively, in turn depressing *GFP* expression. **(b)** AND-gate receivers (red) infected by non-activating combinations of phage senders (00, 01, 10), and receivers (green) infected by activating combination (11) of phage senders, were grown for 16 hours in a plate reader, with OD_600_ and fluorescence recorded at regular intervals. A time-course of Fluorescence/OD is plotted here. **(c)** Distribution of GFP fluorescence in receiver cells incubated with sgRNA-A and sgRNA-B senders for 16 hours, with appropriate antibiotic selection. These representative plots are from a single experiment (N=1). **(d)** Same experiment as in (c), except the mean±SD of fluorescence data from N=3 repeats are plotted here. **(e)** Schematic of the intercellular CRISPR interference system for AND-AND-NOT logic gate behaviour. Senders iA, iB, and iC secrete phage particles encoding either null (state ‘0’) phages, or *sgRNA-A*, *sgRNA-B*, and *sgRNA-C* (state ‘1’). Only upon infection of receiver cells by state ‘1’ phages A and B and state ‘0’ phage C, *sgRNA-A* and *sgRNA-B* repress *sgRNA-X* and *sgRNA-Y*, respectively, in turn depressing *GFP* expression. If state ‘1’ phage C infects receivers, it directly represses *GFP*. **(f)** AND-AND-NOT gate receivers (red) infected by non-activating combinations of phage senders (000, 001, 010, 011, 100, 101, 111), and receivers (green) infected by activating combination (110) of phage senders, were grown for 16 hours in a plate reader, with OD_600_ and fluorescence recorded at regular intervals. A time-course of Fluorescence/OD is plotted here. **(g)** Distribution of GFP fluorescence in receiver cells incubated with sgRNA-A, sgRNA-B, and sgRNA-C senders for 16 hours, with appropriate antibiotic selection. These representative plots are from a single experiment (N=1). **(h)** Same experiment as in (g), except the mean±SD of fluorescence data from N=3 repeats are plotted here.

The AND-AND-NOT (A.B.∼C) gate receiver (TOP10F_dCas9-GFP_AAN_4) has a circuit plasmid with an sgRNA binding site downstream of the *GFP* promoter for repression by an additional sgRNA-C produced by sender C (TOP10_H_sgRNA-C1_AmpΦ) (**Fig.** 6e). The AND-AND-NOT gate circuit presented here was identified from 7 designs (**Fig.** S10.1) using a 3-step characterisation process (see **Note** S10, **Fig.** S10.2 & S10.3). The circuit involves three sender cells for signal ‘1’ (sgRNA-A, sgRNA-B, sgRNA-C) and three for signal ‘0’ (dummy sgRNAs). The AND-AND-NOT gate was implemented in a 3-sender 1-receiver circuit, tested with combinations (‘000’, ‘001’, ‘010’, ‘011’, ‘100’, ‘101’, ‘110’, ‘111’) for 4 hours without selection, followed by 16 hours with antibiotics (**Fig.** 6f, S8.3). Using flow cytometry, a 7.7-fold increase in GFP expression was observed between the ON-state (‘110’) and the highest of all OFF-states (**Fig.** 6gh, S8.4). Senders A1, B1, and null sender C0 (TOP10_H_sgRNA-C0_AmpΦ) were incubated with receivers to achieve the ON-state.

The 20% switching response time increased to 8.52 hours for AND gate and 10.71 hours for AND-AND-NOT gate compared to single-input circuits (**Fig.** 5, 6b, 6f, S9.1), due to the time lag from additional CRISPRi steps required in the repression cascade (**Fig.** S9.1).

## Discussion

In this work, we have leveraged key advantages of a phage-derived DNA messaging system, high programmability and message-channel decoupling (42), to implement several distributed logic gates in multi-strain bacterial consortia. We investigated the transmission (secretion) and reception (infection) dynamics of the messaging system, revealing that both steps are significantly influenced by the growth phase of sender and receiver cells, though in contrasting ways. Secretion rates decrease as senders transition from early to late growth phases (**Fig.** 1g), whereas infection rates of receivers increase during a similar transition (**Fig.** 1j). This emphasizes the need to assess phage production and infection kinetics not just in the exponential phase, as is commonly done, but across different growth phases. It is especially relevant since bacteria in many natural environments exist in different phases of growth with potentially different cell surface properties that influence phage infection (59, 60). Additionally, the phage yield and secretion rate varied between -gp3φ and +gp3φ sender variants, suggesting that phage machinery expression level is crucial for optimal phage production (**Fig.** 1f-g).

Studying phage infection kinetics in growing receiver cells, we found high infection variability with low phage and receiver concentrations (**Fig.** 2d-f, S4.5). This is consistent with previous findings that implicated stochastic phage-bacterial interactions, due to spatial heterogeneity in the gut, for the poor efficacy of phage therapy (61). Using sender cells rather than isolated phages reduced this variability substantially, due to phage dose amplification (**Fig.** S4.5, S5.9–5.10). This suggests that using sender cells instead of isolated phages for phage-mediated DNA delivery can improve outcomes in phage therapy and microbiome editing applications. However, phage communication kinetics between senders and receivers are shaped by resource competition among cells. Increasing sender ratios does not guarantee more infected receivers, as too many senders can deplete the resources needed for receiver growth (**Fig.** 3). Therefore, horizontal transmission to a few receivers followed by vertical transmission by growth of those infected receivers might be a more effective strategy to obtain more infected receivers containing phage DNA.

Our results demonstrate that the rates of M13 phage-mediated intercellular gate activation compare favourably against both small molecules and conjugation. Using isolated phages for induction of our simplest circuit (NOT gate), we achieve 20% induction in 64 min (**Fig.** 4c), compared to the 35–102 minutes using different concentrations of HSL molecules in previous studies (62, 63). Using phage-secreting sender cells to communicate with receivers in a co-culture, we achieved 20% activation in 42 minutes (**Fig.** S9.1), which is faster than the 146 minutes previously seen for AHL molecules (64). Due to the slow signal accumulation in sender-receiver co-cultures, many small molecule communication experiments instead use conditioned media for receiver activation (65, 66), or other specialised strategies to enrich signal accumulation (9). For example, a recently implemented distributed solution for a cryptographic problem built a set of 41 bacterial strains, each containing a different subcircuit, that can communicate using 4 small molecules (66). However, communication between pairs of strains was primarily achieved using conditioned media, with only two example pairs tested by co-culturing. Our system demonstrated significant fold activation of circuits without such enrichment strategies: 21-fold (**Fig.** 5d), 14.3-fold (**Fig.** 6d), and 7.7-fold (**Fig.** 6h) for single-, dual-, and triple-input circuits, respectively. However, these are lower than fold-changes of ∼6–125-fold seen with 2–4 input unicellular circuits using transcription factors or CRISPRi (23, 67). In a recent preprint, Kusumawardhani *et al.* report similar findings using phage-mediated communication in multicellular circuits (68). While our work characterises the rates of phage-mediated communication in detail and uses up to 6 sender strains as inputs, Kusumawardhani *et al.* use up to 4 small molecules as inputs combined with the inducible secretion of 2 intercellular phage signals.

M13-mediated communication was faster than conjugation, with ∼95% of receivers infected within 5 hours (**Fig.** 4d), versus 50% in 6 hours using conjugation (27). This is reflected in the transfer frequency obtained using M13 phages (**Fig.** 4f), which is about 5 orders of magnitude higher than for conjugation (27). Furthermore, phage-mediated communication does not require cell-to-cell contact, making it applicable even in sparse populations. Phage messages can stay viable in harsh environments before they find a receiver cell for delivery (69). Unlike conjugation that requires an active F-pilus as DNA conduit (70), M13 needs the F-pilus for the initial surface attachment but not as a DNA conduit into the cell (71, 72). As a result, M13 infection can continue into the stationary phase of cell growth while conjugation cannot (73). In both cases, cell surface receptors can be engineered to modify the transfer rates of both M13 transduction and conjugation (26, 74).

We combined phage-mediated communication with CRISPRi to develop the i-CRISPRi system, implementing NOT and YES gates, as well as AND and AND-AND-NOT gates in co-cultures with up to 4 cell types at a time. Our most complex circuit implemented, the AND-AND-NOT gate, uses a dual-rail encoding with 3 sender strains (of 6 possible sender strains) and 1 receiver strain in a co-culture. The ability of several -gp3φ phages to simultaneously infect the same receiver enables multiplex information processing, which is not possible using many conjugation systems due to surface exclusion after the first transfer event (75). However, conjugation can be used to re-transmit messages as receiver cells can assume the role of senders after the initial message delivery (27). Due to gp3-mediated immunity (46), such re-transmission is not normally possible for -gp3φ phagemids but it is possible for +gp3φ phagemids, provided the receivers carry an appropriate helper. Alternatively, other solutions that combine signal multiplexing with re-transmission could employ conditional expression of *gp3* as done in stringency-modulated PACE (49).

Future enhancements to the M13 messaging system could include selective packaging of orthogonal messages, targeting receivers with specific surface receptors (74), and combining the system with re-addressable delivery of messages (27). Our data suggest that changing plasmid replication origins, or the sgRNAs encoded, can modulate repression rates (**Fig.** S6.9), thereby generating an array of messaging variants with different communication properties. If only transient DNA delivery is required, mini-phagemids with a split M13 packaging signal and no bacterial replication origin can be employed (76). For applications requiring both analogue and digital signals, small molecule communication may be combined with conjugation- and phage-mediated DNA messaging. The phage-mediated i-CRISPRi system developed in this work is amenable for interfacing with other cellular communication systems, thereby expanding distributed computing circuits and DNA delivery applications for microbiome engineering.

## Data Availability

Data presented in all figures has been added to the Source data file in the Supplementary materials.

## Acknowledgements

We thank the two anonymous referees for their thoughtful comments, which we believe have helped improve the manuscript substantially. We thank Jean-Loup Faulon for generous access to laboratory equipment. We thank Alfonso Jaramillo for the kind gift of bacterial strains and plasmids. We thank Vijai Singh and Sai Akhil Golla for their help in cloning some plasmids used in this study. We thank Hadi Jbara, Roman Luchko, Tom Zaplana, and Anchita Sharma for technical assistance with M13 phage protocols. We acknowledge the use of GPT-4o (Open AI, https://chatgpt.com/) for text editing / rephrasing.

## Author contributions

M.F., T.N., and M.K. conceived the study.

A.Pu., A.Pat., and M.K. designed the wet-lab experiments.

A.Pu., A.Pat., C.H., A.Pan., C.R.C., and M.K. performed the wet-lab experiments.

A.Pu., A.Pat., M.F., T.N., and M.K. analysed the data.

M.F., T.N., and M.K. acquired the funding.

A.Pu. and M.K. wrote the manuscript with contributions from all authors. All authors read and approved the final manuscript.

