## Supplementary Information for "Phage-mediated intercellular CRISPRi for biocomputation in bacterial consortia"

**Title**

### Table of Contents:

**Title** 1
**Table of Contents:** 2
**Supplementary Notes** 4
Note S1: PFU and CFU assays 4
Fig. S1.1: Receivers at different physiologies (harvest OD600) infected with the same concentration of phages. 5
Fig. S1.2: Receivers at different densities (resuspended OD600) infected with the same concentration of phages. 5
Fig. S1.3: Relationship between CFU and PFU counts. 6
Note S2: Converting optical density to cell density 7
Fig. S2.1: Conversion of OD600 to cell density. 7
Note S3: gp3 causes infection immunity 8
Fig. S3.1: Expression of gp3 in receiver cells inhibits phage infection. 8
Fig. S3.2: Multiple and simultaneous infection of receiver cells. 8
Note S4: Phage infection in growing bacteria 8
Fig. S4.1: Growth curves of receiver cells infected with 12 different concentrations of phages. 9
Fig. S4.2: Phage dose-dependent growth rescue of receiver culture with different starting OD600. 10
Fig. S4.3: Receiver dose-dependent effect on growth rescue. 10
Fig. S4.4: Phage-dose dependent growth rescue. 11
Fig. S4.5: Coefficients of variation over time between repeats. 11
Fig. S4.6: Heatmap of CoV of OD600 at 18th hour from CoV over time shown in S4.5. 12
Fig. S4.7: Heatmap of the sum of OD600 CoVs through time (0-18 h). 12
Fig. S4.8: Phage adsorption by receiver cells over time. 13
Note S5: Cell-to-cell communication from sender to receiver cells 14
Fig. S5.1: Cell-to-cell communication with -gp3φ senders & receivers. 14
Fig. S5.2: OD600 at 5h and 18h, grouped by starting receiver densities (-gp3φ set). 15
Fig. S5.3: OD600 at 5h and 18h, grouped by starting sender densities (-gp3φ set). 15
Fig. S5.4: OD600 at 5h and 18h, showing data for wells with the same total starting OD of senders and receivers (-gp3φ set). 16
Fig. S5.5: Cell-to-cell communication with +gp3φ senders & receivers. 17
Fig. S5.6: OD600 at 5h and 18h, grouped by starting receiver densities (+gp3φ set). 18
Fig. S5.7: OD600 at 5h and 18h, grouped by starting sender densities (+gp3φ set). 18
Fig. S5.8: OD600 at 5h and 18h, showing data for wells with the same total starting OD of senders and receivers (+gp3φ set). 19
Fig. S5.9: CoV% over Time calculated between 3 repeats for -gp3φ sender set.20
Fig. S5.10: CoV% over Time calculated between 3 repeats for +gp3φ sender set. 21
Fig. S5.11: Specific growth rates of receiver, senders and infected receiver

strains. 22

Note S6: Using CRISPRi to track real-time receiver infection 23

Fig. S6.1: NOT Gate cotransformation. 23

Fig. S6.2: Arabinose-induced sgRNA expression in bacteria to repress GFP by CRISPRi. 24

Fig. S6.3: IPTG-induced sgRNA expression in bacteria to repress GFP by CRISPRi. 25

Fig. S6.4: Phage-delivered sgRNA-1 repressed GFP by CRISPRi. 26

Fig. S6.5: Infection over time of receiver cells grown with sgRNA-1 sender cells. 27

Fig. S6.6: Transfer frequency calculated for sgRNA-1 sender-to-receiver cells without antibiotics. 28

Fig. S6.7: Infection over time of receiver cells grown with sgRNA-A sender cells. 29

Fig. S6.8: Infection over time of receiver cells grown with sgRNA-C sender cells. 30

Fig. S6.9: Time to repression of 50% receivers when co-cultured with 3 different i-CRISPRi sender variants. 31

Note S7: NOT- & YES- gate implementation in bacterial cocultures 32

Fig. S7.1: NOT Gate: fluorescence over time (post-selection). 32

Fig. S7.2: NOT Gate: mean fluorescence of receiver cells at 16th hour (post-selection). 33

Fig. S7.3: YES Gate: fluorescence over time (post-selection). 34

Fig. S7.4: YES Gate: mean fluorescence of receiver cells at 16th hour (post-selection). 35

Fig. S7.5: YES Gate: mean fluorescence of receiver cells at 4th-hour (pre-selection). 36

Note S8: AND- & AND-AND-NOT- gate implementation in bacterial cocultures 37

Fig. S8.1: AND Gate: fluorescence over time (post-selection). 37

Fig. S8.2: AND Gate: mean fluorescence of receiver cells at 16th hour (post-selection). 38

Fig. S8.3: AND-AND-NOT Gate: fluorescence over time (post-selection). 39

Fig. S8.4: NOT Gate: mean fluorescence of receiver cells at 16th hour (post-selection). 40

Note S9: 20% activation time of ‘ON-state’ receiver cells. 41

Fig. S9.1: 20% activation time for single- & multi- input receivers. 41

Note S10: Screening AND-AND-NOT circuit design variants. 42

Fig. S10.1: Strain variants designed to implement the AND-AND-NOT gate in receiver cells. 44

Fig. S10.2: AND-AND-NOT gate co-transformations. 45

Fig. S10.3: AND-AND-NOT gate variants: sender to receiver communication. 46

**Supplementary Tables 47**

Table S1: Transformed Strains 47

Table S2: Plasmids 49

Table S3: Core Strains 51

**References 52**

### Supplementary Notes

#### Note S1: PFU and CFU assays

Bacteriophage counting methods estimate the number of phages by directly or indirectly counting infectious virions (phage particles) or their constituent nucleic acids or proteins (1, 2). Depending on the counting method used, different phage counts may be obtained from the same phage preparations. In this section, we discuss how this is relevant for estimating the phage titres from our -gp3φ and +gp3φ phage preps.

***PFU (plaque forming unit) assay:*** If a phage can productively infect cells, releasing its progeny after the primary infection, a PFU assay can be used to count the number of viable phages. As growing cells immobilised in the soft agar encounter phages, one infected cell releases (by lysis or secretion) several progeny phages that infect the neighbouring cells to form a visible zone of clearance (due to lysis by lytic phages) or low turbidity (due to reduced growth of cells infected by filamentous phages). The +gp3*φ* phages used in this work are capable of forming plaques, provided the receiver cells have the F-pilus receptor for entry (encoded by the F-plasmid) and the rest of the M13 phage machinery for packaging and re-secretion (encoded by the Helper plasmid). Since receivers carrying the full Helper (with *gp3* gene) are immune to M13 infection (**Fig.** S3.1b), due to superinfection inhibition by gp3 (3), we use a Δgp3 Helper plasmid for PFU assay with +gp3*φ* phages.

During the PFU assay, there is no pre-incubation between the phages and the cells; the two are mixed together in soft agar and quickly poured onto the plate. As receiver cells continue to multiply in the soft agar they form a full lawn, except in places where their growth is retarded due to infection by M13 phage. The plaques observed at the end of the overnight incubation contain not just initially infected receiver cells but also their neighbouring cells subsequently infected by newly secreted phages. As a result, the physiological state of the receiver cells at the time of mixing with phages has little impact on the PFU counts (**Fig.** S1.1). While this means that PFU counts of phage numbers are more accurate than CFU counts (see below), an infection event occurring very late in the incubation period may still be missed in counting because there is not enough time for the plaque to grow to a visible size before cell growth stops due to nutrient or space exhaustion in the soft agar.

***CFU (colony forming unit) assay:*** If a phage can infect cells but cannot produce progeny phages, a PFU assay is not possible. However, the infection events can still be counted by a CFU assay that involves spreading the infected cells on a selective antibiotic plate to count infected cells that form resistant colonies (4). A typical CFU assay relies on the assumption that at low multiplicity of infection (MOI << 1, $10^{-3}$ to $10^{-6}$), since the number of receiver cells is in vast excess of the number of phages, most phages will infect a cell within 20 minutes of incubation and express the antibiotic resistance gene. However, when comparing the CFUs and PFUs obtained at different receiver densities and physiology from the same phage prep (**Fig.** S1.1 and S1.2), we find that the CFU method underestimates phage counts by several fold. This is because the CFU method only counts the infections that occur in the first 20 minutes of pre-selection incubation period (or shortly afterwards). Given the slow adsorption of M13 phages (5), a much longer time would be needed for infection by 100% phages even at low MOI.

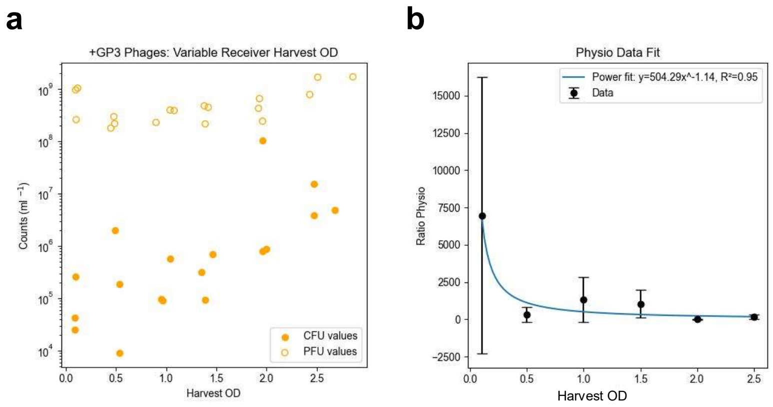


**Fig. S1.1: Receivers at different physiologies (harvest $OD_{600}$) infected with the same concentration of phages.**

**(a)** Filled circles represent CFU values for +gp3φ phages with ER2738F as receivers (data same as in Fig. 1j). Hollow circles represent PFU values for +gp3φ phages with ER2738F_HelperΔgp3 as receivers.
**(b)** PFU/CFU ratios were obtained for corresponding physiologies (harvest $OD_{600}$) using the +gp3φ data in (a). The fitted curve represents the relationship between PFU and CFU numbers measured for the same phage concentration, when estimated using receiver cells harvested at different $OD_{600}$. PFU/CFU ratio = 504.29*(harvest $OD_{600}$)^-1.14

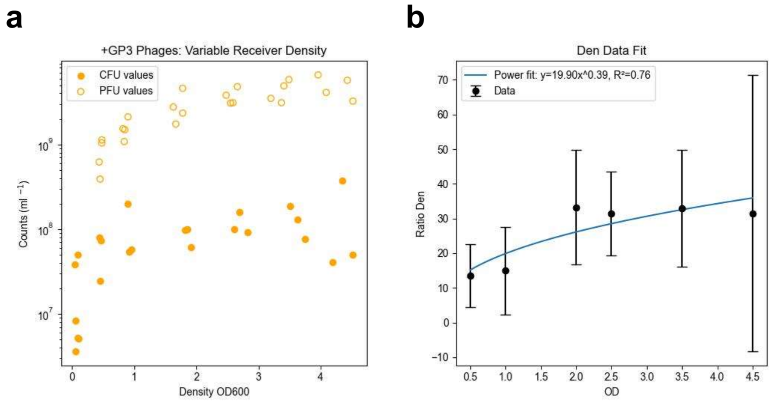


**Fig. S1.2: Receivers at different densities (resuspended $OD_{600}$) infected with the same concentration of phages.**

**(a)** Filled circles represent CFU values for +gp3φ phages with ER2738F as receivers. Hollow circles represent PFU values for +gp3φ phages with ER2738F_HΔgIII as receivers.
**(b)** PFU/CFU ratios were obtained for corresponding densities (resuspended $OD_{600}$) using the +gp3φ data in (a). The fitted curve represents the relationship between PFU and CFU numbers measured for the same phage concentration, when estimated using receiver cells harvested at $OD_{600}$ ~1 and resuspended to different $OD_{600}$ values. PFU/CFU ratio = 19.90*(resuspended $OD_{600}$)^0.039

***PFU/CFU ratio:*** Although -gp3φ phages can only be assayed by CFU assay, the +gp3*φ* phages can be assayed by both PFU and CFU assays. We compared CFU and PFU titres obtained from the same +gp3*φ* phage prep and established a relationship between the numbers, for different receiver physiologies (**Fig.** S1.1 a&b) as well as different receiver densities (**Fig.** S1.2 a&b). These PFU/CFU relationships (**Fig.** S1.1b, S1.2b) show how much the choice of the counting method can underestimate the true phage counts in a sample. We determine an empirical relationship between CFU and PFU counts obtained using our protocol for the same phage prep (**Fig.** S1.3). We find that CFU counts can be converted to PFU counts by multiplying with a factor of 14.7.

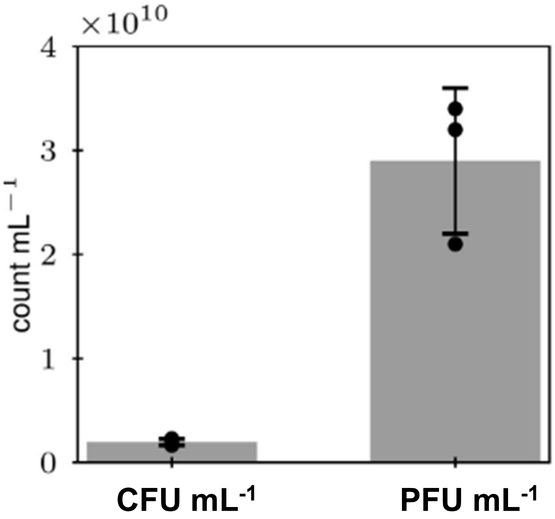


**Fig. S1.3: Relationship between CFU and PFU counts.**

Concentration of a +gp3φ phage stock was determined using CFU and PFU assays as 19.7x$10^8$ CFU $mL^{-1}$ and 29x$10^9$ PFU $mL^{-1}$, respectively, resulting in a conversion factor of 14.7-fold.

#### Note S2: Converting optical density to cell density

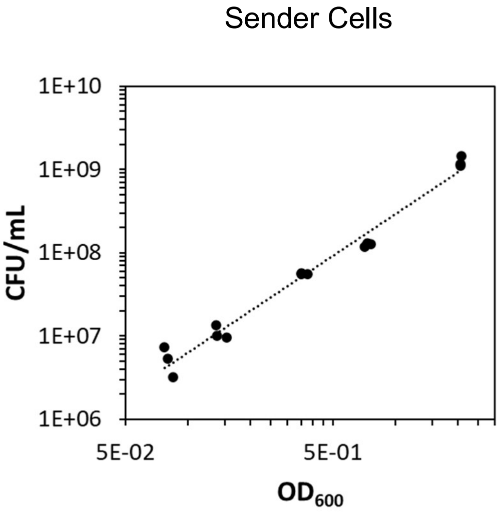


**Fig. S2.1: Conversion of $OD_{600}$ to cell density.**

Sender cell (TOP10) densities plotted against spectrophotometer $OD_{600}$ with a power fit.

Spectrophotometric measurements of optical density ($OD_{600}$) are often used in microbiology as an indirect measure of cell density in a liquid culture. Following from the Beer-Lambert law, a linear fit is often used to convert $OD_{600}$ to cell density (6). If cells in the population are uniform in size, this would be a correct approximation (7). However, the sizes of cells are not the same through the duration of a batch culture. They grow when going from the lag phase to the exponential phase, and then shrink back in the stationary phase (8). Therefore, non-linear relationships between OD and CFU are better estimates of cell numbers when their sizes vary between ODs (9).

Here, we established the relationship between the absorbance ($OD_{600}$) and cell density for the sender strain: TOP10. This was done by diluting by 1000x an overnight culture and re-growing at 37 °C in fresh LB media. Aliquots were taken from the growing cultures at several time-points and their absorbance ($OD_{600}$, spectrophotometer UVisco V-1100D) recorded, cooled down on ice for ~30 min, following which the $OD_{600}$ was measured again. The chilled cultures were serially diluted, $10^{-4}$ to $10^{-7}$ dilutions were spread on LBA plates, and the colony forming units (CFU) obtained were counted the next day. A power fit (**Fig.** S2.1) describes the relationship more accurately than a linear fit or a quadratic fit (not shown). The power function fit to go from spectrophotometer $OD_{600}$ to cell concentration (CFU $mL^{-1}$) for sender cells is:

*Cell density = factor * ($OD_{600}$ ^ exponent)*

Senders (TOP10)
CFU = 293192345 * ($OD_{600}$ ^ 1.67)

#### Note S3: gp3 causes infection immunity

Due to the superinfection immunity caused by the expression of the M13 minor coat protein gp3 (3), F+ receiver cells expressing it do not get infected by the M13 phage.

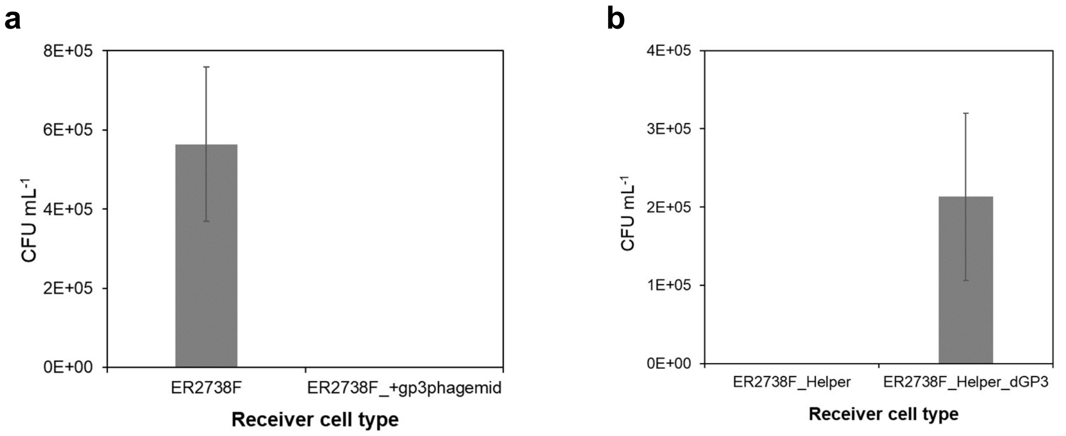


**Fig. S3.1: Expression of gp3 in receiver cells inhibits phage infection.**

Infection of four receiver strains by the same concentration of a -gp3φ phage (pSEVA19_sgRNA-C0_M13ps, *AmpR*) and subsequent CFU counts (N=3). Only the receiver strains without gp3 expression show CFUs upon infection by the -gp3*φ* phage.

(a) CFU counts obtained using the standard receiver strain (left, ER2738F), and the same receiver strain already transformed with a +gp3φ phagemid (right, ER2738F_+gp3φphagemid)

(b) CFU counts obtained using the receiver strains with the full Helper plasmid that expresses the gp3 gene (left, ER2738F_H), and the Δgp3 Helper plasmid lacking the gp3 gene (right, ER2738F_HΔgIII)

Next, we tested if multiple -gp3φ phages secreted by different senders can simultaneously infect a single receiver cell. With selection for each additional phagemid, the growth of the receivers is negatively affected. In **Fig.** S3.2, we show the $OD_{600}$ of infected receiver cells at t=18h with (a) single, (b) double and (c) triple infections. Each phage type carries a different antibiotic resistance gene that allows antibiotic selection of infected receivers.

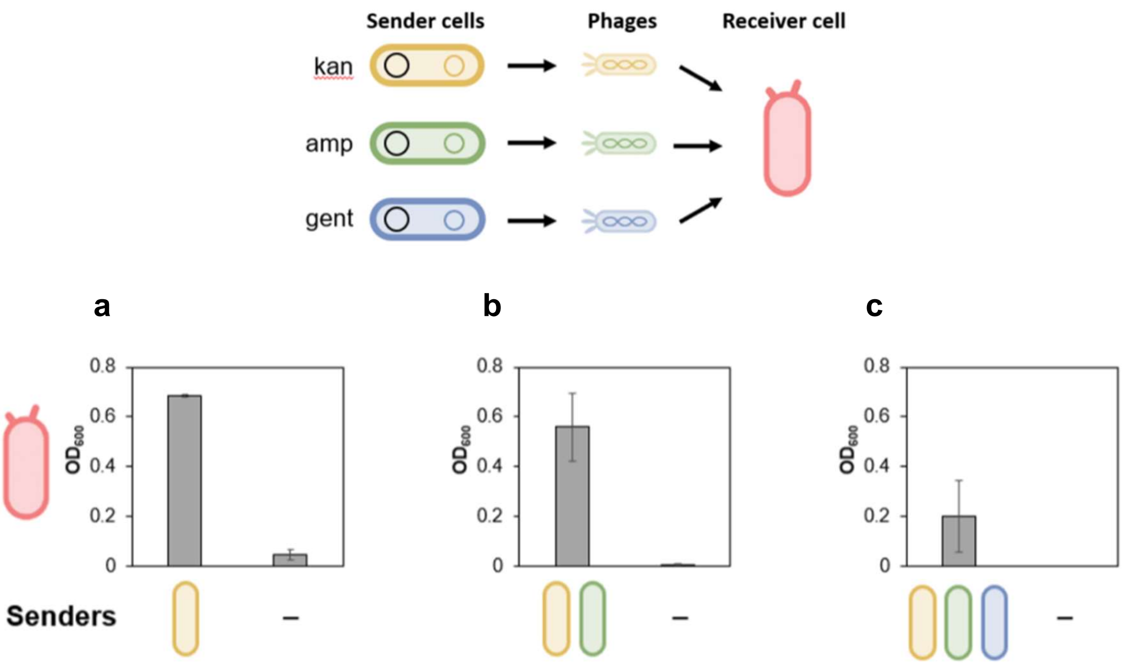


**Fig. S3.2: Multiple and simultaneous infection of receiver cells.**

Growth of receiver cells infected by (a) one, (b) two & (c) three phages secreted by -gp3*φ* sender cells. Phages carry different antibiotic-resistance genes. Infected cells were selected on one, two, or three antibiotics.

#### Note S4: Phage infection in growing bacteria

This Supplementary Note presents additional data related to the experiments shown in Figure 2.

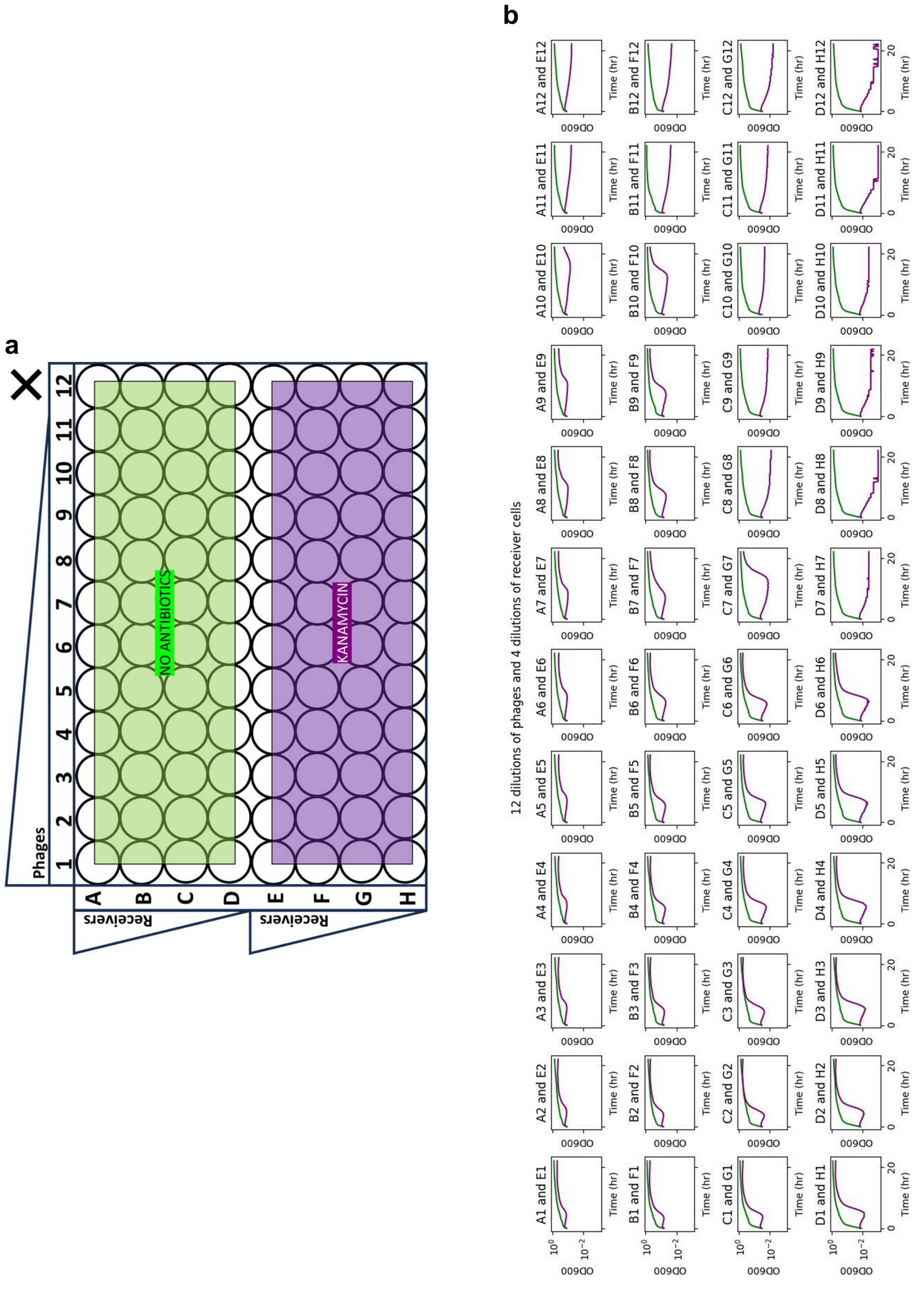


**Fig. S4.1: Growth curves of receiver cells infected with 12 different concentrations of phages.** (a) Plate map and colour code. (b) In a 96-well plate, receiver cells (starting $OD_{600}$ = 0.250, 0.125, 0.062, 0.031) were incubated with varying amounts of phages (dilutions: 1, 3^1, 3^2, … 3^10, 0 = no-phage) for 20 mins and then grown in a plate-reader with (purple) and without (green) antibiotics (N=1).

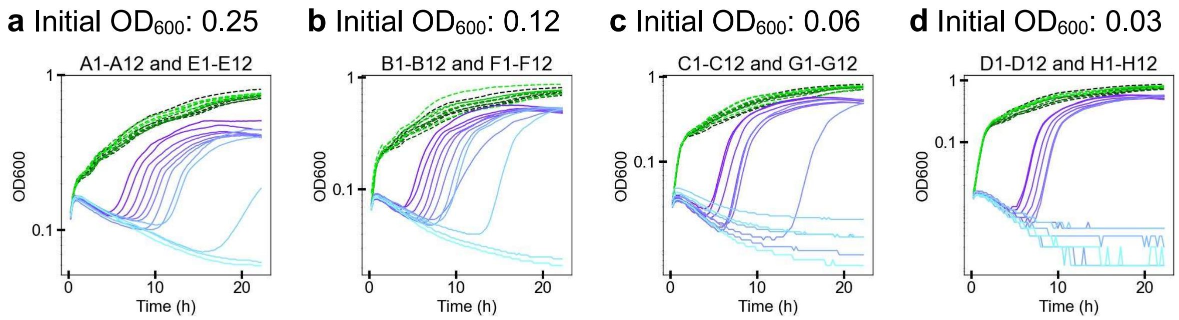


**Fig. S4.2: Phage dose-dependent growth rescue of receiver culture with different starting $OD_{600}$.**

Growth curves in dark blue & dark green come from receiver cultures incubated with the most amount of phages (undiluted phage prep), and light blue & light green growth curves come from cultures incubated with the least amount of phages (dilution: 3^10 or 0). Growth curves are for receiver cells at densities ($OD_{600}$): (a) 0.25 (b) 0.125 (c) 0.06 (d) 0.03. Growth curves in blue were grown with antibiotic selection for infected receivers (kan+tet). Well labels are the same as used in Fig. S4.1a.

**a**

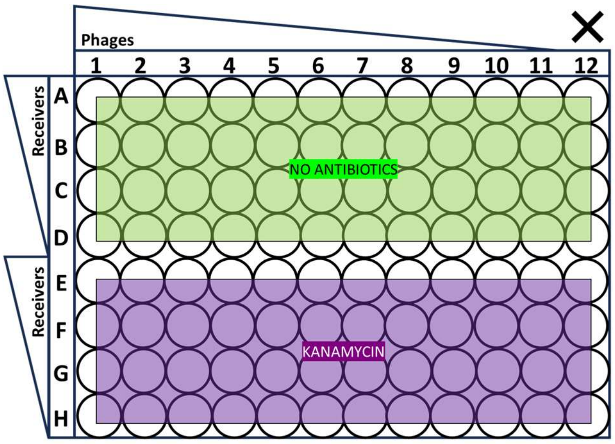


**b**

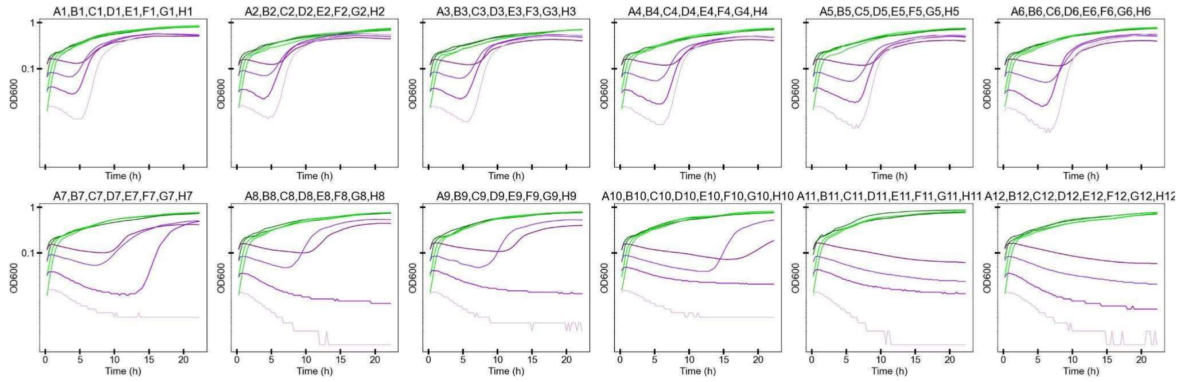


**Fig. S4.3: Receiver dose-dependent effect on growth rescue.**

(a) Plate map and colour code. (b) Each subplot has growth curves that come from receiver cultures incubated with the same amount of phages. Dark purple & dark green growth curves come from receiver cultures that have the most number of starting receiver cells. Light purple & light green have the least number of starting receiver cells. Growth curves in purple were grown with antibiotic selection and growth curves in green were grown without antibiotics.

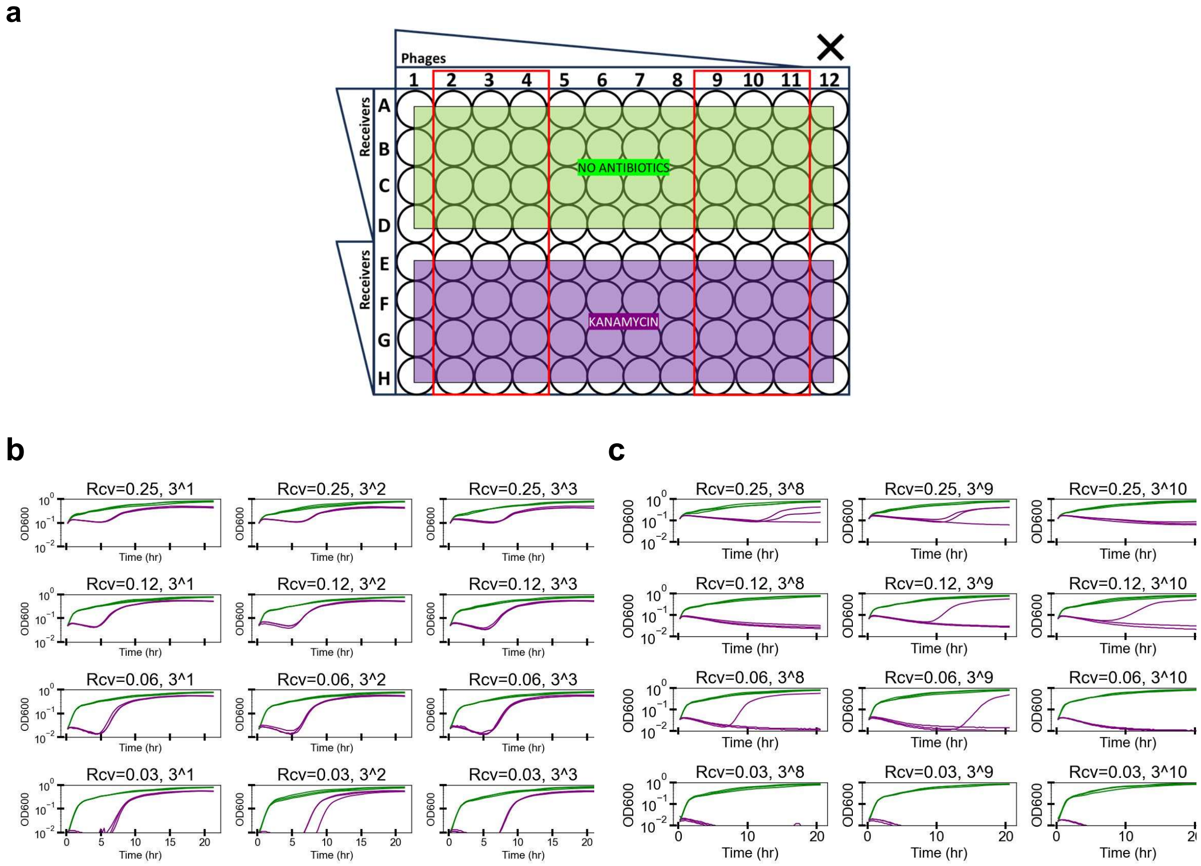


**Fig. S4.4: Phage-dose dependent growth rescue.**

(a) Plate map & colour code. Growth curves of receiver cultures of different starting $OD_{600}$, incubated with phages (carrying *KanR*) at different dilutions: (b) 3^1, 3^2, 3^3 and (c) 3^8, 3^9, 3^10. Infected receivers were selected using kanamycin in LB media (purple). Growth curves in green were the same cultures grown without antibiotic selection. N=3.

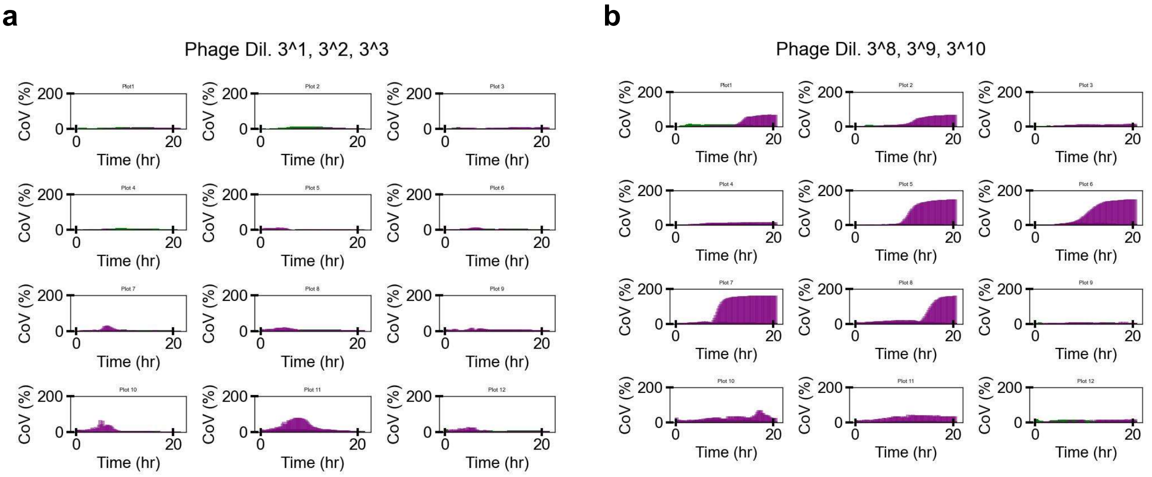


**Fig. S4.5: Coefficients of variation over time between repeats.**

CoV through time was analysed and plotted for growth curves presented in Fig. S4.4. Receiver cultures of different starting $OD_{600}$, incubated with phages (carrying *KanR*) of different dilutions: (a) 3^1, 3^2, 3^3 and (b) 3^8, 3^9, 3^10. CoVs are shown both for cultures grown without (green) or with (purple) kanamycin selection, although green plots are hard to see because of very low CoVs.

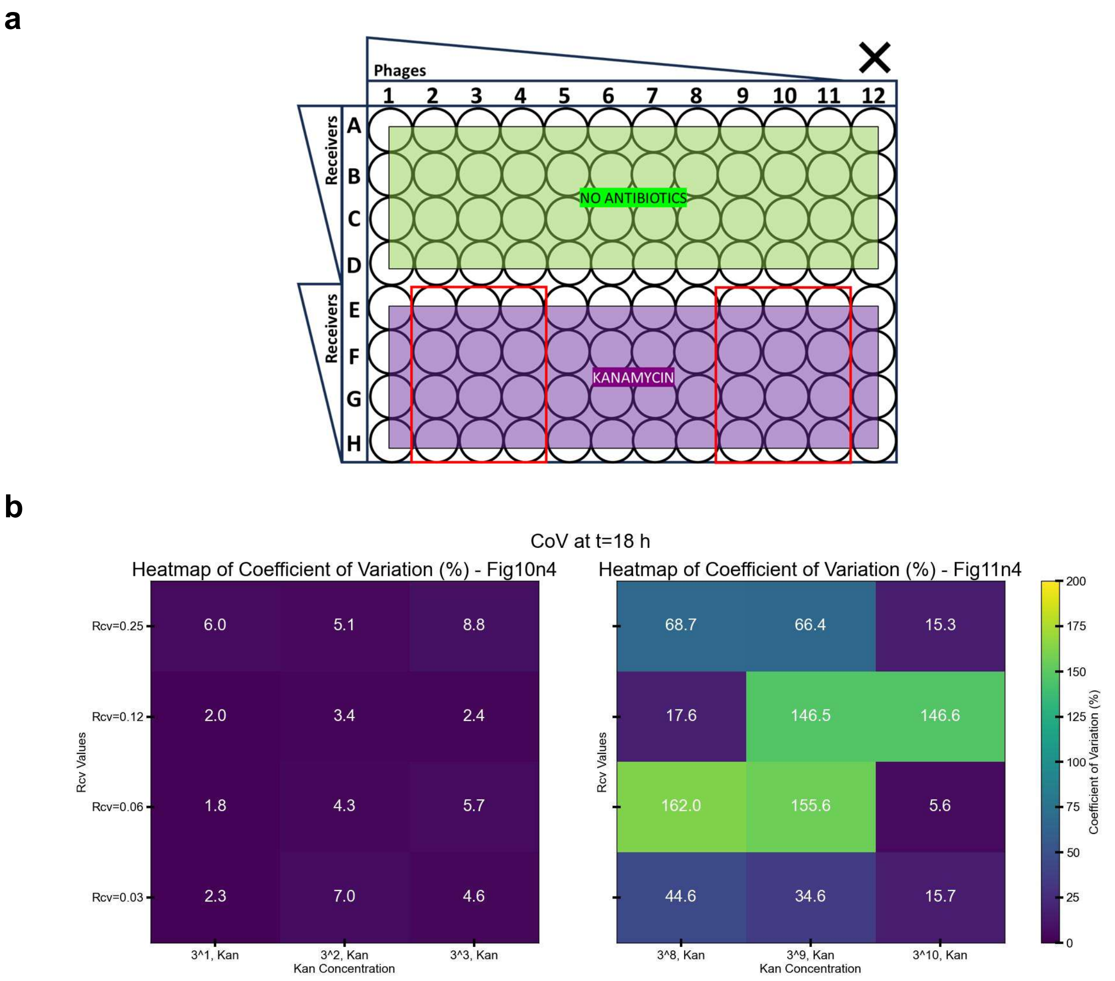


**Fig. S4.6: Heatmap of CoV of $OD_{600}$ at 18th hour from CoV over time shown in S4.5.**
(a) Plate-map. Red box represents the wells analysed here. (b) CoV% Heatmap at t=18 h.

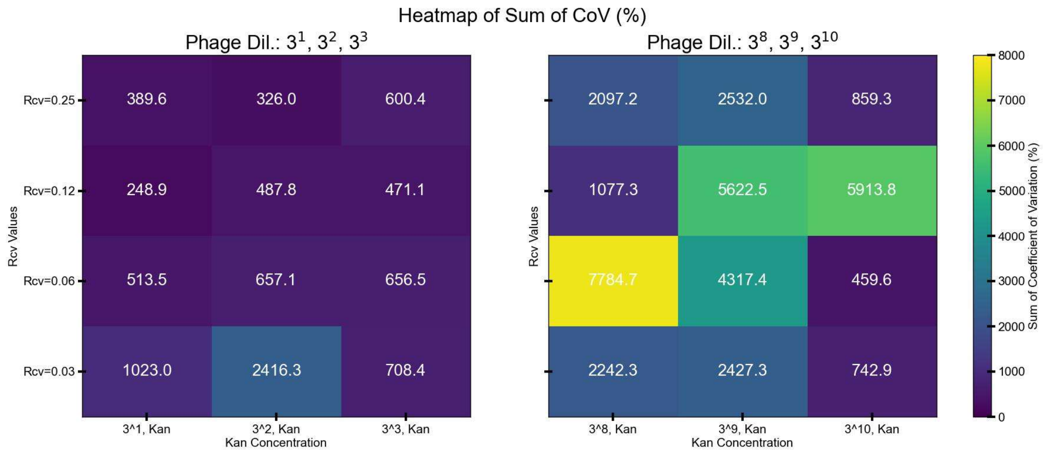


**Fig. S4.7: Heatmap of the sum of $OD_{600}$ CoVs through time (0-18 h).**
Data from wells marked by the red box in Fig. S4.6(a).

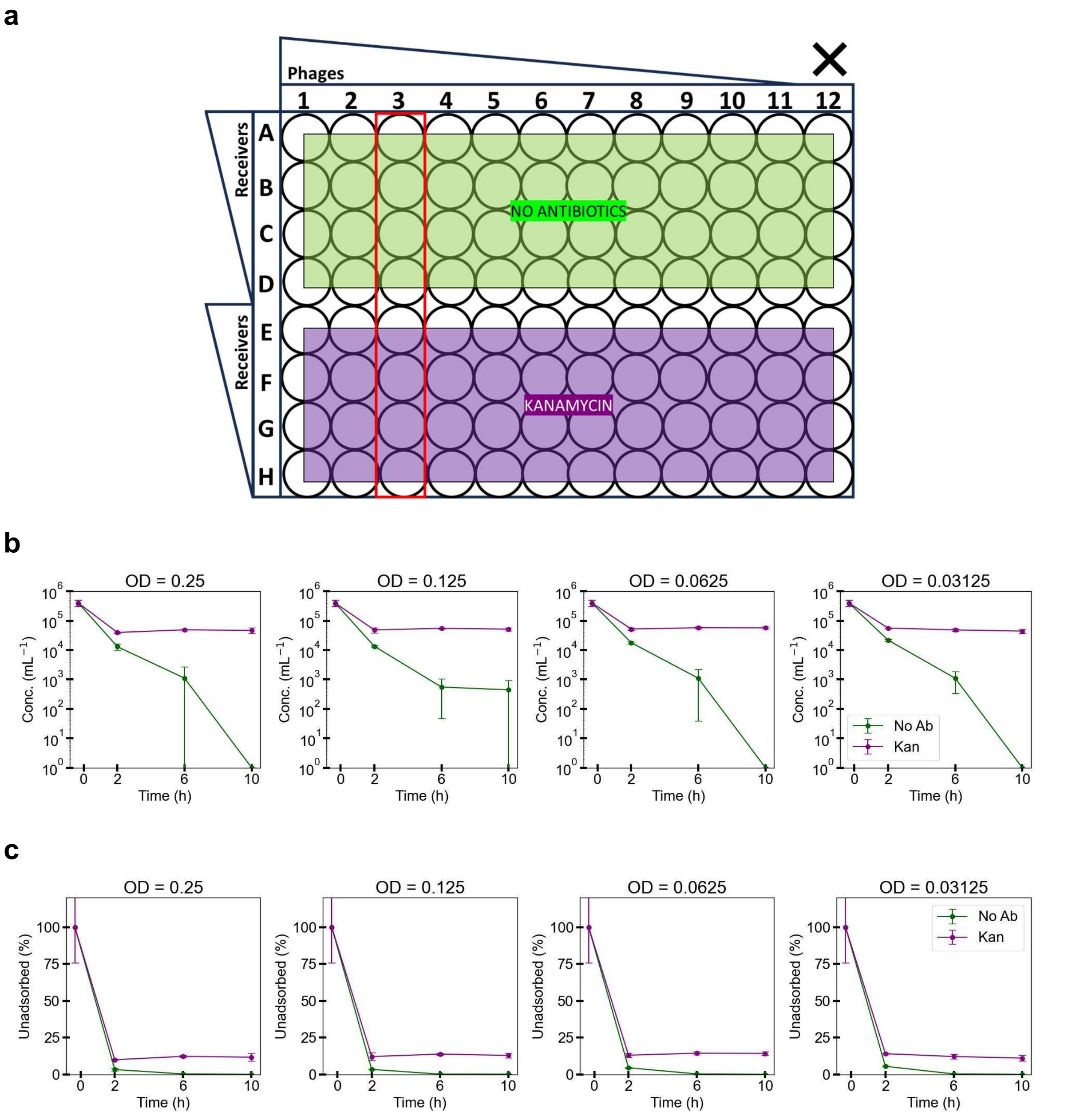


**Fig. S4.8: Phage adsorption by receiver cells over time.**

(a) Plate map and colour code. Red box indicates wells analysed. 45 µL of 4 x $10^5$ PFU/mL phages were mixed with different densities of 150 µL receiver cells (equivalent to $OD_{600}$ = 0.25, 1.25, 0.06 and 0.03; equivalent to wells A3–D3, E3–H3) and pre-incubated for 20 mins at RT. This was done in two sets. 1 set of cultures were grown with antibiotics (kanamycin) and another set was grown without antibiotics for 10 hours in a plate reader with shaking at 37°C. 3 µL culture samples were collected at t=2, 6, and 10 hours after the pre-incubation / plate reader was started. Unadsorbed phages were counted using PFU assay and shown as per mL concentration in (b), and as %unadsorbed in (c). Data points in green are unadsorbed phages from cultures grown without antibiotics, and purple data points from cultures grown with antibiotics.

#### Note S5: Cell-to-cell communication from sender to receiver cells

This Supplementary Note presents additional data related to the experiments shown in Figure 3.

##### -gp3φ *sender set*

**a**

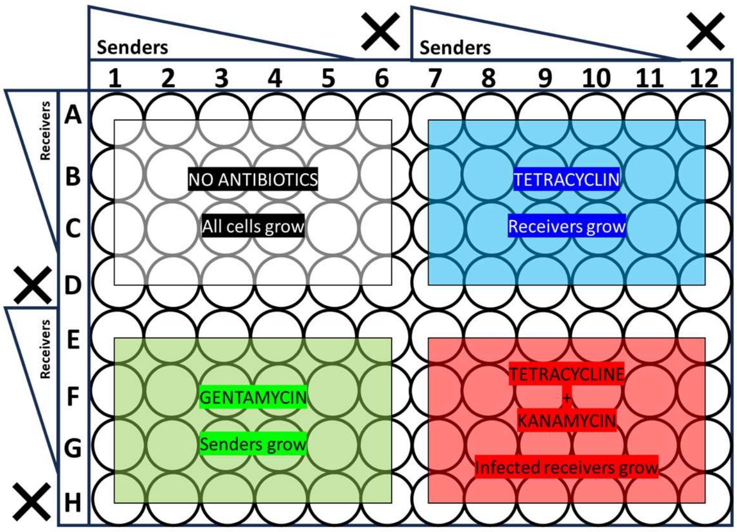


**b**

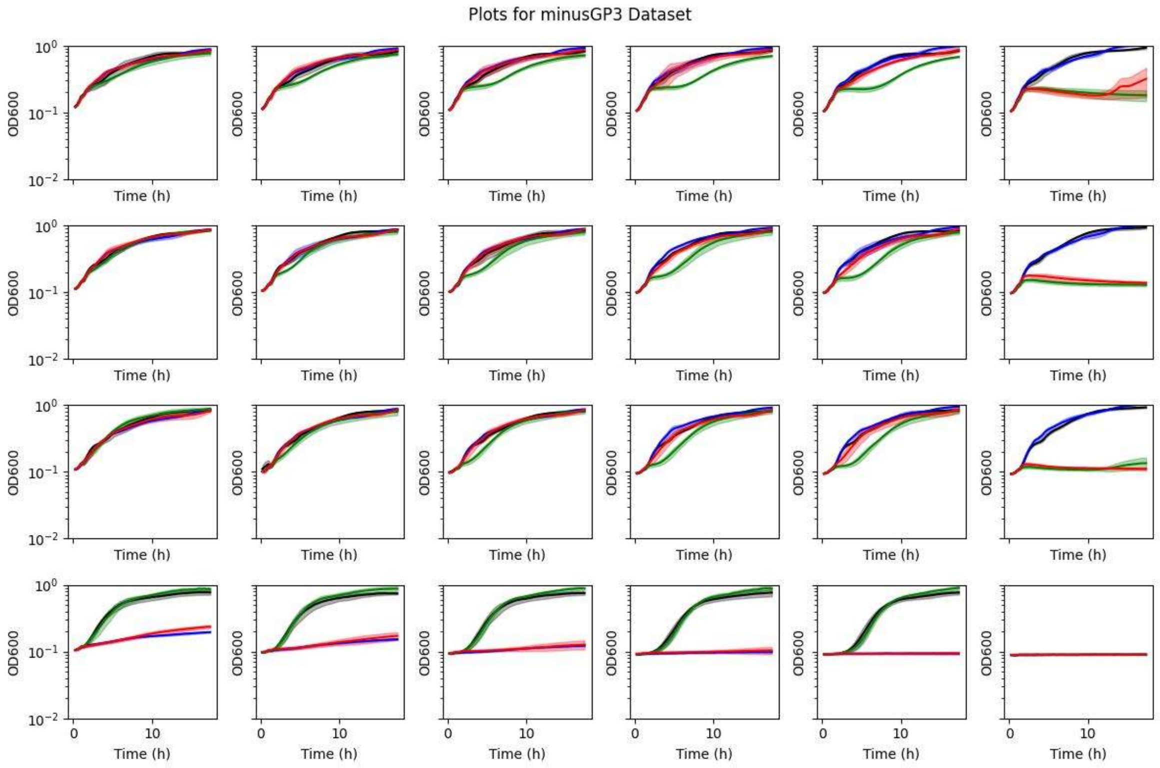


**Fig. S5.1: Cell-to-cell communication with -gp3φ senders & receivers.**

(a) Plate map and antibiotic selection colour code. (b) Growth curves: cell-to-cell communication with -gp3φ senders (N=3). Growth curves for cocultures with different initial densities of the receiver and sender cells. Cocultures were incubated for 1 hour without antibiotics, and antibiotic selection was subsequently applied for receivers (tet; blue), senders (gent; green), and infected receivers (kan+tet; red). Cocultures were grown without antibiotics to allow for the growth of all cell types (black).

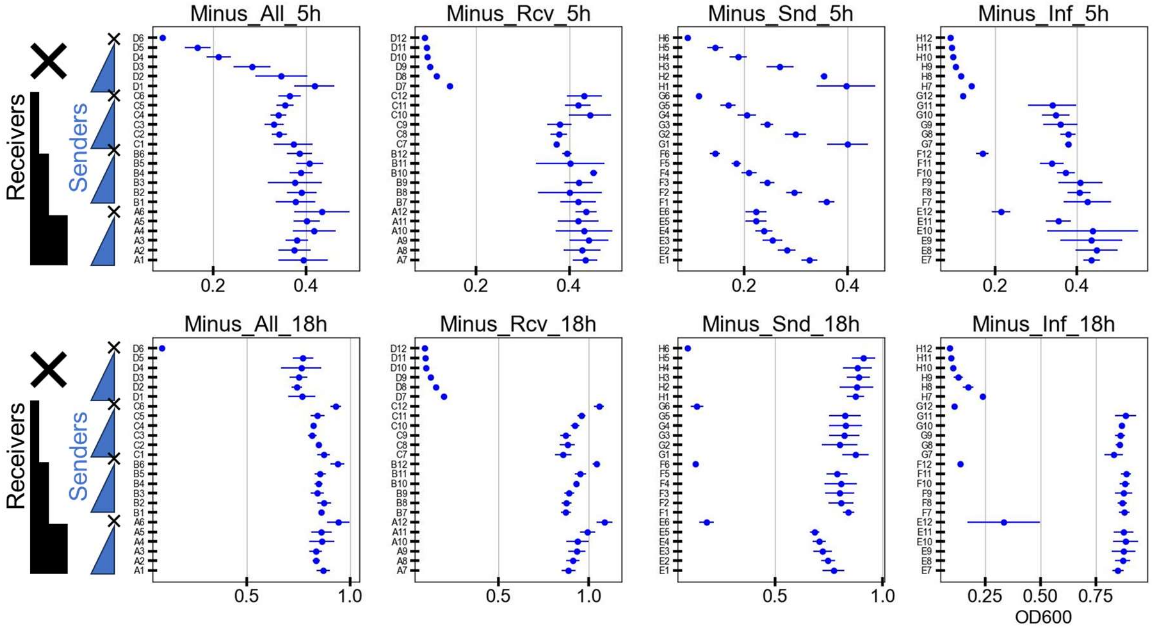


**Fig. S5.2: $OD_{600}$ at 5h and 18h, grouped by starting receiver densities (-gp3φ set).**

$OD_{600}$ at time t=5h (top row) and t=18h (bottom row) for different cultures selected for different cell types in communication experiments with -gp3φ senders. Senders and receivers of different initial densities were grown (N=3), as indicated in the plate map in Fig. S5.1a.

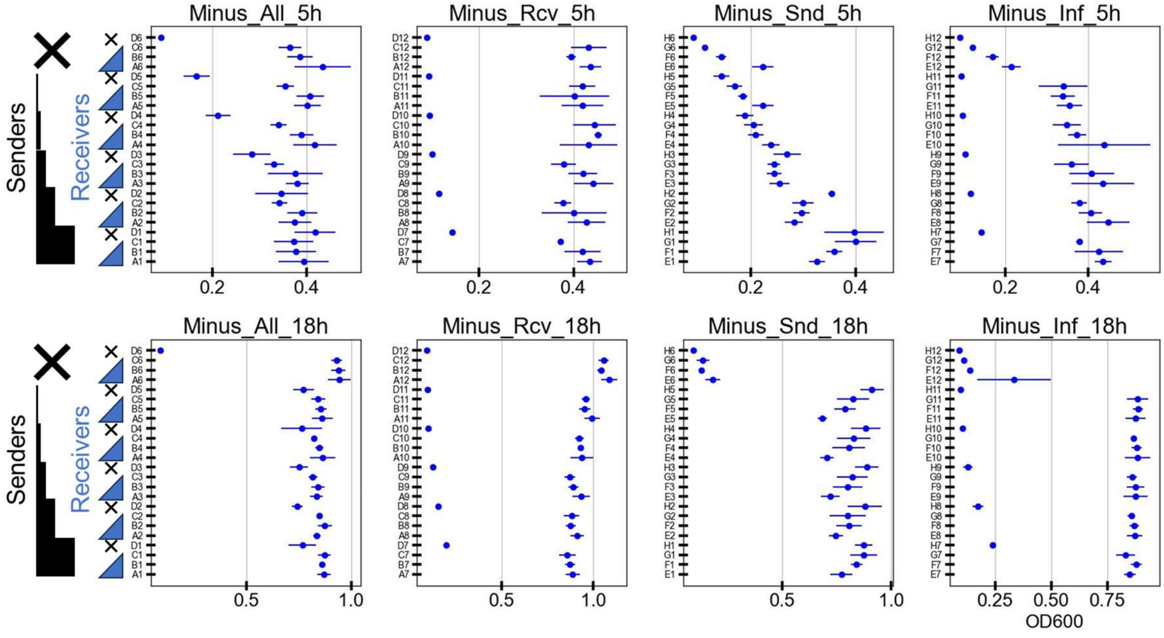


**Fig. S5.3: $OD_{600}$ at 5h and 18h, grouped by starting sender densities (-gp3φ set).**

$OD_{600}$ at time t=5h (top row) and t=18h (bottom row) for different cultures selected for different cell types in communication experiments with -gp3φ senders. Senders and receivers of different initial densities were grown (N=3), as indicated in the plate map in Fig. S5.1a.

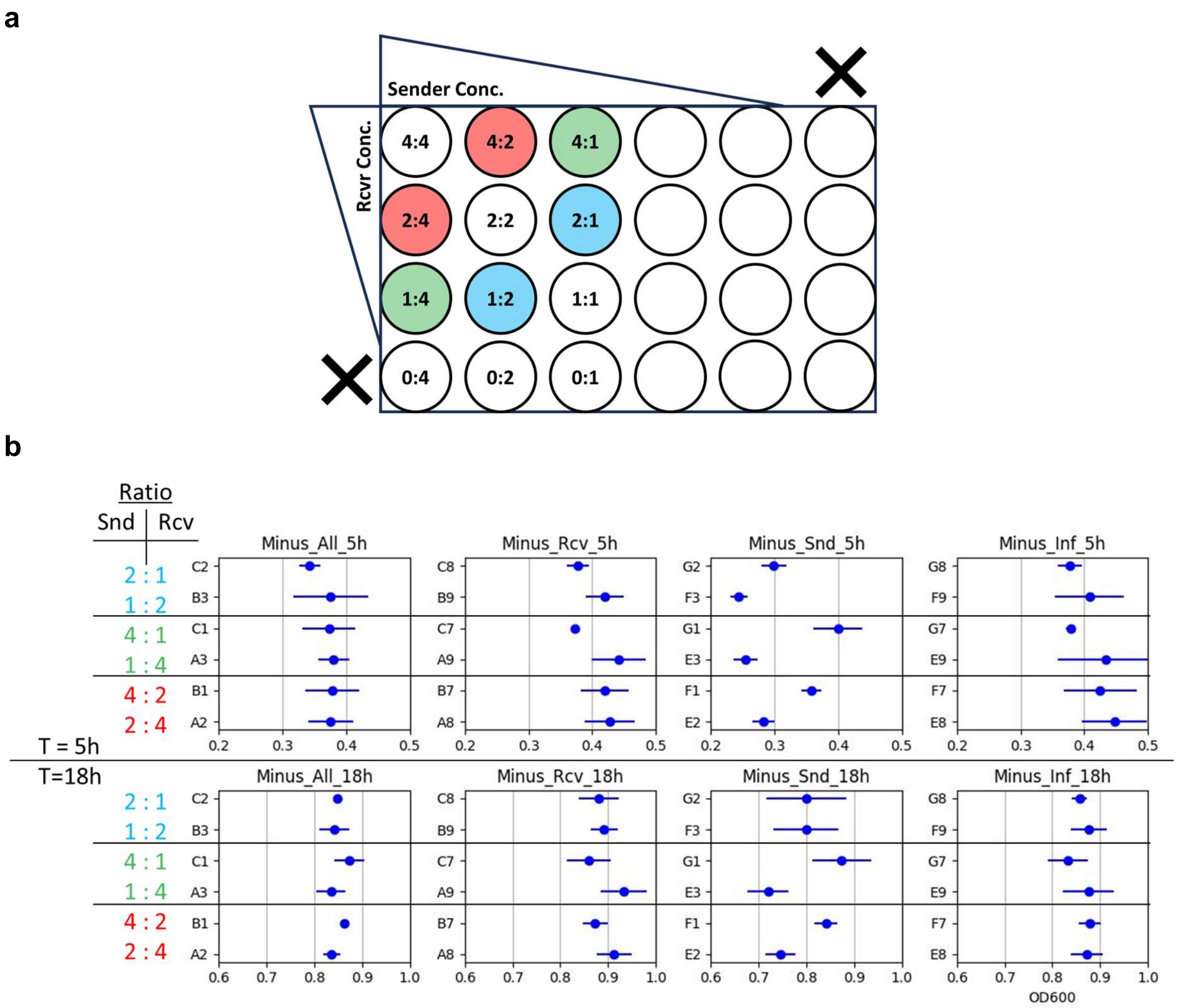


**Fig. S5.4: $OD_{600}$ at 5h and 18h, showing data for wells with the same total starting OD of senders and receivers (-gp3φ set).**

(a) Plate map of the wells in one quadrant, grouped together in three sets (red, green, blue), each pair of which have the same total (senders+receivers) starting OD. (b) A subset of the data from Fig. S5.2 / S5.3. $OD_{600}$ at time t=5h (top row) and t=18h (bottom row) for different cultures selected for different cell types in communication experiments with -gp3φ senders. Senders and receivers of different initial densities were grown (N=3), as indicated in the plate map in Fig. S5.1a.

##### +gp3*φ sender set*

**a**

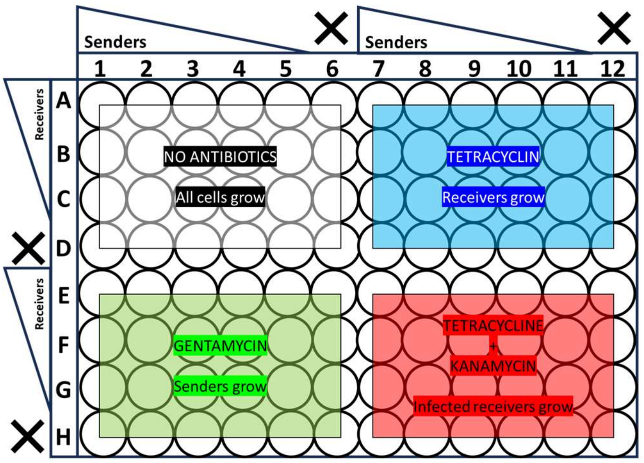


**b**

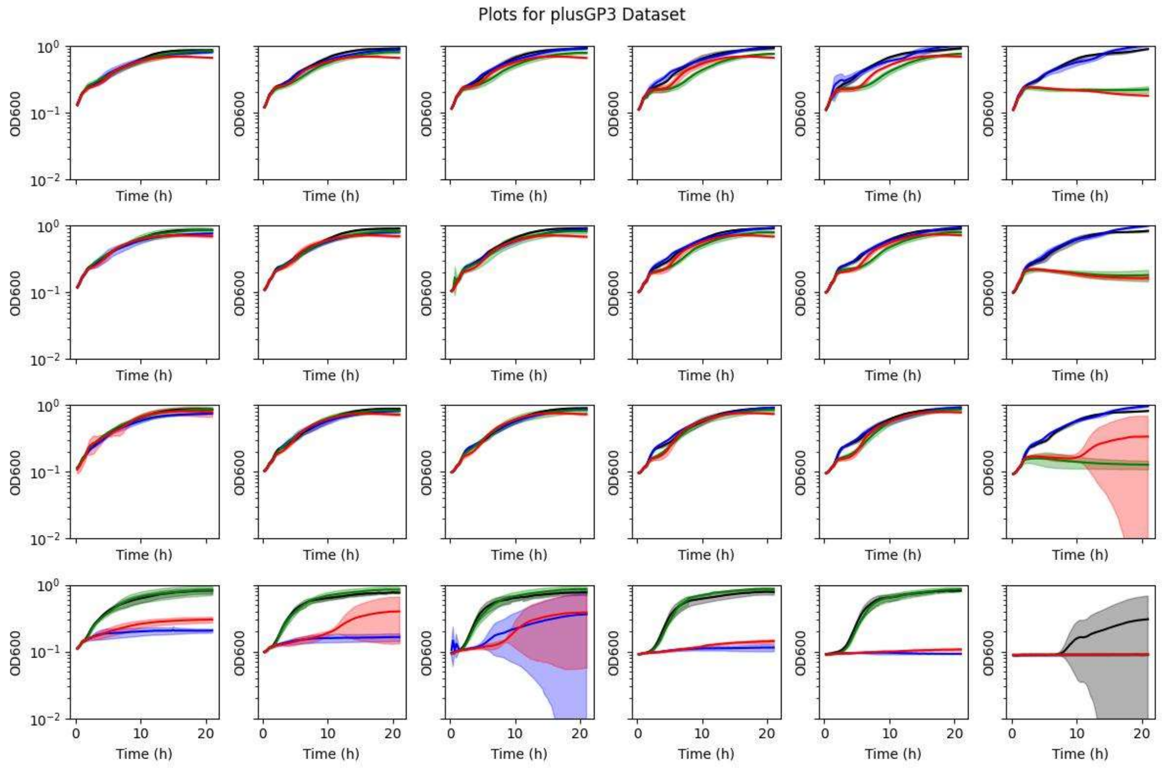


**Fig. S5.5: Cell-to-cell communication with +gp3φ senders & receivers.**

(a) Plate map and antibiotic selection colour code. (b) Growth Curves: cell-to-cell communication with +gp3φ senders (N=3). Growth curves for cocultures with different initial densities of the receiver and sender cells. Cocultures were incubated for 1 hour without antibiotics, and antibiotic selection was subsequently applied for receivers (tet; blue) senders (gent; green), and infected receivers (kan+tet; red). Cocultures were grown without antibiotics to allow for the growth of all cell types (black).

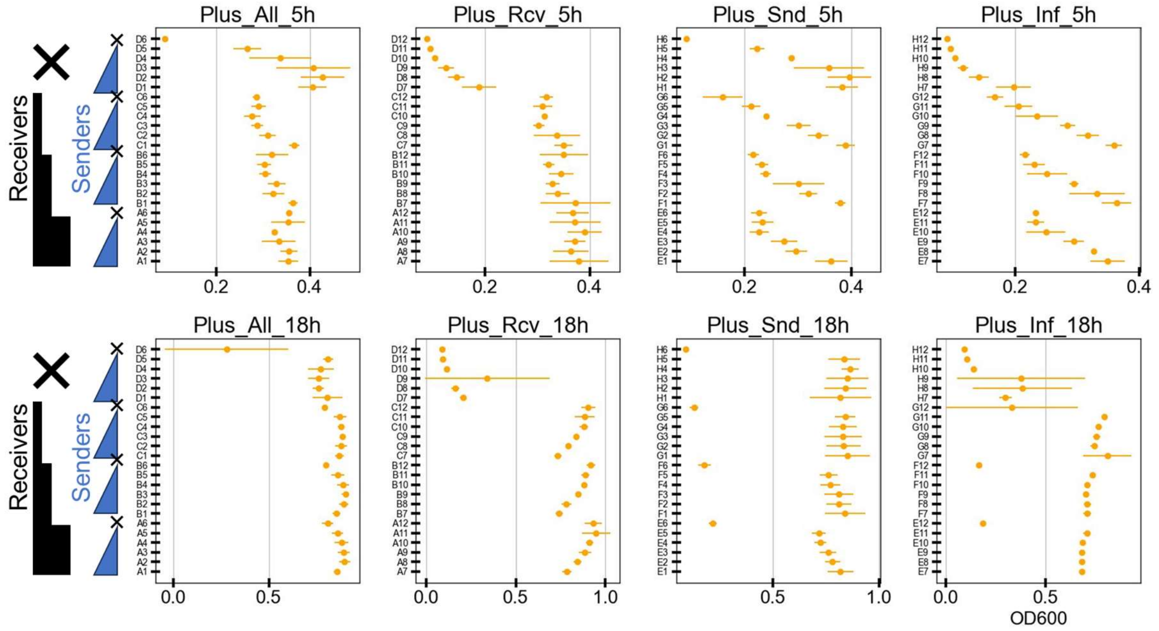


**Fig. S5.6: $OD_{600}$ at 5h and 18h, grouped by starting receiver densities (+gp3φ set).**

$OD_{600}$ at time t=5h (top row) and t=18h (bottom row) for different cultures selected for different cell types in communication experiments with +gp3φ senders. Senders and receivers of different initial densities were grown (N=3), as indicated in the plate map in Fig. S5.5a.

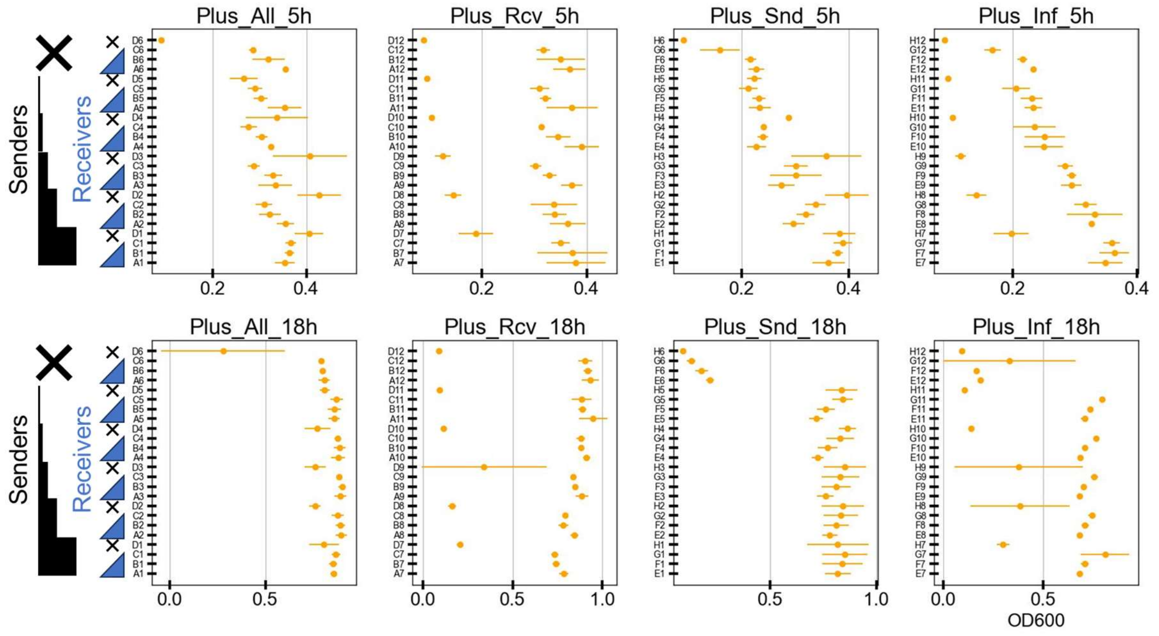


**Fig. S5.7: $OD_{600}$ at 5h and 18h, grouped by starting sender densities (+gp3φ set).**

$OD_{600}$ at time t=5h (top row) and t=18h (bottom row) for different cultures selected for different cell types in communication experiments with +gp3φ senders. Senders and receivers of different initial densities were grown (N=3), as indicated in the plate map in Fig. S5.5a.

**a**

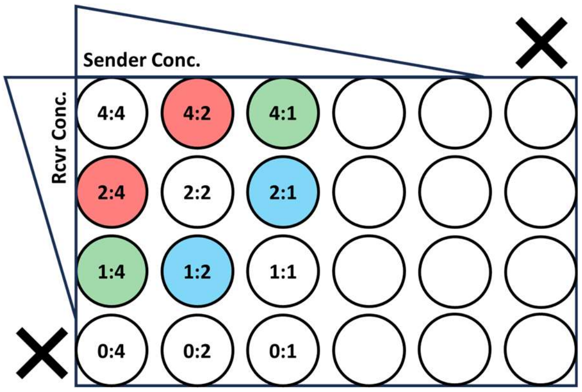


**b**

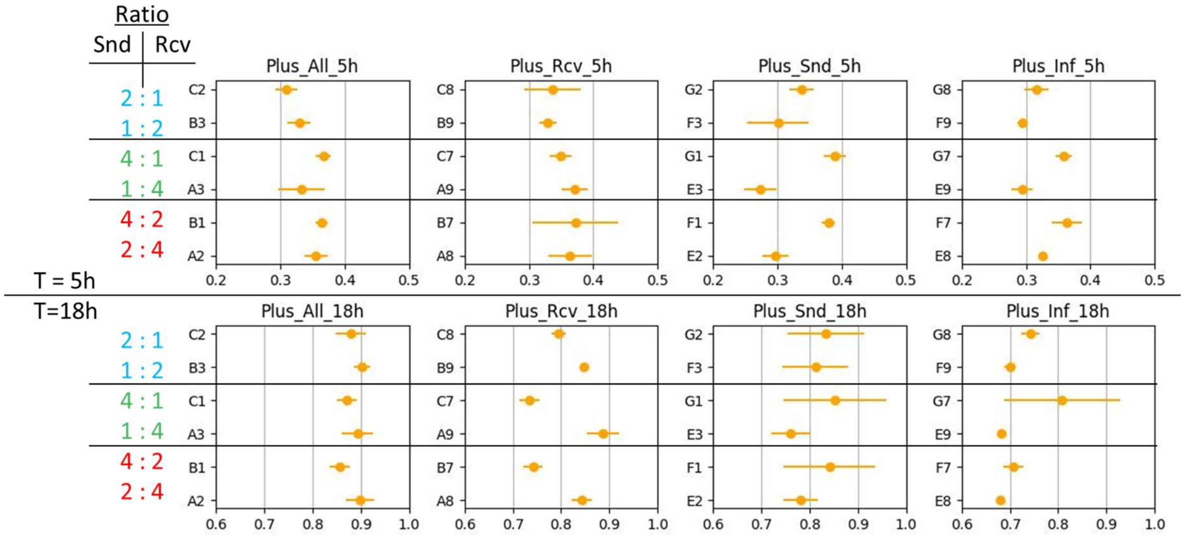


**Fig. S5.8: $OD_{600}$ at 5h and 18h, showing data for wells with the same total starting OD of senders and receivers (+gp3φ set).**

(a) Plate map of the wells in one quadrant, grouped together in three sets (red, green, blue), each pair of which have the same total (senders+receivers) starting OD. (b) A subset of the data from Fig. S5.6 / S5.7. $OD_{600}$ at time t=5h (top row) and t=18h (bottom row) for different cultures selected for different cell types in communication experiments with +gp3φ senders. Senders and receivers of different initial densities were grown (N=3), as indicated in the plate map in Fig. S5.5a.

a

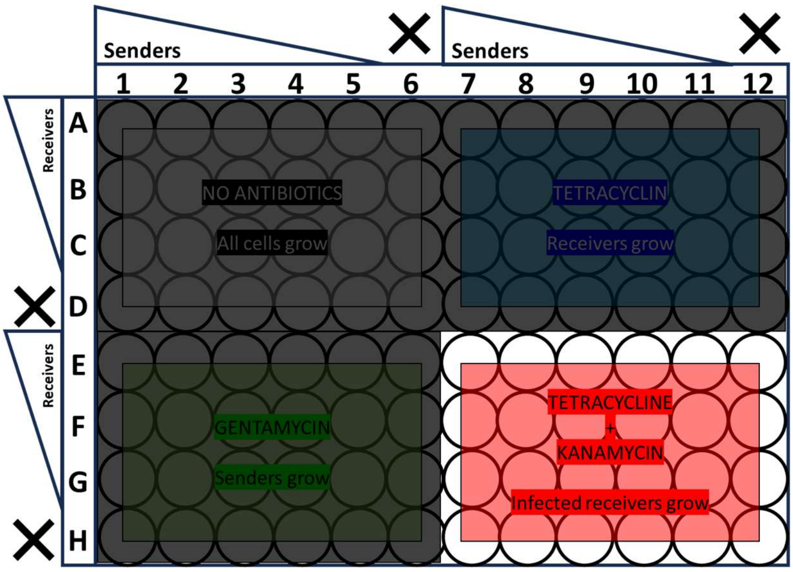


b

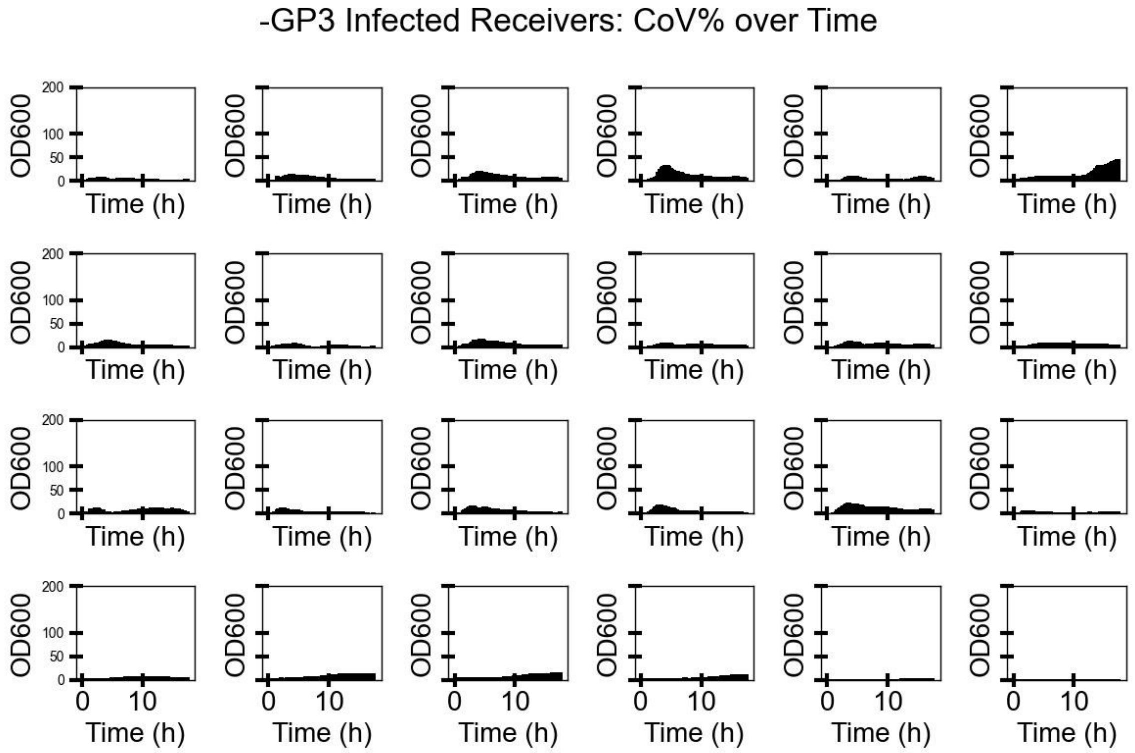


**Fig. S5.9: CoV% over Time calculated between 3 repeats for -gp3φ sender set.**

(a) The plate map highlights which wells are shown in (b). Data from Fig. S5.1.

**a**

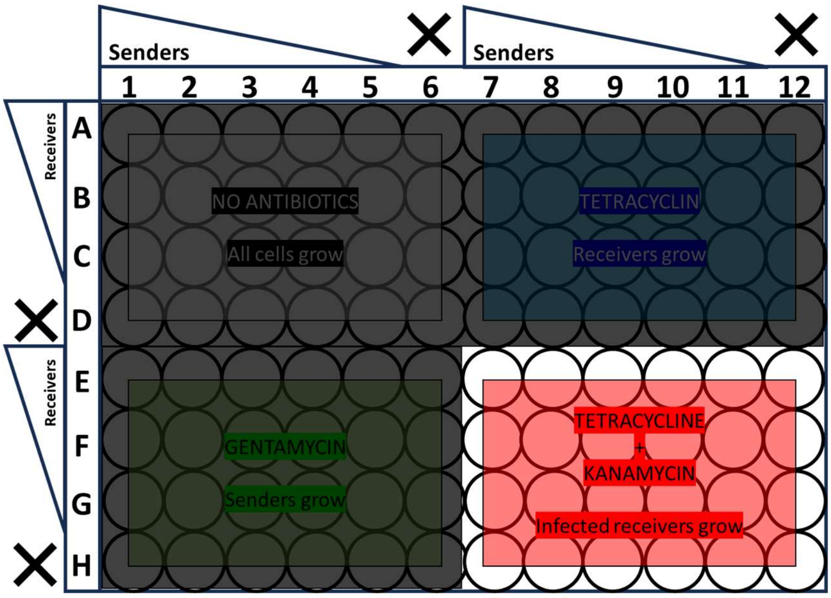


**b**

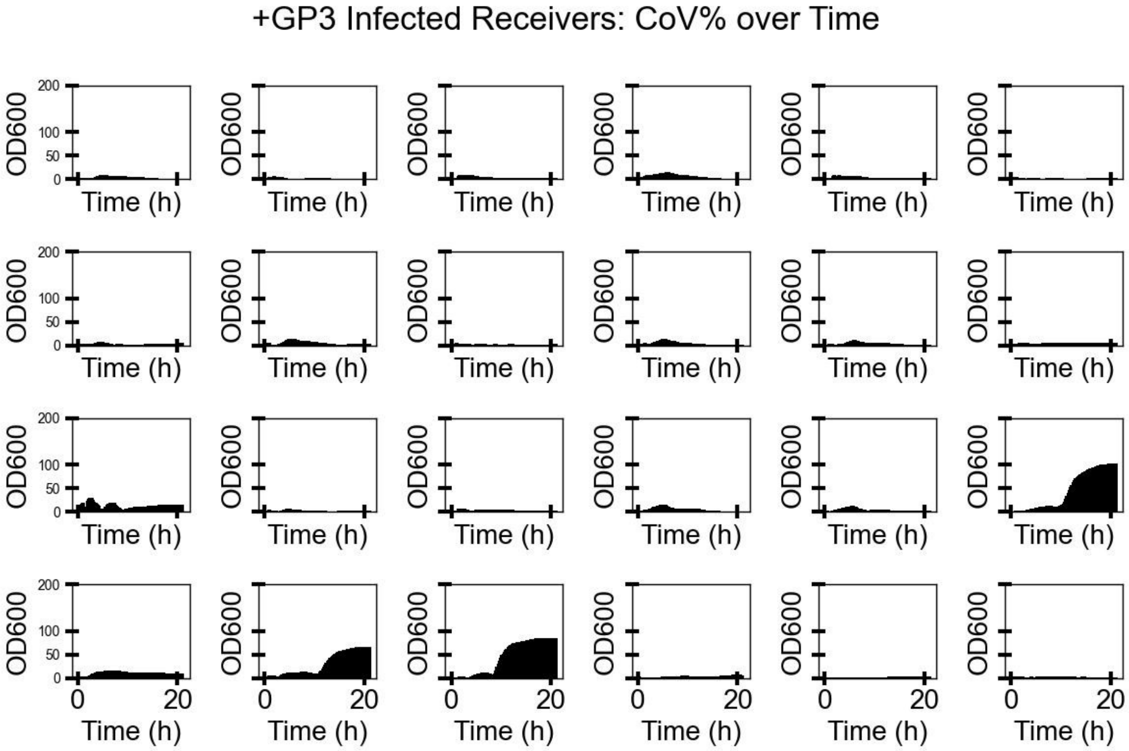


**Fig. S5.10: CoV% over Time calculated between 3 repeats for +gp3φ sender set.**
(a) The plate map highlights which wells are shown in (b). Data from Fig. S5.5.

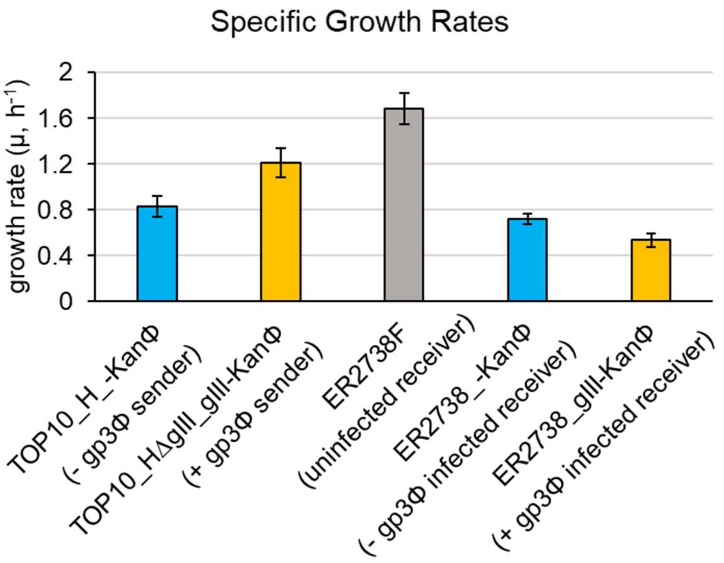


**Fig. S5.11: Specific growth rates of receiver, senders and infected receiver strains.**

Specific growth rates of -gp3φ sender and -gp3φ infected receiver are in blue bars, and +gp3φ senders and +gp3φ infected receivers are in orange bars. Growth rate of uninfected receiver bacteria is shown in grey.

#### Note S6: Using CRISPRi to track real-time receiver infection

This Supplementary Note presents additional data related to the experiments shown in Figure 4.

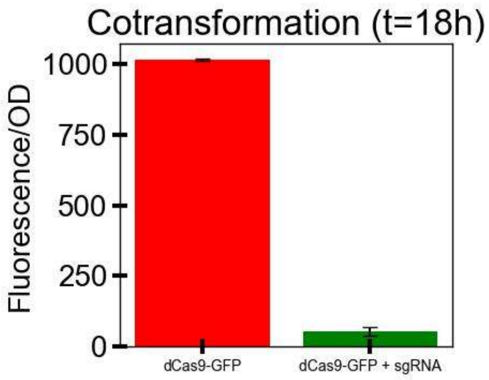


**Fig. S6.1: NOT Gate cotransformation.**

Normalised fluorescence of Top10F cells transformed with pSEVA47_dCas9-GFP_NOT_i1 (red bar), and Top10F cells transformed with pSEVA47_dCas9-GFP_NOT_i1 and pSEVA19_sgRNA-1_M13ps (green bar). (N=3, s.d. shown as error bars).

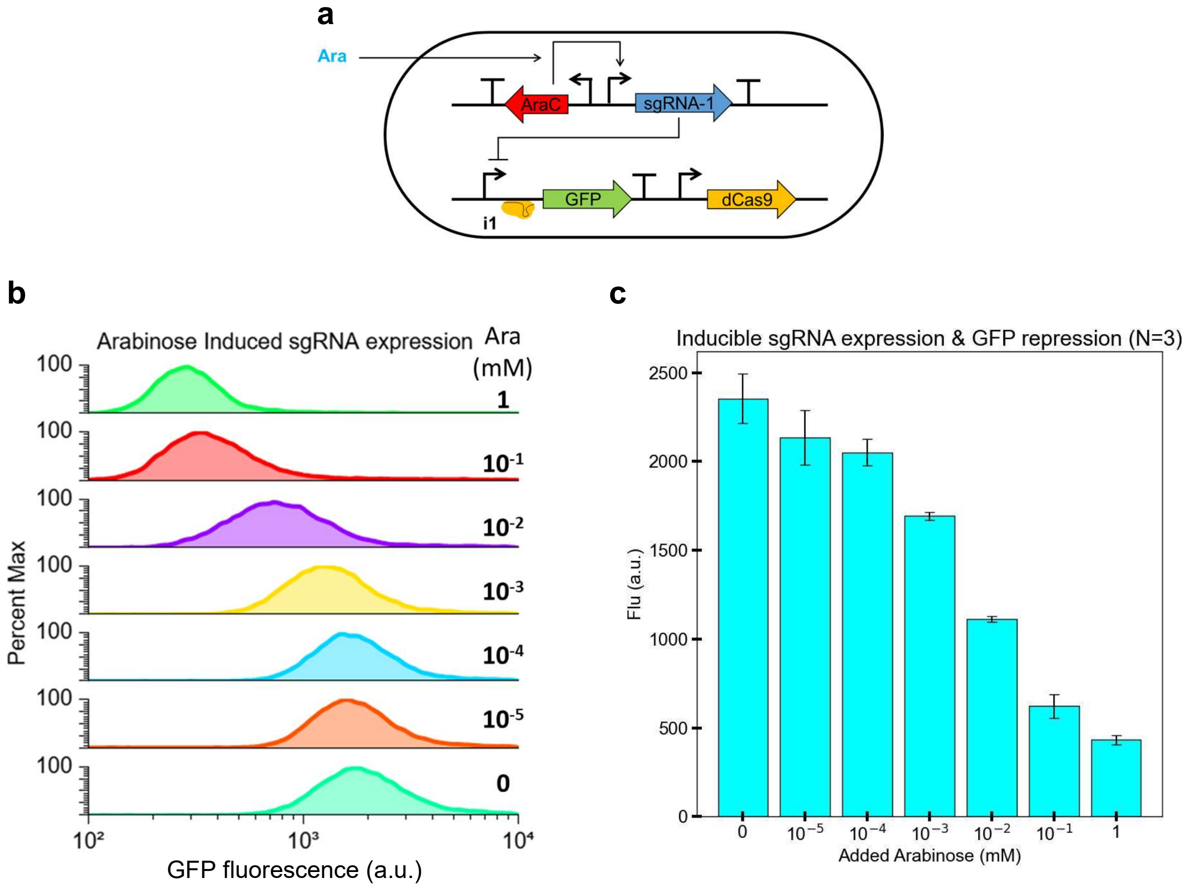


**Fig. S6.2: Arabinose-induced sgRNA expression in bacteria to repress GFP by CRISPRi.**

(a) Schematic of the TOP10_pBAD-sgRNA-1_dCas9-GFP_NOT_i1 cell, where *sgRNA-1* is transcribed from the arabinose-inducible pBAD promoter. Once expressed, *sgRNA-1* forms a complex with dCas9 protein and binds to the *GFP* promoter to repress it. (b) Bacterial cells were grown for 16 hours in LB media supplemented with varying concentrations of arabinose. Samples were run through a flow cytometer to record the fluorescence of the bacterial population. These representative plots are from a single experiment (N=1). (c) Bar plots show the population's mean fluorescence value at the end of the 16 hour induction (N=3).

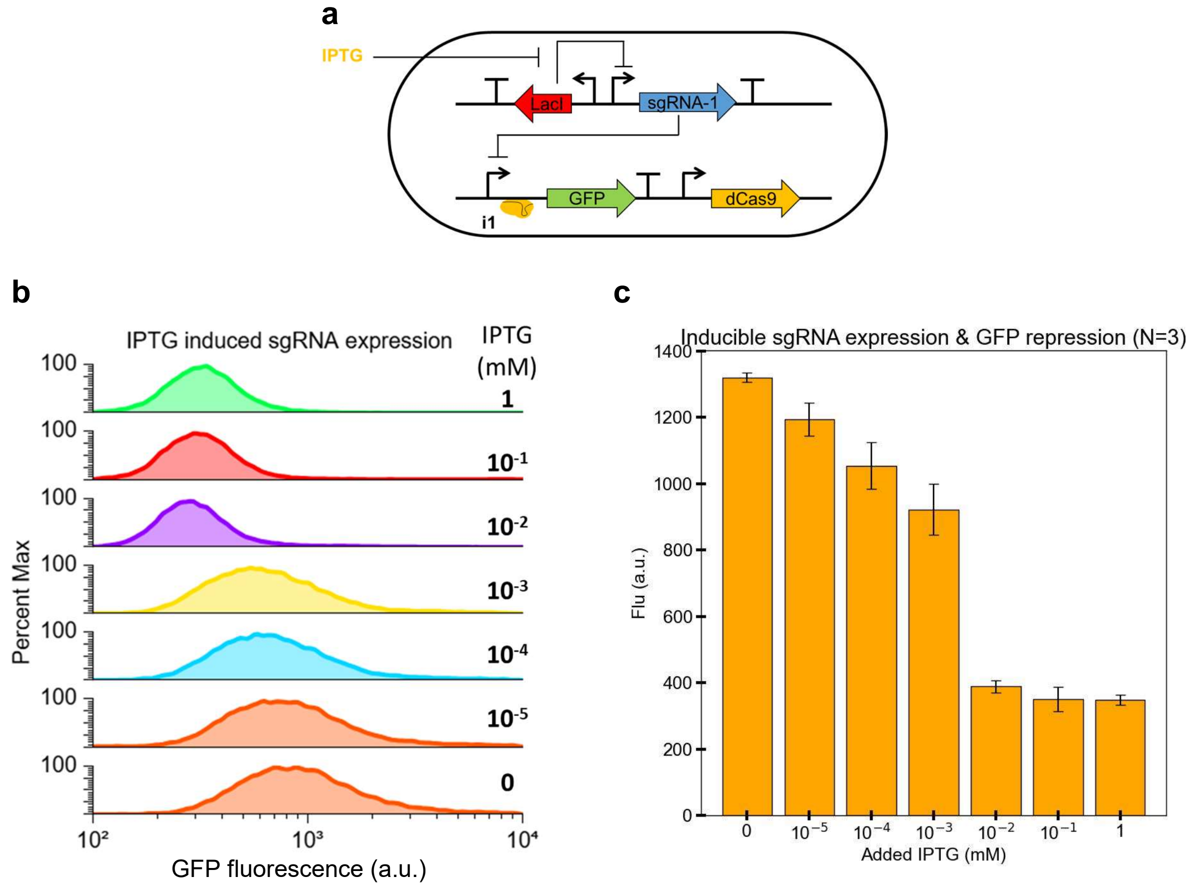


**Fig. S6.3: IPTG-induced sgRNA expression in bacteria to repress GFP by CRISPRi.**

(a) Schematic of the TOP10_pLac-sgRNA-1_dCas9-GFP_NOT_i1 cell, where *sgRNA-1* is transcribed from the IPTG-inducible pLac promoter. Once expressed, *sgRNA-1* forms a complex with dCas9 protein and binds to the *GFP* promoter to repress it. (b) Bacterial cells were grown for 16 hours in LB media supplemented with varying concentrations of IPTG. Samples were run through a flow cytometer to record the fluorescence of the bacterial population. These representative plots are from a single experiment (N=1). (c) Bar plots show the population's mean fluorescence value at the end of the 16 hour induction (N=3).

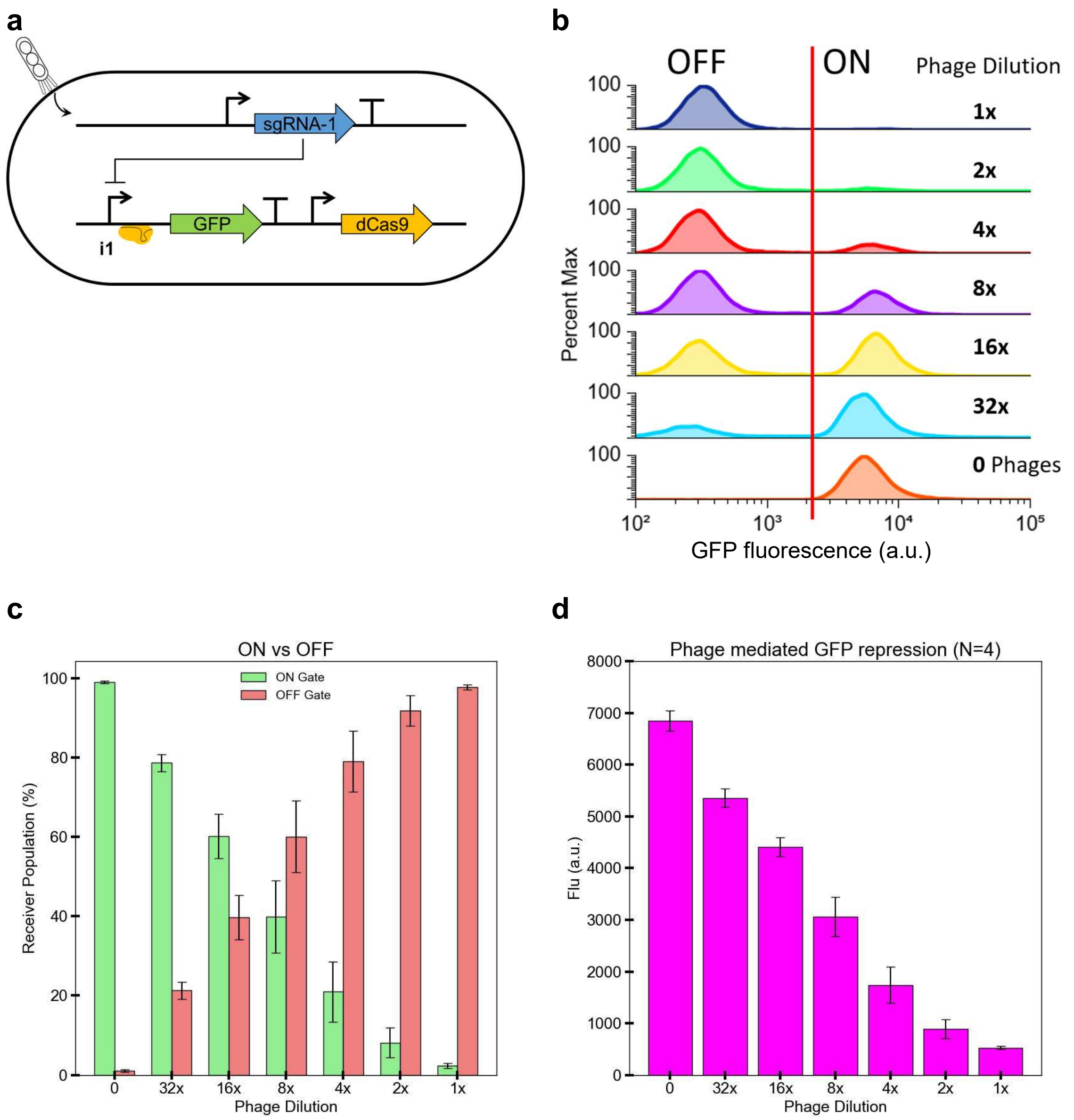


**Fig. S6.4: Phage-delivered *sgRNA-1* repressed GFP by CRISPRi.**

(a) Schematic of the TOP10F_dCas9-GFP_NOT_i1 receiver cell after infection by the sgRNA-1 phage. Upon infection, the constitutively expressed *sgRNA-1* forms a complex with dCas9 protein and binds to the *GFP* promoter to repress it. (b) Receiver cells were grown for 16 hours in LB media with varying concentrations of isolated sgRNA-1 phages (undiluted 1x concentration = $3.6 \times 10^{13}$ $mL^{-1}$, CFU assay), without antibiotic selection. Samples were run through a flow cytometer to record the fluorescence of the bacterial population. Number of GFP ON and OFF cells was determined using a fluorescence threshold value (vertical red line). These representative plots are from a single experiment (N=1). (c) Plot of the percentage receiver populations at the end of the 16 hour incubation for each phage dilution: infected (OFF) vs uninfected (ON) (N=3). (d) Bar plots show the population's mean fluorescence value at the end of the 16 hour incubation for each phage dilution (N=3).

**a**

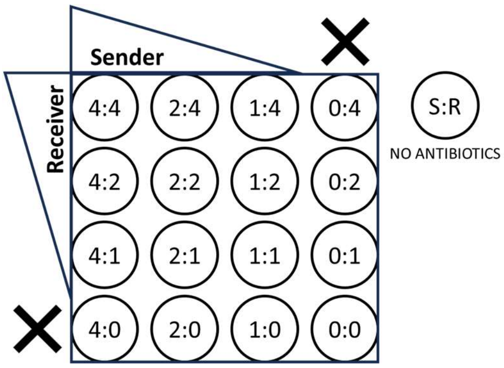


**b**

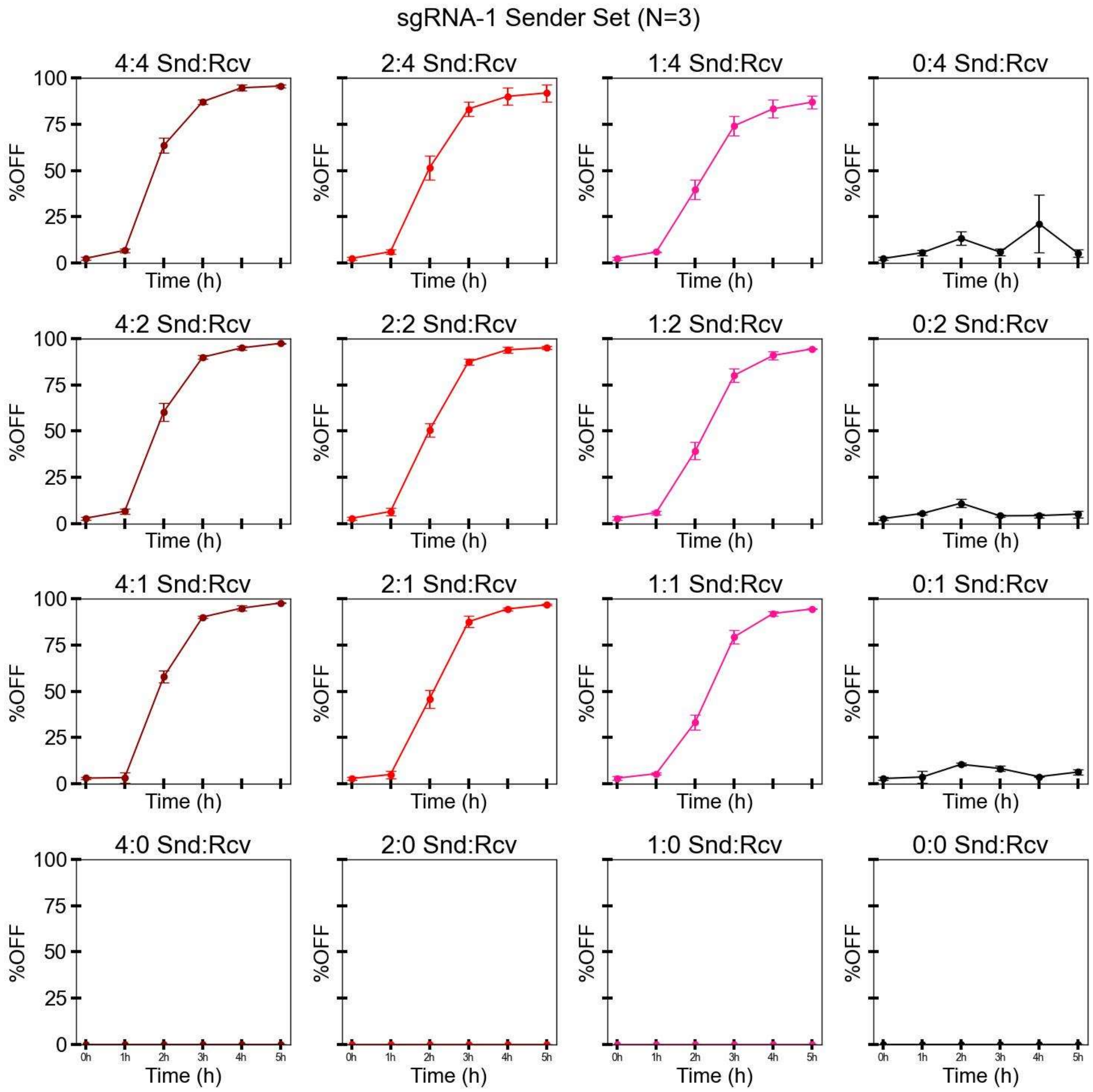


**Fig. S6.5: Infection over time of receiver cells grown with sgRNA-1 sender cells.**

Receiver and sender cells were mixed in different density ratios (a), before incubating them at 37°C 205 cpm for 5 hours. Each hour a fraction of the culture was used for flow cytometry to obtain %infected cells (b). sgRNA-1 plasmid is based on the pSEVA191 vector (*AmpR*, pBBR322).

a

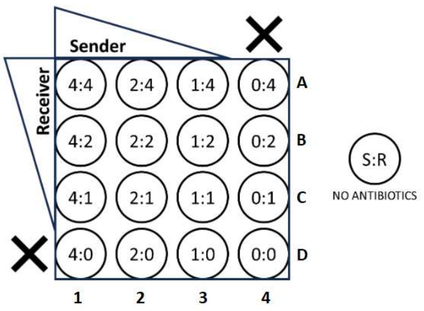


b

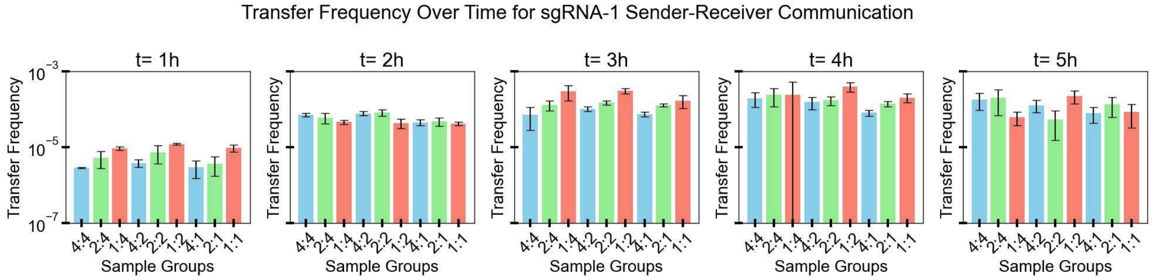


**Fig. S6.6: Transfer frequency calculated for sgRNA-1 sender-to-receiver cells without antibiotics.**

Receiver and sender cells were mixed in different density ratios according to the plate map shown in (a), and incubated at 37 °C, 200 rpm, for 5 hours. (b) Each hour a fraction of the culture was used for flow cytometry to obtain the number of senders (S), infected receivers (Ri, GFP OFF), and uninfected receivers (Ru, GFP ON). Transfer frequency at each time-point was calculated as Ri/S*(Ri+Ru), according to the formula used to obtain transconjugant frequency in conjugation experiments (10, 11). sgRNA-1 plasmid is based on the pSEVA191 vector (*AmpR*, pBBR322). Data from Fig. S6.5. (N=3).

**a**

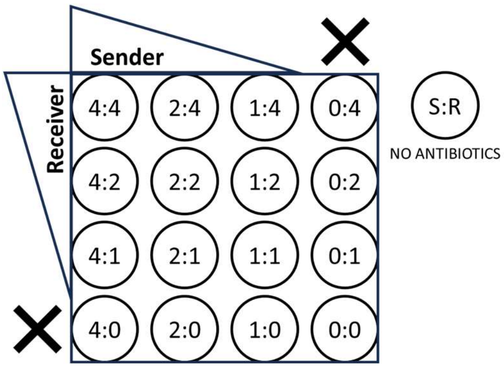


**b**

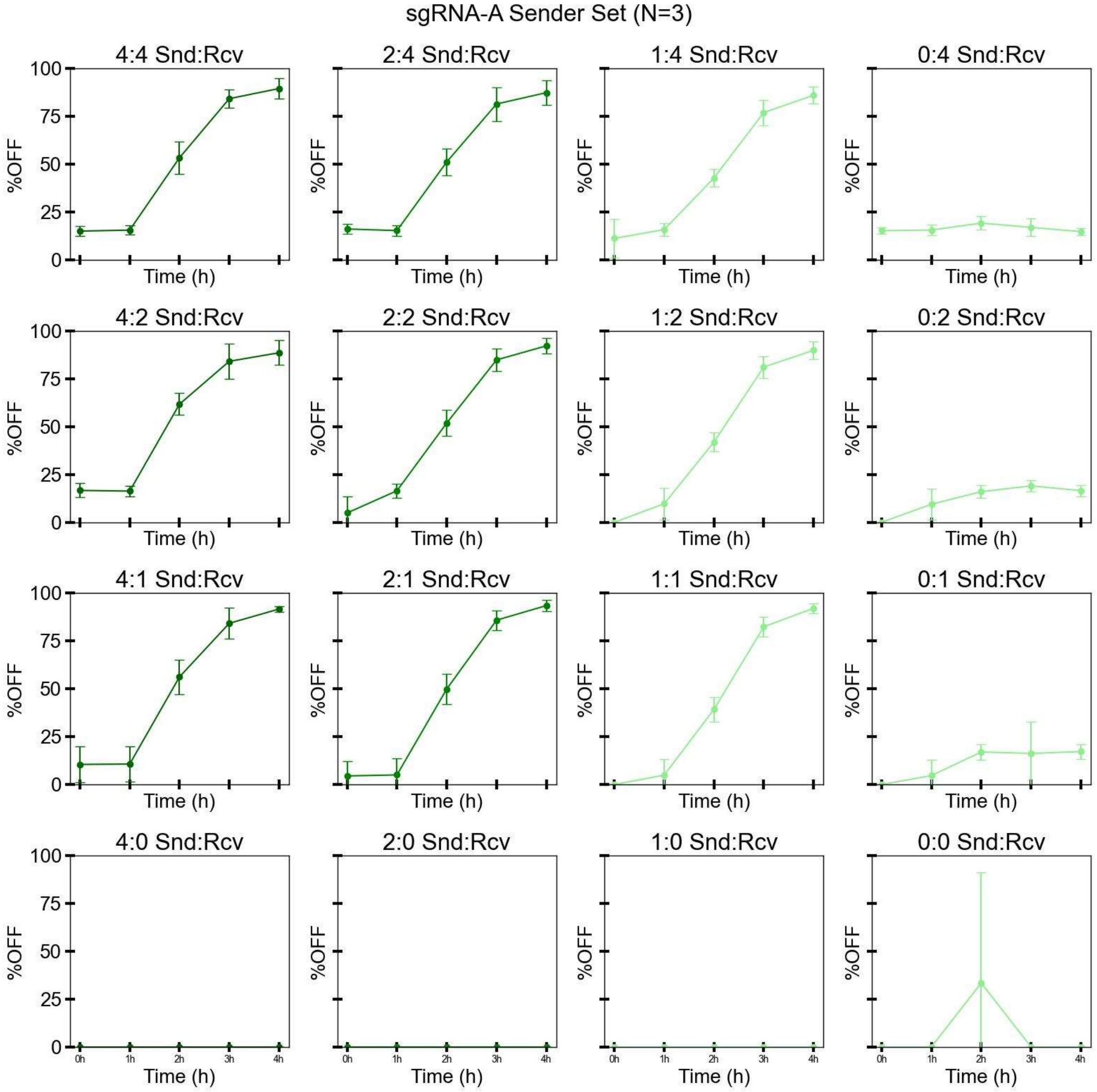


**Fig. S6.7: Infection over time of receiver cells grown with sgRNA-A sender cells.**

Receiver and sender cells were mixed in different density ratios (a) before incubating them at 37°C 205 cpm for 4 hours. Each hour a fraction of the culture was used for flow cytometry to obtain %infected (=%OFF) cells (b). sgRNA-A plasmid is based on the pSEVA261 vector (*KanR*, p15A).

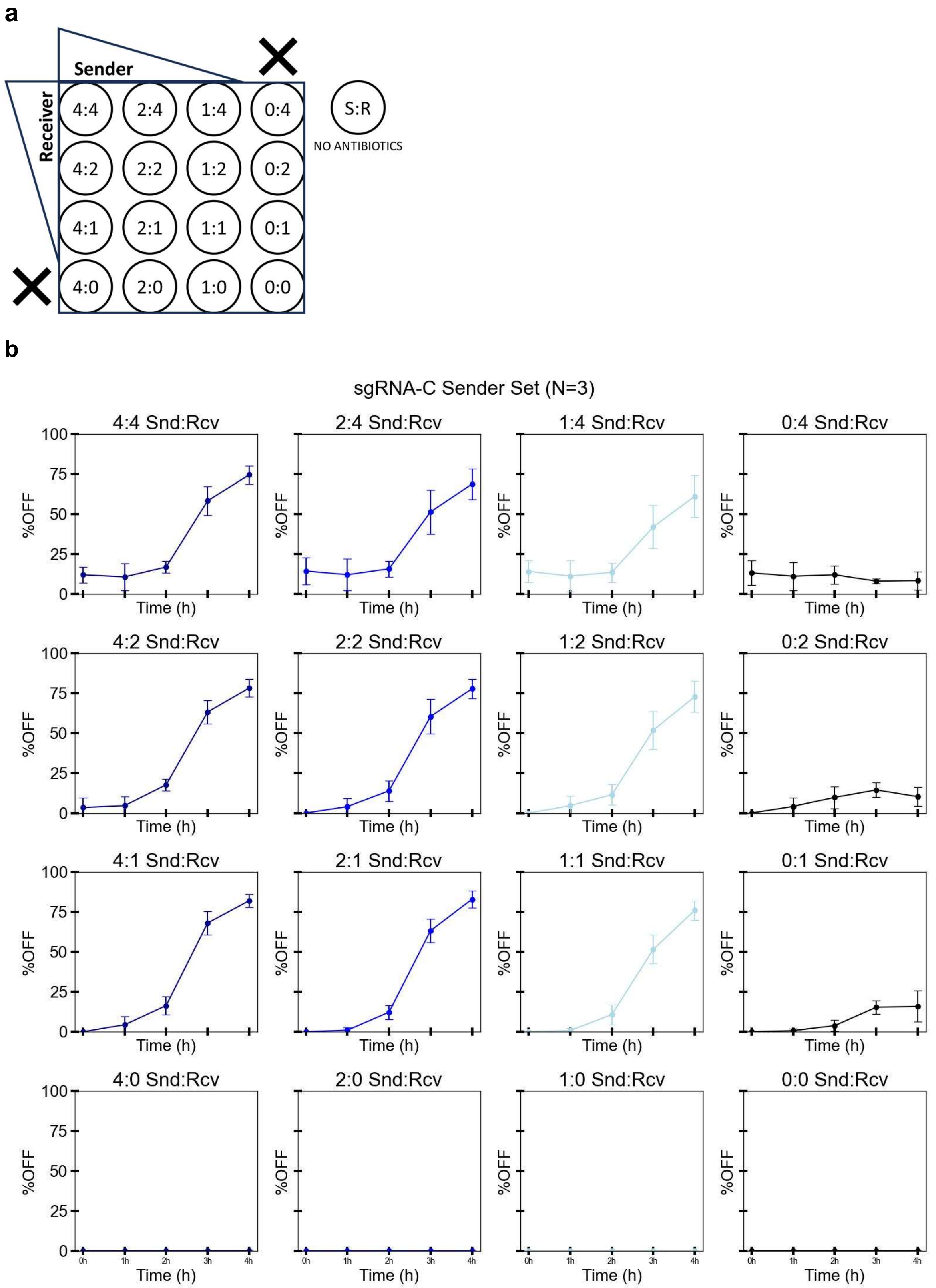
a
Sender
Receiver
4:4
2:4
1:4
0:4
S:R
NO ANTIBIOTICS
4:2
2:2
1:2
0:2
4:1
2:1
1:1
0:1
4:0
2:0
1:0
0:0
b
sgRNA-C Sender Set (N=3)
4:4 Snd:Rcv
2:4 Snd:Rcv
1:4 Snd:Rcv
0:4 Snd:Rcv
4:2 Snd:Rcv
2:2 Snd:Rcv
1:2 Snd:Rcv
0:2 Snd:Rcv
4:1 Snd:Rcv
2:1 Snd:Rcv
1:1 Snd:Rcv
0:1 Snd:Rcv
4:0 Snd:Rcv
2:0 Snd:Rcv
1:0 Snd:Rcv
0:0 Snd:Rcv
%OFF
Time (h)
0
25
50
75
100
0h
1h
2h
3h
4h

**Fig. S6.8: Infection over time of receiver cells grown with sgRNA-C sender cells.**

Receiver and sender cells were mixed in different density ratios (a) before incubating them at 37°C 200 cpm for 4 hours. Each hour a fraction of the culture was used for flow cytometry to obtain %infected (=%OFF) cells (b). sgRNA-C plasmid is based on the pSEVA191 vector (*AmpR*, pBBR322).

**a**

| Sender sgRNA | Replication origin (vector) | Selection marker |
|---|---|---|
| sgRNA-1 | pBBR322 (pSEVA191) | AmpR |
| sgRNA-A | p15A (pSEVA261) | KanR |
| sgRNA-C | pBBR322 (pSEVA191) | AmpR |

**b**

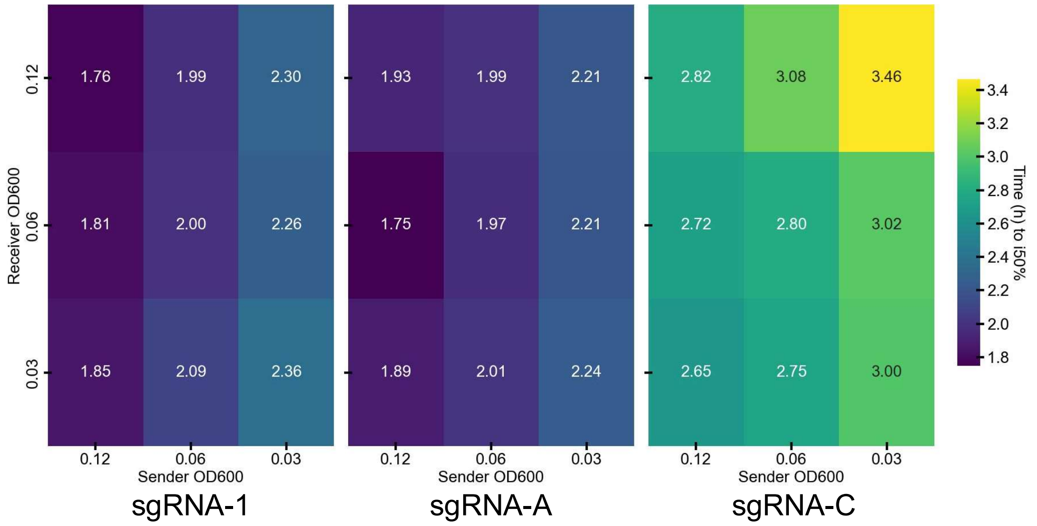


**Fig. S6.9: Time to repression of 50% receivers when co-cultured with 3 different i-CRISPRi sender variants.**

(a) Table outlines key properties of the 3 sender variants used for repression of the GFP gene in the cognate receiver variants. The sender is named after the sgRNA its phagemid encodes for.

(b) Data from figures S6.5, S6.7, and S6.8 were used to calculate the time (in hours) it takes for 50% receivers cells to see repression of their GFP gene, when co-cultured with different sender cells (sgRNA-1, sgRNA-A, sgRNA-C) without antibiotic selection.

#### Note S7: NOT- & YES- gate implementation in bacterial cocultures

This Supplementary Note presents additional data related to the experiments shown in Figure 5.

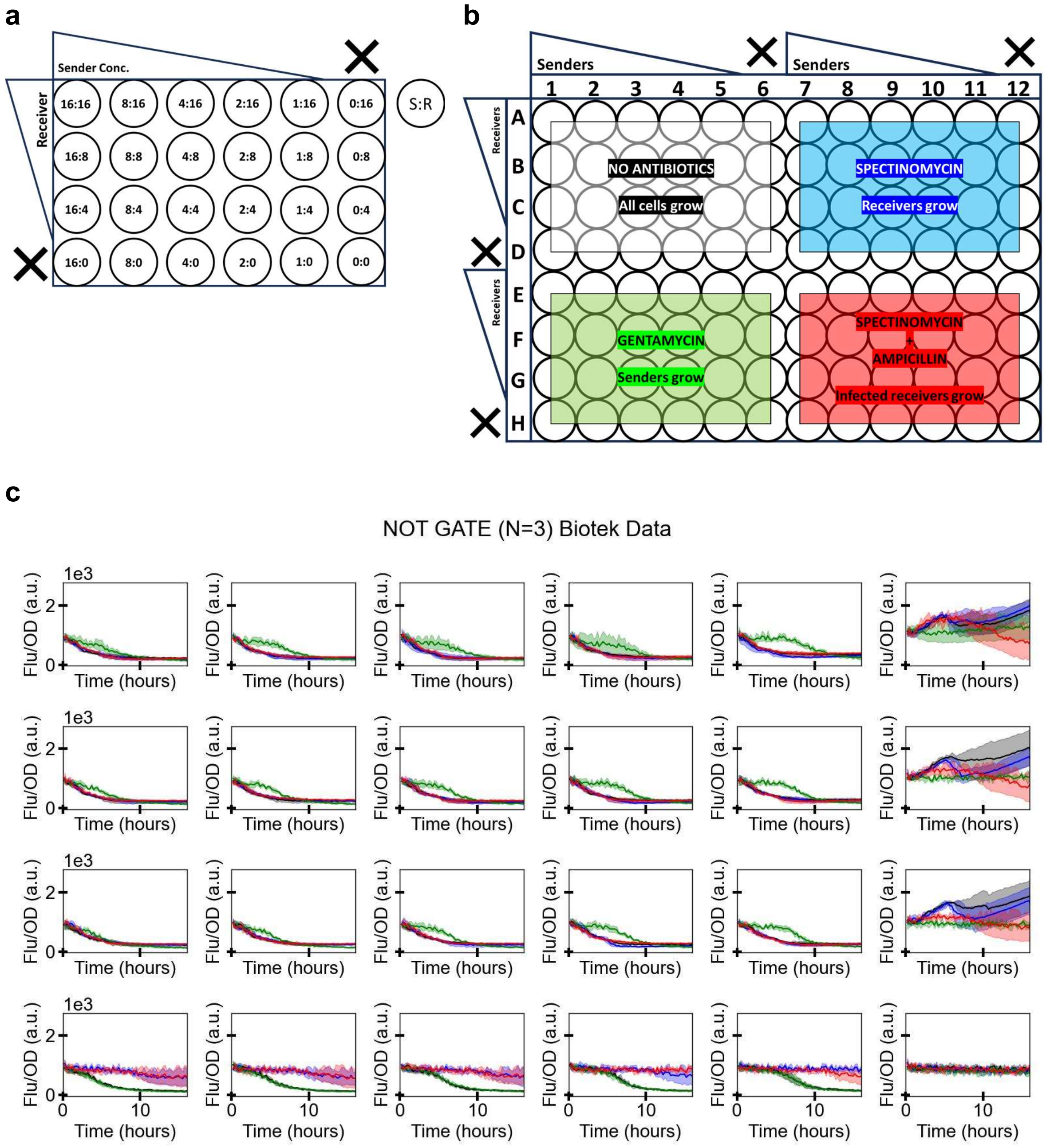


**Fig. S7.1: NOT Gate: fluorescence over time (post-selection).**

(a) Plate map of sender-receiver ratios used. (b) Antibiotic selection colour code. (c) Fluorescence over time (plate-reader data; fluorescence normalised by $OD_{600}$) for receivers (Spec; blue), senders (Gent; green), infected receivers (Amp+Spec; red), and all cells (N/A ab; black). Senders and receivers of different initial densities were grown (N=3).

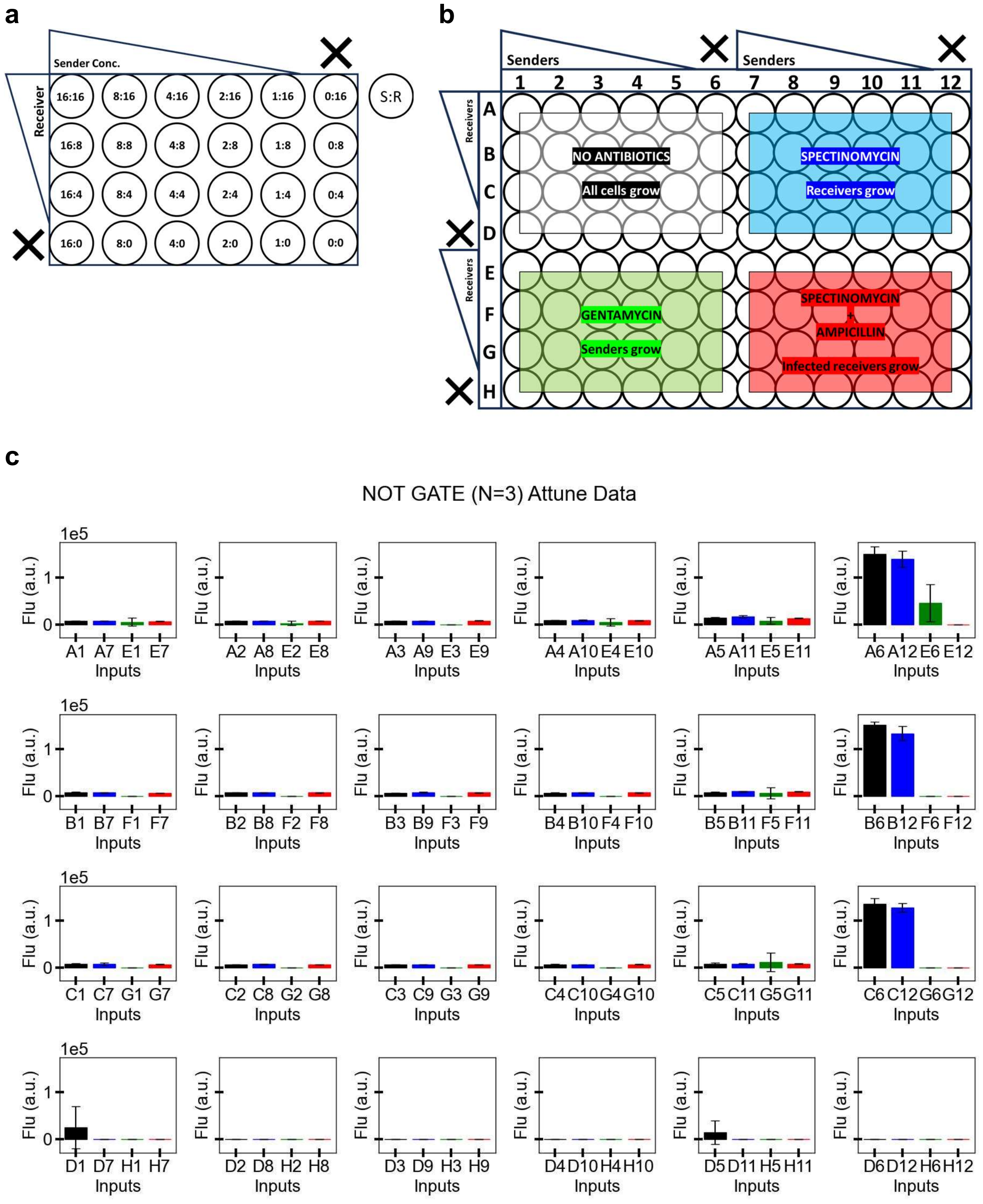


**Fig. S7.2: NOT Gate: mean fluorescence of receiver cells at 16th hour (post-selection).**

(a) Plate map of sender-receiver ratios used. (b) Antibiotic selection colour code (c) NOT GATE: Mean fluorescence at t=16h post selection (flow cytometer data) for receivers (Spec; blue), senders (Gent; green), infected receivers (Amp+Spec; red), and all cells (N/A ab; black). Senders and receivers of different initial densities were grown (N=3).

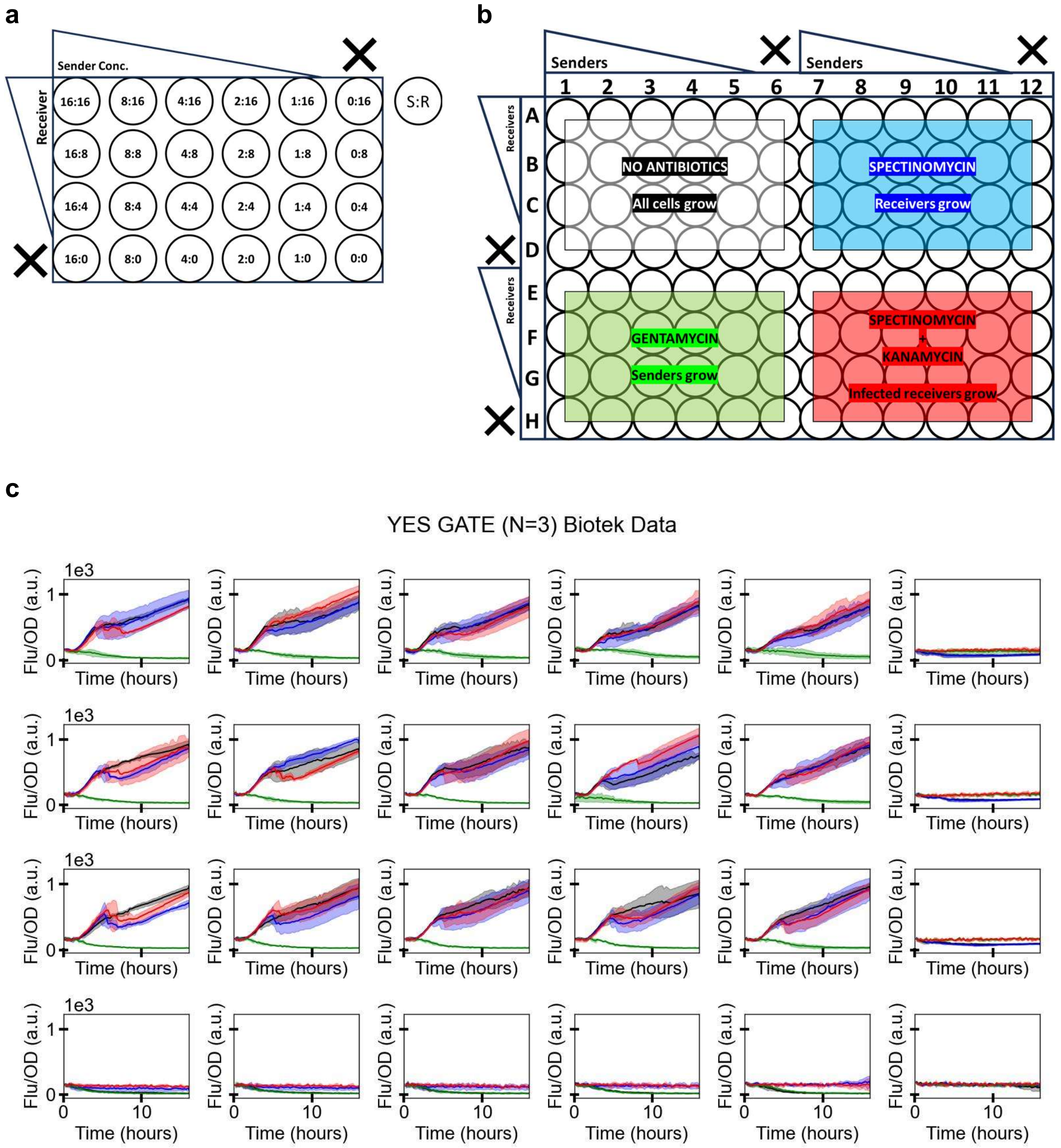


**Fig. S7.3: YES Gate: fluorescence over time (post-selection).**

(a) Starting ratios of sender:receiver used. (b) 96-well plate map and antibiotic selection colour code. (c) YES GATE: Fluorescence over time (plate-reader data; fluorescence normalised by OD) for receivers (Spec; blue), senders (Gent; green), infected receivers (Amp+Spec; red), and all cells (N/A ab; black). Senders and receivers of different initial densities were grown (N=3).

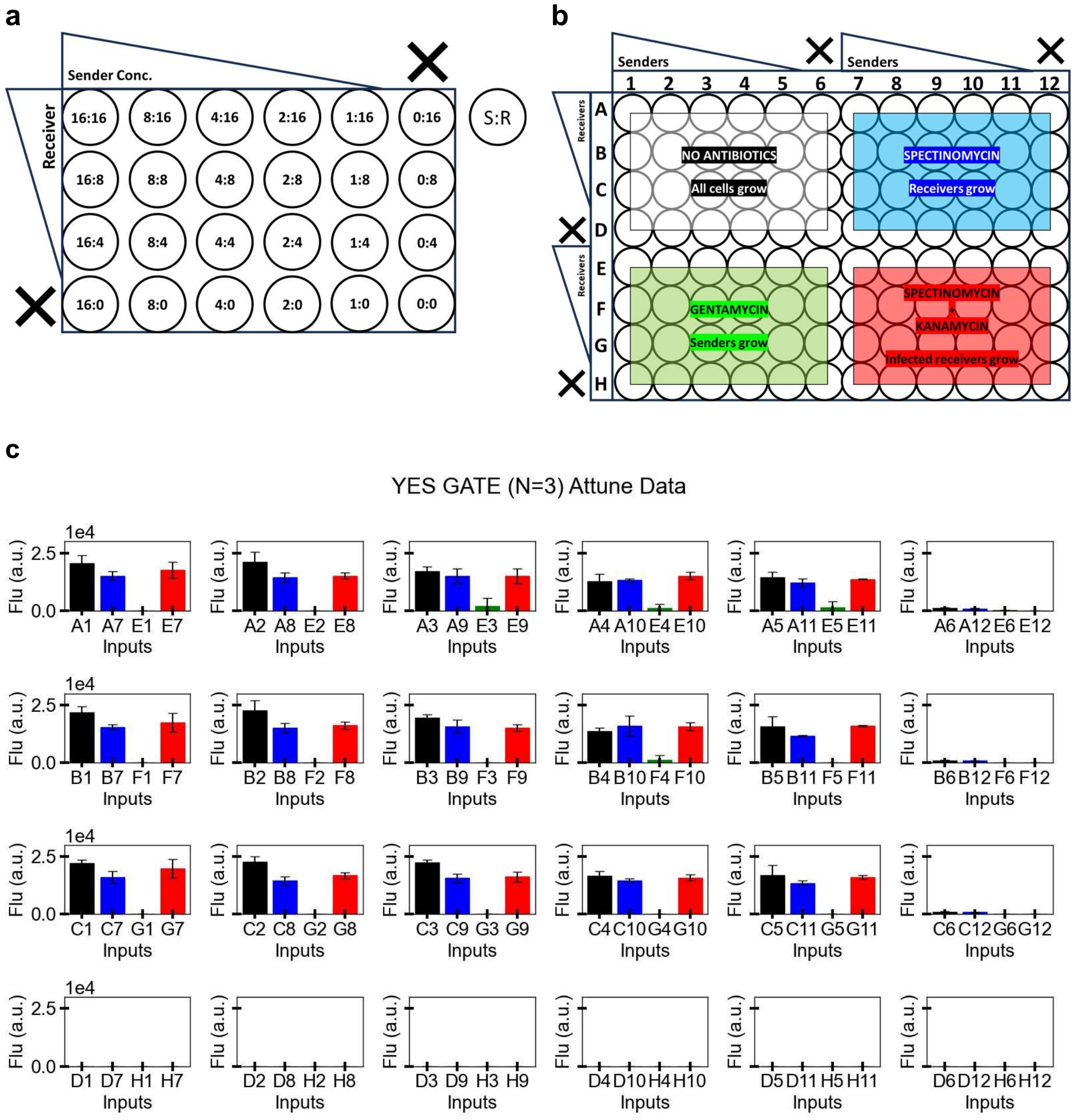


**Fig. S7.4: YES Gate: mean fluorescence of receiver cells at 16th hour (post-selection).**

(a) Plate map of sender-receiver ratios used. (b) Antibiotic selection colour code. (c) Mean fluorescence at t=16h post selection (flow cytometer data) for receivers (Spec; blue), senders (Gent; green), infected receivers (Amp+Spec; red), and all cells (N/A ab; black). Senders and receivers of different initial densities were grown (N=3).

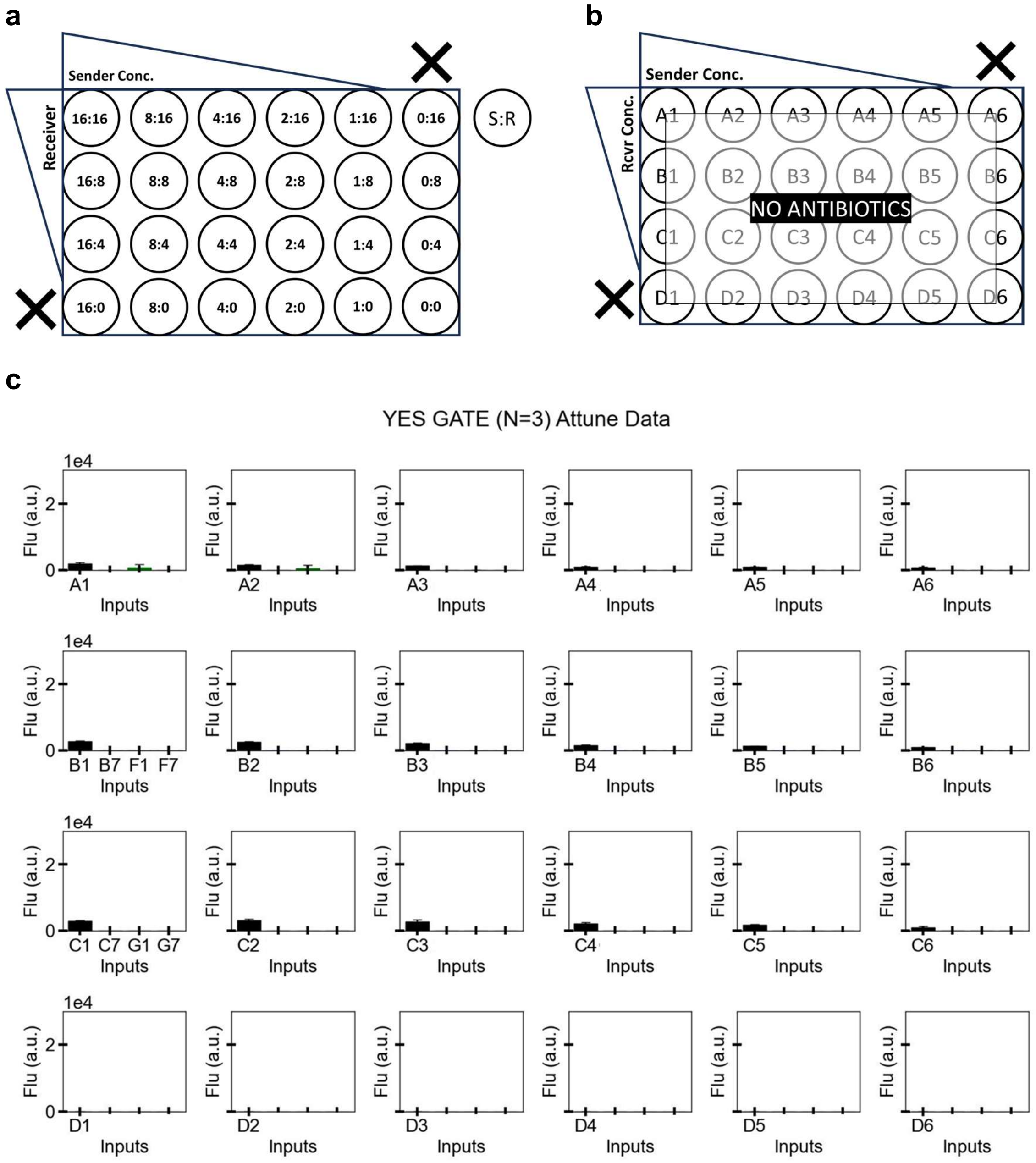


**Fig. S7.5: YES Gate: mean fluorescence of receiver cells at 4th-hour (pre-selection).**

(a) Plate map of sender-receiver ratios used. (b) Well labels on the plate for different sender-receiver ratios used. (c) Mean fluorescence at t=4h of sender-receiver coculture before antibiotic selection is applied (flow cytometer data) (N=3).

#### Note S8: AND- & AND-AND-NOT- gate implementation in bacterial cocultures

This Supplementary Note presents additional data related to the experiments shown in Figure 6.

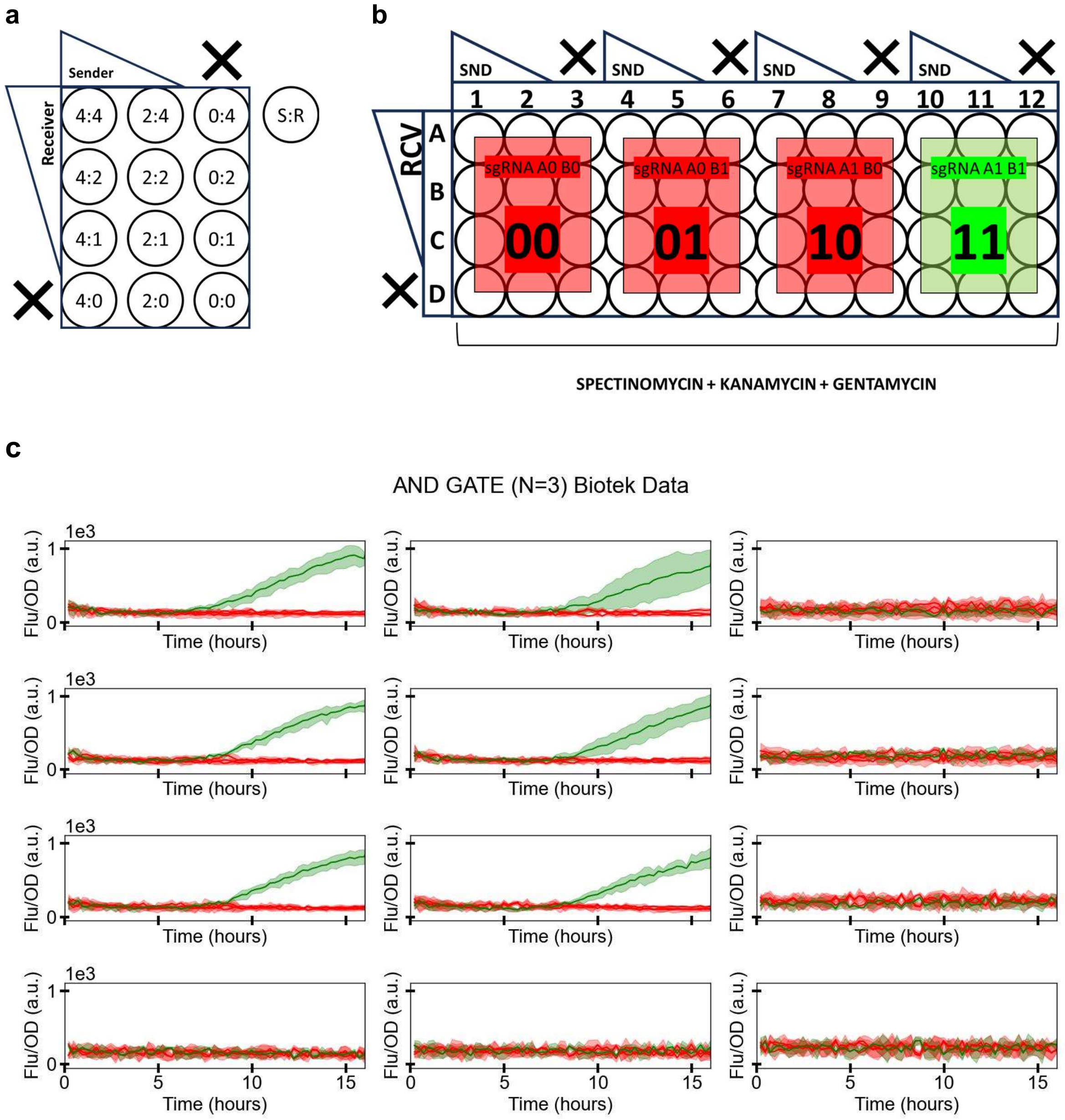


**Fig. S8.1: AND Gate: fluorescence over time (post-selection).**

(a) Plate map of sender-receiver ratios used. (b) Well labels on the plate for different sender-receiver ratios and input senders used. (c) AND gate: Fluorescence over time (Biotek data; normalised by $OD_{600}$) for receiver cells infected with phage inputs 00, 01, 10 (red), and 11 (green), in sender-receiver cocultures of different starting cell densities (N=3).

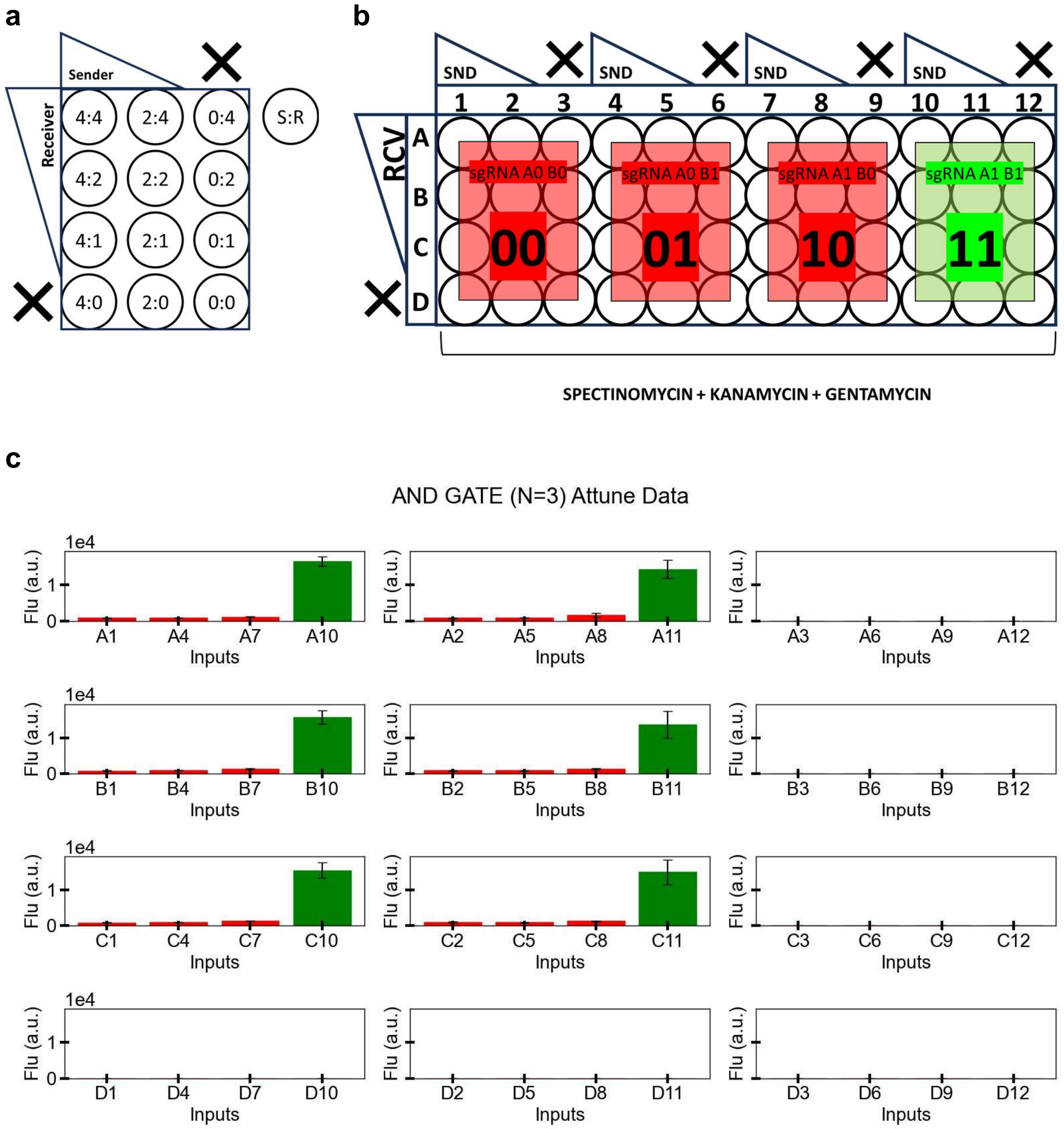


**Fig. S8.2: AND Gate: mean fluorescence of receiver cells at 16th hour (post-selection).**

(a) Plate map of sender-receiver ratios used. (b) Well labels on the plate for different sender-receiver ratios and input senders used. (c) Mean fluorescence at t=16h post selection (flow cytometer data) for receiver cells infected with phage inputs 00, 01, 10 (red), and 11 (green), in sender-receiver cocultures of different starting cell densities (N=3).

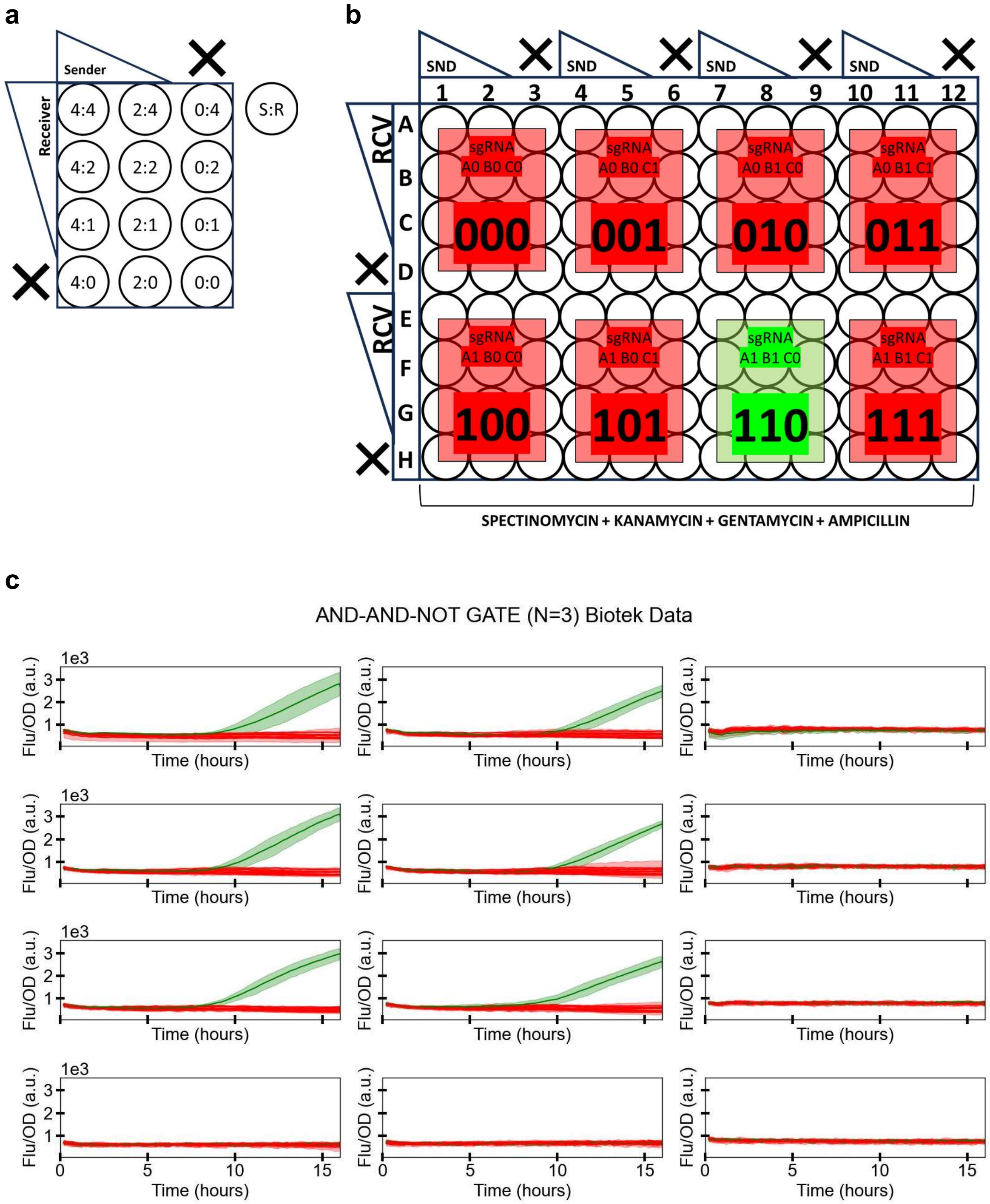


**Fig. S8.3: AND-AND-NOT Gate: fluorescence over time (post-selection).**

(a) Plate map of sender-receiver ratios used. (b) Well labels on the plate for different sender-receiver ratios and input senders used. (c) AND-AND-NOT gate: Fluorescence over time (Biotek data; normalised by $OD_{600}$) for receiver cells infected with phage inputs 000, 001, 010, 011, 100, 101, 111 (red), and 110 (green), in sender-receiver cocultures of different starting cell densities (N=3).

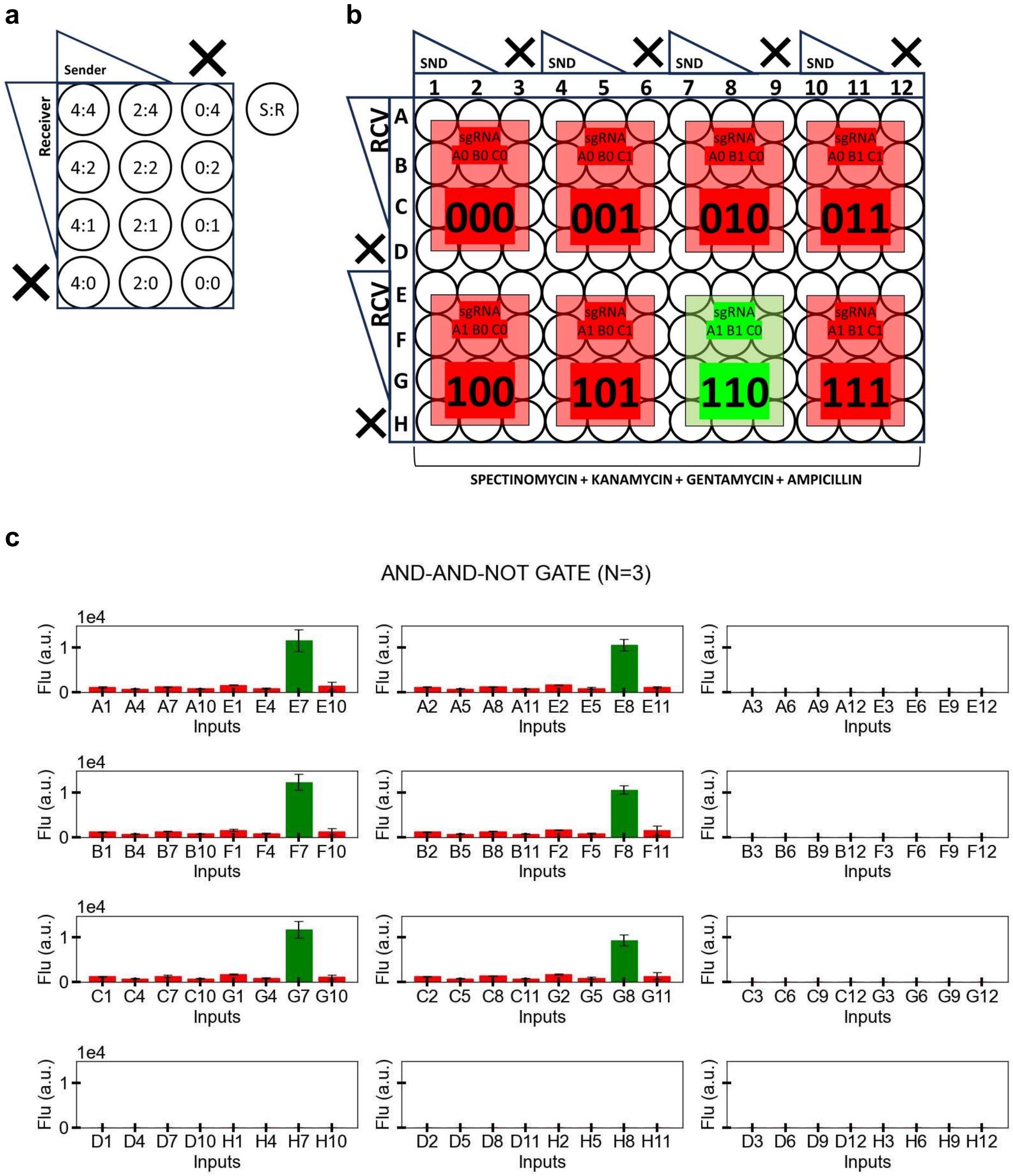


**Fig. S8.4: NOT Gate: mean fluorescence of receiver cells at 16th hour (post-selection).**

(a) Plate map of sender-receiver ratios used. (b) Well labels on the plate for different sender-receiver ratios and input senders used. (c) AND-AND-NOT gate: Mean fluorescence at t=16h post selection (flow cytometer data) for receiver cells infected with phage inputs 000, 001, 010, 011, 100, 101, 111 (red), and 110 (green), in sender-receiver cocultures of different starting cell densities. Senders and receivers of different initial densities were grown (N=3).

**Note S9:** 20% activation time of ‘ON-state’ receiver cells.

In this Supplementary Note, we calculate the time of switching of the different logic gates using plate reader data. We chose the 20% activation threshold because by this fold-change the fluorescence (Flu/$OD_{600}$) values reach above the basal noise levels for all gates.

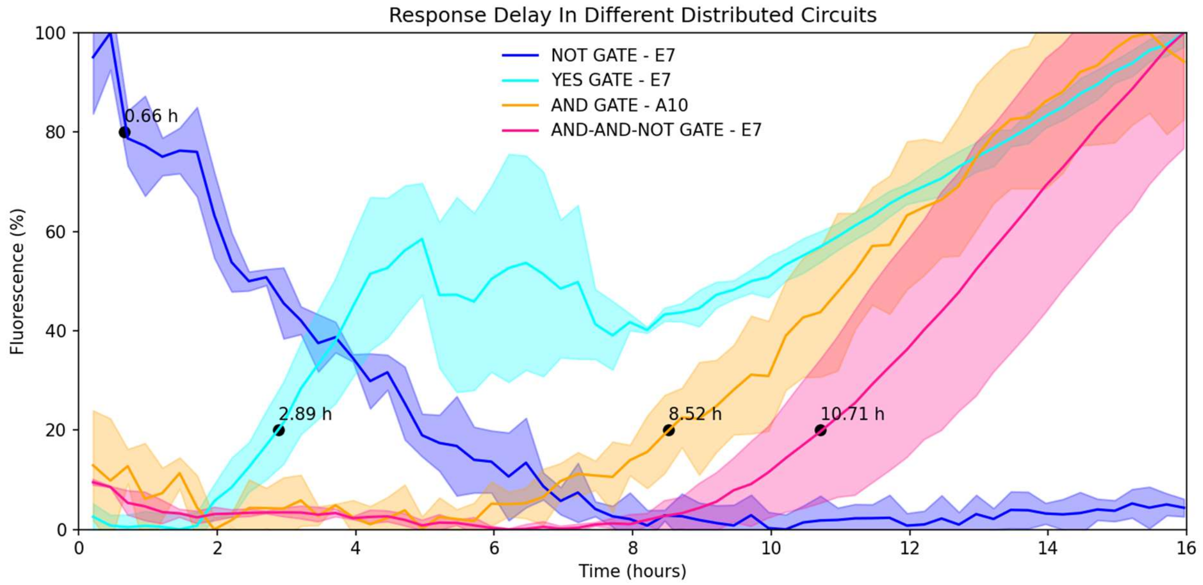


**Fig. S9.1: 20% activation time for single- & multi- input receivers.**

Fluorescence (normalised) of ‘ON-state’ infected receiver cells over time. Black dots represent 20% change from their initial fluorescence values. Solid lines are mean values from 3 repeats, and the shaded region represents the standard deviation.

The figure above illustrates the GFP-based response times for 'ON-state' infected receivers of the constructed NOT (dark blue), YES (cyan), AND (orange), and AND-AND-NOT (pink) biological gates. Here, the response time is defined as the time at which a 20% (expected) change in fluorescence is observed. The NOT gate receivers exhibit the fastest response time at 0.66 hours, followed by YES at 2.89 hours, AND at 8.52 hours, and AND-AND-NOT gate receivers at 10.71 hours.

**Note S10:** Screening AND-AND-NOT circuit design variants.

We built and characterised a total of 7 different designs of the AND-AND-NOT circuits (**Fig.** S10.1), which varied in one or more of three properties:

(1) *the sgRNA sequences used*, since sgRNA sequences with full complementarity to the promoter region can also have different CRISPRi targeting efficiencies (12),

(2) *sgRNA targeting the template (T) or the non-template (NT) strand*, since previous work showed transcription repression on both strands (albeit more efficiently on NT) when targeting the promoter region (13), and

(3) *the positions of multiple sgRNA targets at / near the same promoter*, since multiple targeting sites regulating a promoter can effectively create NOR-gate logic from multiple inputs (14).

The final circuit presented in the main text (AAN_4) was identified by a three step screening process. First, the 7 designs were co-transformed with the phagemids constitutively expressing the sgRNAs to test if the circuit showed the expected behaviour for 3 (000, 110, 111) key input combinations (**Fig.** S10.2). Only 5 of these 7 showed an ON state for the '110' input combination with significantly higher fluorescence output than the other two combinations. Both designs that showed no GFP activation have common sgRNA inputs (iA, iC, and iE) and fail to activate GFP expression, suggesting that at least one of input sgRNAs iA and iC failed to release repression due to internal sgRNAs X and Y (**Fig.** S10.1). However, both iA and iC inputs work in other designs 4–7, wherein iA is used at exactly the same position. This suggests that a likely cause of the failure of designs 2–3 is the difference in the position of iC targeting. iC is able to repress the GFP promoter directly (designs 4–7), but not the promoter of the internal sgRNA-Y (2–3).

Next, the 5 promising circuits (AAN_1, AAN_4, AAN_5, AAN_6, AAN_7) from the previous step were tested using co-culturing between sender and receiver cells for 20 h, using the 3 input combinations (**Fig.** S10.3). Circuit AAN_1 showed no activation in the '110' input combination, suggesting that at least one of sgRNAs iF and i2 failed to transmit from sender to receiver cells and / or release repression due to internal sgRNAs P and Q. In line with the co-transformation results above, circuits AAN_5 and AAN_7 showed only weak activation. Designs 5 & 7 use the same external and internal sgRNAs as the strongly activating designs 4 & 6, with the key difference being the DNA strand targeted for repressing the internal sgRNAs (**Fig.** S10.1). While NT targeting in designs 5 & 7 leads to weak repression, T targeting in designs 4 & 6 leads to strong repression, which seems contrary to previous findings (13). However, it must be noted that the position of sgRNA targeting is also different in the two designs: in 5 & 7 the region immediately downstream of the promoter is targeted, whereas in 4 & 6 the region inside the promoter (between -10 and -35, inspired from (15)) is targeted.

Finally, the best designs from the previous step (AAN_4, AAN_6) were tested again with all input combinations using co-culturing between sender and receiver cells, where they showed a conservative activation fold of 7.17 and 4.78, respectively. Therefore, circuit AAN_4 was used for further characterisation and presented in the main text.

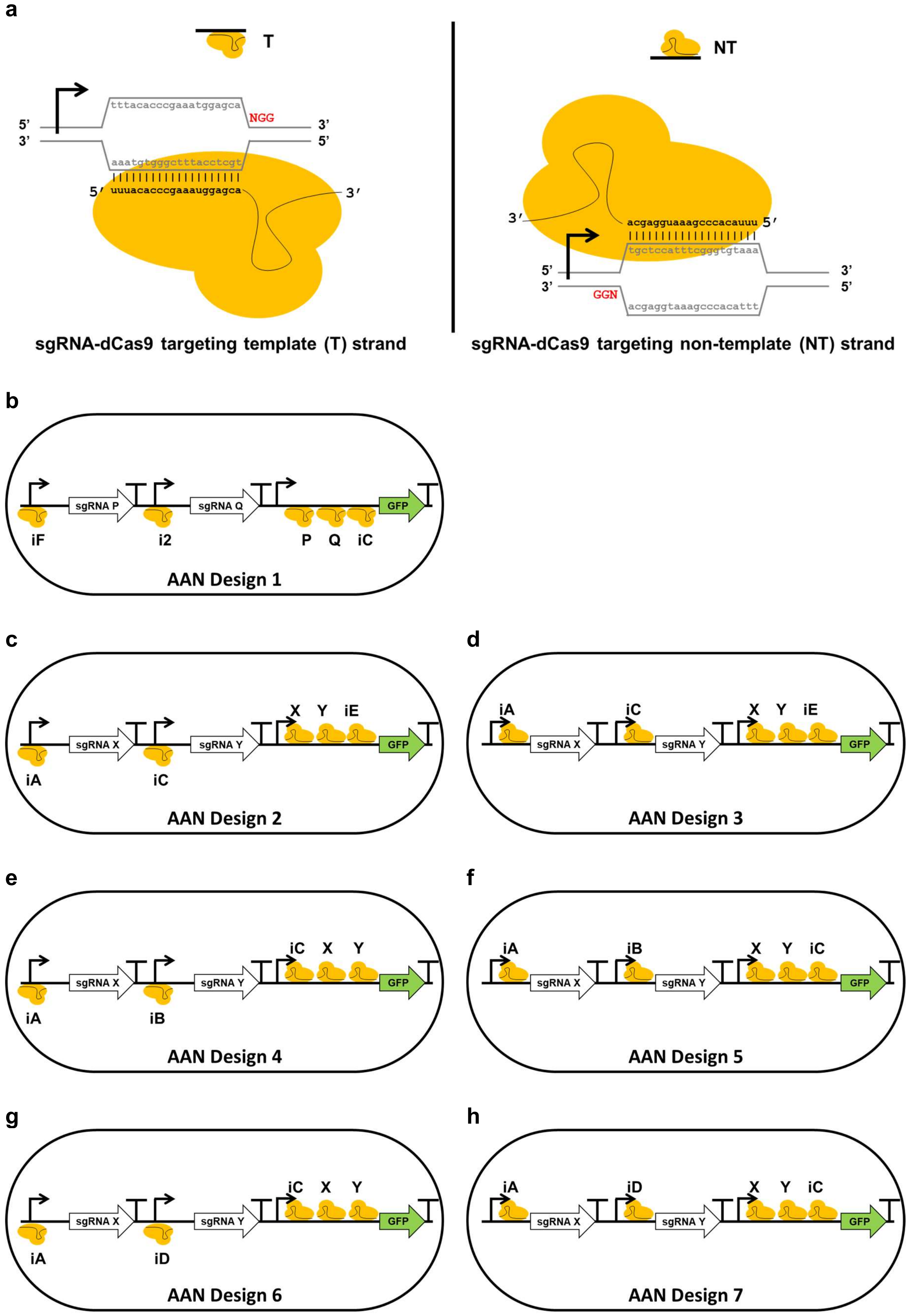


**Fig. S10.1: Strain variants designed to implement the AND-AND-NOT gate in receiver cells.**

(a) Schematics show simplified (top) and more detailed (bottom) cartoon representations of template (T, left) and not-template (NT, right) strand binding by two example sgRNA sequences in sgRNA+dCas9 complexes for

CRISPRi. The protospacer adjacent motif (PAM) sequence is highlighted in red. (b-h) Seven different design variants (AAN_1 to AAN_7) of the AND-AND-NOT circuits. Designs vary in the sgRNAs used, the orientation of the sgRNA targeting (template / non-template), and the position of targeting at / near the promoter. All sgRNA binding sites are identified by a number or a letter, together with an sgRNA+dCas9 symbol. sgRNAs expressed by externally delivered phagemids are prefixed with an ‘i’, whereas those expressed internally are not. All plasmid maps are available in the Supplementary files.

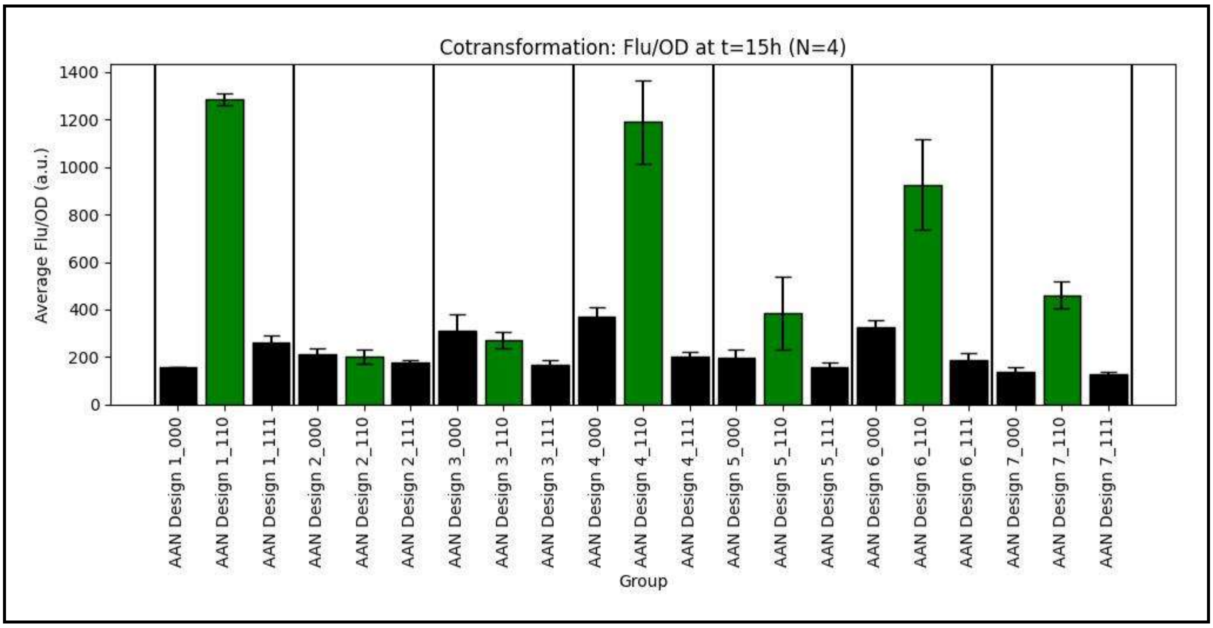


**Fig. S10.2: AND-AND-NOT gate co-transformations.**

Normalised fluorescence of Top10F cells transformed with 1 of 7 AND-AND-NOT variant plasmids (**Fig.** S10.1), and 3x sgRNA phagemids of different input combinations (000, 110, 111). Co-transformants were grown in a plate reader. Data show mean±SD (from N=4 repeats) of fluorescence/$OD_{600}$ at 15 h.

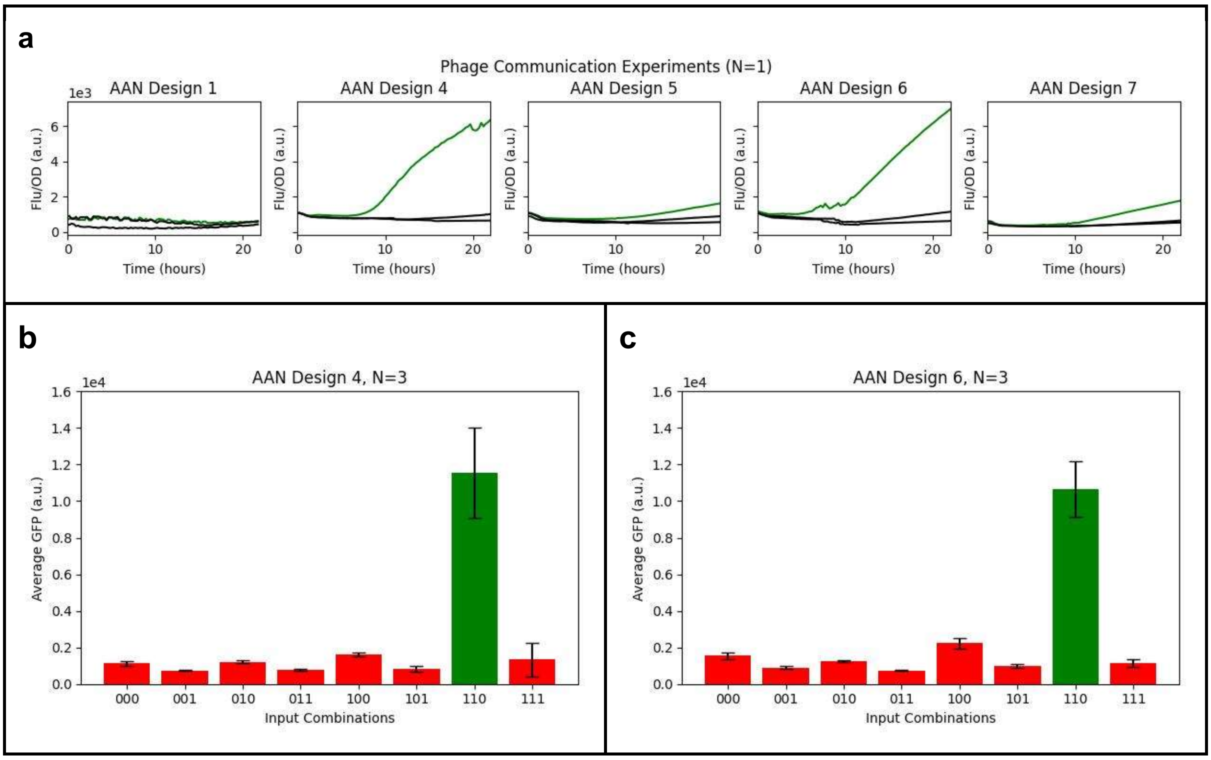


**Fig. S10.3: AND-AND-NOT gate variants: sender to receiver communication.**

(a) Fluorescence over time (Biotek plate-reader data; normalised by $OD_{600}$) for 5 AND-AND-NOT receiver variants (AAN_1, AAN_4, AAN_5, AAN_6, AAN_7) receiving input combinations 000, 110, and 111, in sender-receiver cocultures (N=1). Black lines show 000 & 111 input combinations. Green line shows the 110 input combination. (b,c) Mean fluorescence at t=16h post selection (flow cytometer data) for receiver cells (AAN_4, AAN_6) infected with all 8 combinations of inputs, in sender-receiver cocultures (N=3). Using a conservative estimate, the fold-change between the ON state and the worst OFF state was 7.17 and 4.78 for AAN_4 and AAN_6, respectively. Only data from the AAN_4 circuit is presented in the main text.

### Supplementary Tables

#### Table S1: Transformed Strains

These were made by transforming the core strains (**Table** S3) with the respective plasmids (**Table** S2).

| **Transformed Strain** | **Core Strain** | **Additional Plasmid/s** | **Antibiotic Resistance** | **Source** | **Notes** |
|---|---|---|---|---|---|
| ER2738F_HΔgIII | ER2738F | pSEVA63_HelperΔgIII | $Tet^R$ / $Gen^R$ | This work | Fig. 1, 3, S3.1b |
| ER2738F_H | ER2738F | pSEVA63_Helper | $Tet^R$ / $Gen^R$ | This work | Fig. S3.1b |
| ER2738F_+gp3 φphagemid | ER2738F | pSB1K3_M13ps_LacZα_gIII | $Tet^R$ / $Kan^R$ | This work | Fig. S3.1a |
| TOP10_H_KanΦ | TOP10 | pSEVA63_Helper + pSB1K3_M13ps | $Str^{R\#}$ / $Gen^R$ / $Kan^R$ | This work | Fig. 1, 3 |
| TOP10_HΔgIII_gIII-KanΦ | TOP10 | pSEVA63_HelperΔgIII + pSB1K3_M13ps_LacZα_gIII | $Str^{R\#}$ / $Gen^R$ / $Kan^R$ | This work | Fig. 1, 3 |
| TOP10_H_sgRNA-1_AmpΦ | TOP10 | pSEVA63_Helper + pSEVA19_sgRNA-1_M13ps | $Str^{R\#}$ / $Gen^R$ / $Amp^R$ | This work | Fig. 4, 5 |
| TOP10_pBAD-sgRNA-1_dCas9-GFP_NOT_i1 | TOP10 | pSEVA19_AraC_pBAD-sgRNA-1 + pSEVA47_dCas9-GFP_NOT_i1 | $Amp^R$ / $Spc^R$ | This work | Fig. S6.2 |

| | | | | | |
|---|---|---|---|---|---|
| TOP10_pLac-sgRNA-1_dCas9-GFP_NOT_i1 | TOP10 | pSEVA19_LacI_pLac-sgRNA-1 + pSEVA47_dCas9-GFP_NOT_i1 | $Amp^R$ / $Spc^R$ | This work | Fig. S6.3 |
| TOP10_H_sgRNA-2_KanΦ | TOP10 | pSEVA63_Helper + SEVA26_sgRNA-2_M13ps | $Str^{R\#}$ / $Gen^R$ / $Kan^R$ | This work | Fig. 5 |
| TOP10_H_sgRNA-A1_KanΦ | TOP10 | pSEVA63_Helper + pSEVA26_sgRNA-A1_M13ps | $Str^{R\#}$ / $Gen^R$ / $Kan^R$ | This work | Fig. 6 |
| TOP10_H_sgRNA-C1_AmpΦ | TOP10 | pSEVA63_Helper + pSEVA19_sgRNA-C1_M13ps | $Str^{R\#}$ / $Gen^R$ / $Amp^R$ | This work | Fig. 6 |
| TOP10_H_sgRNA-B1_GentΦ | TOP10 | pSEVA47_Helper + pSEVA63_sgRNA-B1_M13ps | $Str^{R\#}$ / $Gen^R$ / $Spc^R$ | This work | Fig. 6 |
| TOP10F_dCas9-GFP_NOT_i1 | TOP10F | pSEVA47_dCas9-GFP_NOT_i1 | $Str^{R\#}$ / $Tet^R$ / $Spc^R$ | This work | Fig. 4, 5 |
| TOP10F_dCas9-GFP_NOT_iA | TOP10F | pSEVA47_dCas9-GFP_NOT_iA | $Str^{R\#}$ / $Tet^R$ / $Spc^R$ | This work | Fig. S6.7 |
| TOP10F_dCas9-GFP_NOT_iC | TOP10F | pSEVA47_dCas9-GFP_NOT_iC | $Str^{R\#}$ / $Tet^R$ / $Spc^R$ | This work | Fig. S6.8 |
| TOP10F_dCas9-GFP_YES_i2 | TOP10F | pSEVA47_dCas9-GFP_YES_i2 | $Str^{R\#}$ / $Tet^R$ / $Spc^R$ | This work | Fig. 5 |

| | | | | | |
|---|---|---|---|---|---|
| TOP10F_dCas9-GFP_AAN | TOP10F | pSEVA47_dCas9-GFP_AAN_4 | $Str^{R\#}$ / $Tet^{R}$ / $Spc^{R}$ | This work | Fig. 6 |
| TOP10_H_sgRNA-A0_KanΦ | TOP10 | pSEVA63_Helper + pSEVA26_sgRNA-A0_M13ps | $Str^{R\#}$ / $Gen^{R}$ / $Kan^{R}$ | This work | Fig. 5 & 6 |
| TOP10_H_sgRNA-C0_AmpΦ | TOP10 | pSEVA63_Helper + pSEVA19_sgRNA-C0_M13ps | $Str^{R\#}$ / $Gen^{R}$ / $Amp^{R}$ | This work | Fig. 6 |
| TOP10_H_sgRNA-B0_GentΦ | TOP10 | pSEVA47_Helper + pSEVA63_sgRNA-B0_M13ps | $Str^{R\#}$ / $Gen^{R}$ / $Spc^{R}$ | This work | Fig. 5 & 6 |

[#]Although these strains are resistant to streptomycin, it was not used to culture them.

#### Table S2: Plasmids

| Plasmid | Antibiotic Selection | Reference | Source |
|---|---|---|---|
| pSEVA63_Helper | $Gen^{R}$ | PAJ297<br><br>Addgene # 190432 | Gift from the Jaramillo lab. The M13 Helper machinery (without the packaging signal) was sub-cloned from M13KE into the pSEVA631 vector. |
| pSEVA47_Helper | $Spc^{R}$ | PAJ296<br>Addgene # 231420 | Gift from the Jaramillo lab. The M13 Helper machinery (without the packaging signal) was sub-cloned from M13KE into the pSEVA471 vector. |
| pSEVA63_HelperΔgIII | $Gen^{R}$ | Addgene # 190433 | This work |
| pSB1K3_M13ps | $Kan^{R}$ | Addgene # 190434 | This work |

| | | | |
|---|---|---|---|
| pSB1K3_M13ps_LacZα_gIII | $Kan^R$ | Addgene # 190435 | This work |
| pSEVA19_sgRNA-1_M13ps | $Amp^R$ | Addgene # 231409 | This work |
| pSEVA19_AraC_pBAD-sgRNA-1 | $Amp^R$ | Addgene # 231407 | This work |
| pSEVA19_LacI_pLac-sgRNA-1 | $Amp^R$ | Addgene # 231408 | This work |
| SEVA26_sgRNA-2_M13ps | $Kan^R$ | PAJ171<br>Addgene # 231412 | Gift from the Jaramillo lab. |
| pSEVA26_sgRNA-A1_M13ps | $Kan^R$ | PAJ164<br>Addgene # 231414 | Gift from the Jaramillo lab. |
| pSEVA19_sgRNA-C1_M13ps | $Amp^R$ | PAJ272<br>Addgene # 231411 | Gift from the Jaramillo lab. Expresses sgRNA A5T from Nielsen *et al.*, 2014(15). |
| pSEVA63_sgRNA-B1_M13ps | $Gent^R$ | PAJ615<br>Addgene # 231422 | Gift from the Jaramillo lab. |
| pSEVA47_dCas9-GFP_NOT_i1 | $Spc^R$ | Addgene # 231416 | This work |
| pSEVA47_dCas9-GFP_NOT_iA | $Spc^R$ | PAJ285<br>Addgene # 231417 | Gift from the Jaramillo lab. |
| pSEVA47_dCas9-GFP_NOT_iC | $Spc^R$ | PAJ290<br>Addgene # 231418 | Gift from the Jaramillo lab. |
| pSEVA47_dCas9-GFP_YES_i2 | $Spc^R$ | Addgene # 231419 | This work |
| pSEVA47_dCas9-GFP_AAN_4 | $Spc^R$ | Addgene # 231415 | This work |

| | | | |
|---|---|---|---|
| pSEVA26_sgRNA-A0_M13ps | $Kan^R$ | Addgene # 231413 | This work |
| pSEVA19_sgRNA-C0_M13ps | $Amp^R$ | PAJ156<br>Addgene # 231410 | Gift from the Jaramillo lab. |
| pSEVA63_sgRNA-B0_M13ps | $Gent^R$ | PAJ552<br>Addgene # 231421 | Gift from the Jaramillo lab. |

#### Table S3: Core Strains

| Core Strain | Genotype | Plasmid | Antibiotic Phenotype | Source | Used in Figure/s |
|---|---|---|---|---|---|
| TOP10 | K-12 *F*⁻ *mcr*A Δ(*mrr-hsd*RMS-*mcr*BC) φ80*lac*ZΔM15 Δ*lac*X74 *rec*A1 *ara*D139 Δ(*ara-leu*)7697 *gal*U *gal*K λ⁻ *rps*L(Str$^R$) *end*A1 *nup*G | - | Str$^R$ | Gift from the Jaramillo lab.[31] | Fig. 1, 3, 4, 5, 6 |
| TOP10F | *mcr*A Δ(*mrr-hsd*RMS-*mcr*BC) φ80*lac*ZΔM15 Δ*lac*X74 *rec*A1 *ara*D139 Δ(*ara-leu*)7697 *gal*U *gal*K *rps*L(StrR) *end*A1 *nup*G | F' [*lac*Iq, *Tn*10(TetR)] | Str$^{R\#}$/ Tet$^R$ | Invitrogen™ C303003 | Fig. 3, 4, 5, 6 |
| ER2738F | *fhuA2 glnV Δ(lac-proAB) thi-1 Δ(hsdS-mcrB)5* | F' [*proA*$^+$*B*$^+$ *lacI*$^q$ *Δ(lacZ)M15 zzf::Tn*10] | Tet$^R$ | New England Biolabs #E4104S | Fig. 1, S3.1a |
